# Molecular surface mimicry enables CRBN to target G3BP2 for degradation

**DOI:** 10.1101/2025.04.30.651496

**Authors:** Stefano Annunziato, Chao Quan, Etienne J. Donckele, Ilaria Lamberto, Richard D. Bunker, Mary Zlotosch, Laura Schwander, Anastasia Murthy, Lars Wiedmer, Camille Staehly, Michelle Matysik, Samuel Gilberto, Despina Kapsitidou, Daric Wible, Gian Marco de Donatis, Peter Trenh, Rohitha SriRamaratnam, Vaik Strande, Reinaldo Almeida, Elena Dolgikh, Bradley DeMarco, Jennifer Tsai, Amine Sadok, Vladislav Zarayskiy, Magnus Walter, Ralph Tiedt, Kevin J. Lumb, Debora Bonenfant, Bernhard Fasching, John C. Castle, Sharon A. Townson, Pablo Gainza, Georg Petzold

**Author notes:** These authors contributed equally to this work (Stefano Annunziato, Chao Quan, Etienne J. Donckele).

## Abstract

Molecular glue degraders (MGDs) are small molecule compounds that repurpose the ubiquitin-proteasome system to induce degradation of challenging therapeutic targets. Clinically effective MGDs bind cereblon (CRBN), a substrate receptor of the Cullin-4/RING E3 ubiquitin ligase (CRL4^CRBN^), and impose a gain-of-function activity to recruit, ubiquitinate and degrade so-called neosubstrate proteins. Known neosubstrates bind the CRBN/MGD neosurface on the CRBN CULT domain through a structural G-loop recognition motif that mimics contacts of a natural CRBN degron. Here we report the binding mode of G3BP2, a CRBN neosubstrate that bypasses the G-loop requirement by engaging an unconventional binding site on the CRBN LON domain. The MGD-induced ternary complex interface does not resemble known protein-protein interactions (PPI) with CRBN. Instead, CRBN mimics an endogenous binding partner of G3BP2 and repurposes a preexisting PPI hotspot on the target protein. Our findings provide a novel generalizable concept for the rationalization of unconventional neosubstrate binding modes on CRBN, demonstrate unprecedented potential for the reprogrammability of this substrate receptor by MGDs, and offer opportunities for rational expansion of the target repertoire accessible to this modality.

## Main text

The CRL4^CRBN^ E3 ubiquitin ligase binds molecular glue degrader (MGD) compounds that facilitate recruitment and degradation of neosubstrate proteins^1,2^. Neosubstrates engage in MGD-induced protein-protein interactions (PPIs) with an extended surface area on the CRBN substrate receptor with a high predicted propensity to form PPIs^3^. This PPI hotspot spans two flexibly attached CRBN domains, the N-terminal LON domain and C-terminal CULT domain, which undergo major conformational rearrangements upon MGD and neosubstrate binding^3,4^. Known CRBN neosubstrates share a structural recognition motif called the G-loop, which creates surface complementarity with a primary neosubstrate binding site in the CULT domain that is proximal to the MGD-binding pocket^5–7^. G-loops represent a simple secondary-structure element that shows high prevalence in the human proteome^8,9^, enabling computational exploration of the CRBN target space.

We previously established a proteome-wide map of the human CRBN target space through structure-based predictions of surface-exposed G-loop-like motifs^3^. This resource not only facilitates identification of G-loop-containing neosubstrates, but can also guide the prospective discovery of G-loop-independent targets in unbiased proteomic screens. In mass spectrometry (MS)-based global proteomics, we found that 6 h treatment of CAL51 cells with the dihydrouracil-based MGD, MRT-5702 (**Fig. 1a**), potently reduced levels of Ras–GAP SH3 domain binding protein 2 (G3BP2; **Fig. 1b**), and to a lesser extent ubiquitin-specific peptidase 10 (USP10)^10^. Both proteins lack predicted G-loop-like motifs^11^. Reduction of G3BP2 and USP10 protein levels by MRT-5702 was dependent on cullin neddylation, proteasome activity, and the presence of the CRBN substrate receptor of the CRL4^CRBN^ E3 ubiquitin ligase, as could be demonstrated by pathway inhibitors (**Fig. 1c**) and a genetic *CRBN* knockout (**Extended Data Fig. 1a**). In a cellular NanoBRET system, MRT-5702 produced a dose-dependent bioluminescent resonance energy transfer (BRET) signal in HEK293T cells ectopically expressing NanoLuc-G3BP2, but not in cells expressing NanoLuc-USP10 or NanoLuc-G3BP1 (**Fig. 1d**). These data show that G3BP2 is a *bona fide* CRBN neosubstrate whose recruitment and degradation is induced by MRT-5702.

**Fig. 1.**
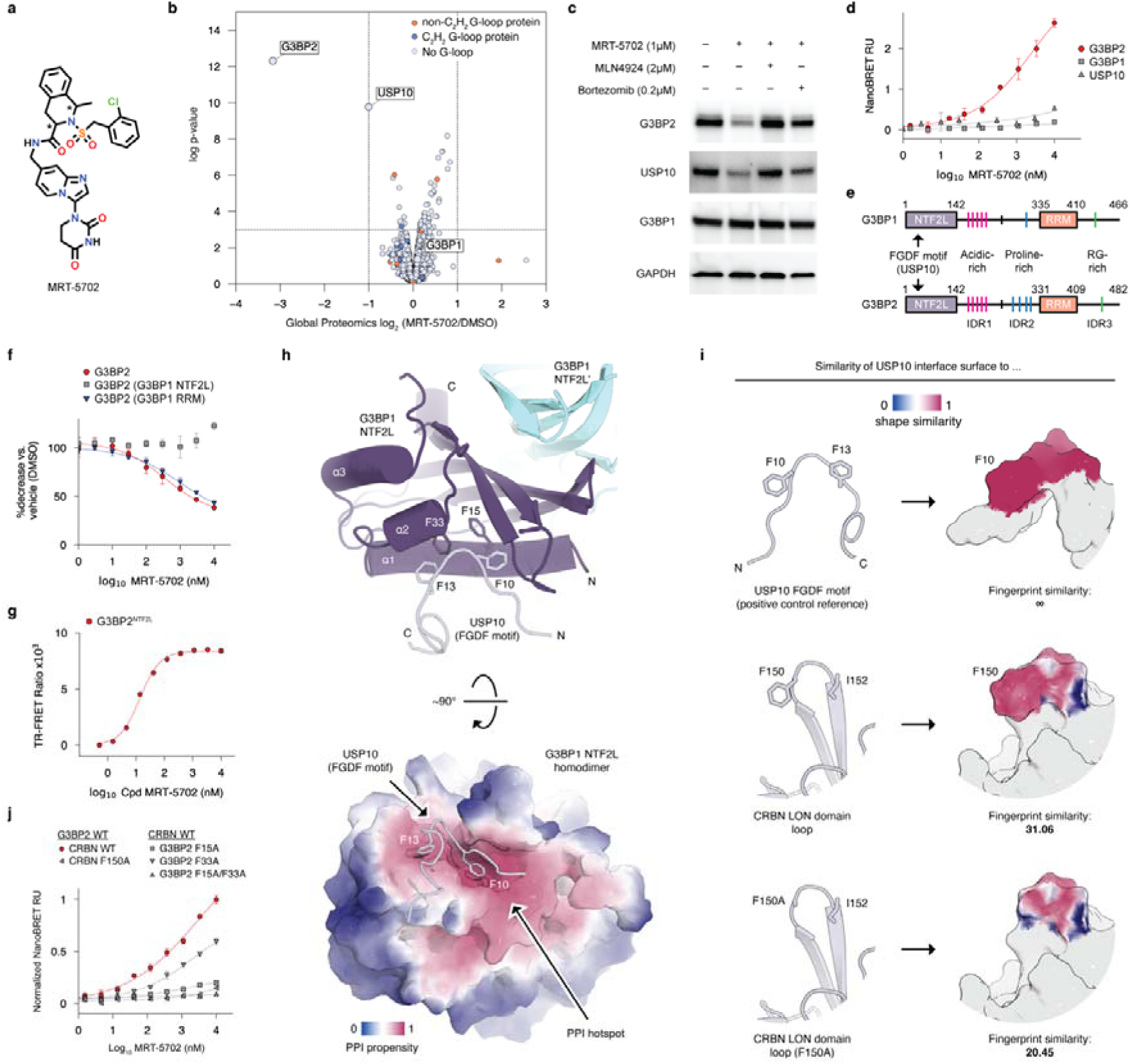
G3BP2 is a G-loop-independent CRBN neosubstrate. **a,** Chemical structure of MRT-5702. Asterisks indicate chiral centers in the compound tail. **b,** Global TMT proteomics in CAL51 cells using MRT-5702 (10 μM, 6 h). Proteins with predicted β-hairpin G-loops in C2H2 and non-C2H2 domains are labeled. **c**, Western blot in CAL51 cells using MRT-5702 (1 μM, 24 h) and rescue of degradation by co-treatment with pathway inhibitors. **d**, Ternary complex formation between CRBN and G3BP2, G3BP1 and USP10, measured by NanoBRET. **e**, Domain architecture of G3BP proteins. **f**. Levels of overexpressed HiBiT-G3BP2 variants in CAL51 cells measured by luminescence after 24 h compound treatment. **g**, Ternary complex formation between recombinantly purified DDB1/CRBN and the NTF2L domain of G3BP2, measured by *in vitro* TR-FRET. **h**. Structure of the G3BP1 NTF2L homodimer bound to the FGDF-motif of USP10 (PDB: 7xhf). Predicted PPI propensity plotted onto the surface of the NTF2L domain shown at the bottom. **i**, Surface similarity between the USP10 interface with G3BP proteins, a surface patch in the CRBN LON domain, and the CRBN F150A mutant. USP10 self-comparison serves as a positive control reference. Fingerprint score is a combined score of shape similarity (shown in surface plots) and electrostatics. Details about surface alignments can be found in the material and methods section. **j**, Ternary complex formation between wild-type and interface mutant forms of CRBN and G3BP2, measured by NanoBRET.

G3BP2 is implicated in RNA processing and stress granule formation^12^. Aberrant G3BP2 expression occurs in multiple human malignancies^13–15^ and G3BP2 stress granule formation is linked to cardiovascular and neurodegenerative disorders^16^. G3BP2 and its paralog, G3BP1, show a high degree of sequence conservation^17^ (64% identity, 76% similarity; **Extended Data Fig. 1b**) and a comparable domain architecture (**Fig. 1e**), suggesting functional redundancy of both G3BP proteins^18^. As G3BP1 levels appear unaffected by MRT-5702 (**Fig. 1b, c**), we performed domain-swap experiments to locate the site on G3BP2 that is required to associate with CRBN. Replacement of the N-terminal nuclear transport factor 2-like (NTF2L) domain, but not of the C-terminal RNA recognition motif (RRM) domain in G3BP2 with the corresponding G3BP1 sequences, abrogated MRT-5702-mediated degradation of the chimeric constructs (**Fig. 1f**). Conversely, MRT-5702-mediated G3BP1 degradation could be engineered by the exchange of its NTF2L domain with that of G3BP2 (**Extended Data Fig. 1c**). As the recombinant NTF2L domain of G3BP2 (G3BP2^NTF2L^; 1-139aa) forms a ternary complex with CRBN and MRT-5702 in TR-FRET *in vitro* (**Fig. 1g**), our data show that the NTF2L domain is necessary and sufficient for MGD-induced recruitment and degradation of G3BP2 by the CRL4^CRBN^ ubiquitin ligase.

As computational analyses of the G3BP2 NTF2L domain did not identify structural elements with similarities to β-hairpin or helical G-loop degrons^3^, we explored other strategies to pinpoint the CRBN binding site on G3BP2. Recently, the G-loop-independent target VAV1 was shown to engage CRBN through surface features that resemble the GSPT1 degron, a well-established CRBN neosubstrate, demonstrating that target compatibility with CRBN depends on surface complementarity rather than the presence of conserved structural motifs^3^. Examination of the G3BP2 NTF2L domain for surface similarities to known CRBN degrons, however, only identified low-scoring surface patches lacking key features of characterized CRBN neosubstrates. This suggests that G3BP2 forms a novel PPI interface with CRBN by engaging in an unprecedented binding mode. Protein surfaces are classified based on their predicted propensity to engage in PPIs^19,20^, and regions of high PPI propensity are frequently utilized by various interaction partners^21–24^. As no degron mimicry could be detected between G3BP2 and known CRBN neosubstrates, we took the inverse approach and examined CRBN for surface similarities to natural interaction partners of G3BP2. Both G3BP paralogs interact with a variety of different proteins, including the known interaction partner USP10^10^, through recruitment of FGDF or FxFG primary sequence motifs to their respective NTF2L domains (**Fig. 1h** and **Extended Data Fig. 2a, b**). These motifs bind a conserved site on the NTF2L domain that is predicted to possess a high propensity to engage in PPIs^25–27^ (**Fig. 1h**). This PPI hotspot is readily accessible on both protomers of the constitutive NTF2L homodimer (**Extended Data Fig. 2c**) and could potentially be deployed by CRBN despite the absence of obvious FGDF or FxFG motifs.

To rationalize the interaction between CRBN and G3BP2, we used the structure of the G3BP1-USP10 complex^26^ (**Fig. 1h**) and generated a model of the USP10-engaged NTF2L domain of the G3BP2 paralog (**Extended Data Fig. 2d**). This model was used as a query to computationally probe the CRBN surface for molecular mimicry to the USP10 interface. We identified a surface patch on the N-terminal LON domain of CRBN centered around residues F150, G151, and I152 that mimics features of the FGDF motif (**Fig. 1i**). Using the predicted CRBN surface patch to guide the creation of a ternary complex docking model (see Methods) resulted in an overall compatible orientation of the G3BP2 NTF2L homodimer with the closed conformation of CRBN^4^ (**Extended Data Fig. 2e and f**). Although engaged with the LON domain, the model positioned the NTF2L homodimer proximal to the MGD-binding pocket located in the neighboring CULT domain of CRBN, which in principle could allow for interactions with MRT-5702 (**Extended Data Fig. 2f**). To probe the interface prediction experimentally, we mutated key residues in CRBN and G3BP2 and tested ternary complex formation of the mutant constructs in NanoBRET. Mutation of CRBN F150 to alanine ablated binding of full-length wild-type G3BP2 (**Fig. 1j**). Similarly, mutations in the PPI hotspot of G3BP2 affected ternary complex formation with CRBN (**Fig. 1j**). These data support that, in the presence of MRT-5702, CRBN uses a molecular surface mimicry mechanism to engage the NTF2L domain by repurposing a preexisting PPI hotspot on G3BP2.

To validate the ternary complex interface at a molecular level, we used single-particle cryo-electron microscopy (cryo-EM) to determine the structure of an all-human complex composed of DDB1^ΔBPB^-CRBN^Δ^^1–40^, MRT-5702, and the G3BP2 NTF2L homodimer (**Fig. 2a**). The 3D reconstruction was refined to a resolution of 2.5 Å, with a local resolution of the CRBN-G3BP2 interface ranging from 3.5 to 4.5 Å (**Extended Data Fig. 3 and 4**). Local refinements using soft masks revealed noticeable conformational flexibility of the ternary complex interface but confirmed occupancy of MRT-5702 (**Extended Data Fig. 4i**). MRT-5702 carries two chiral centres in the compound tail^28^ (**Fig. 1a**), which contacts G3BP2 in the cryo-EM model (**Extended Data Fig. 4i**).

**Fig. 2.**
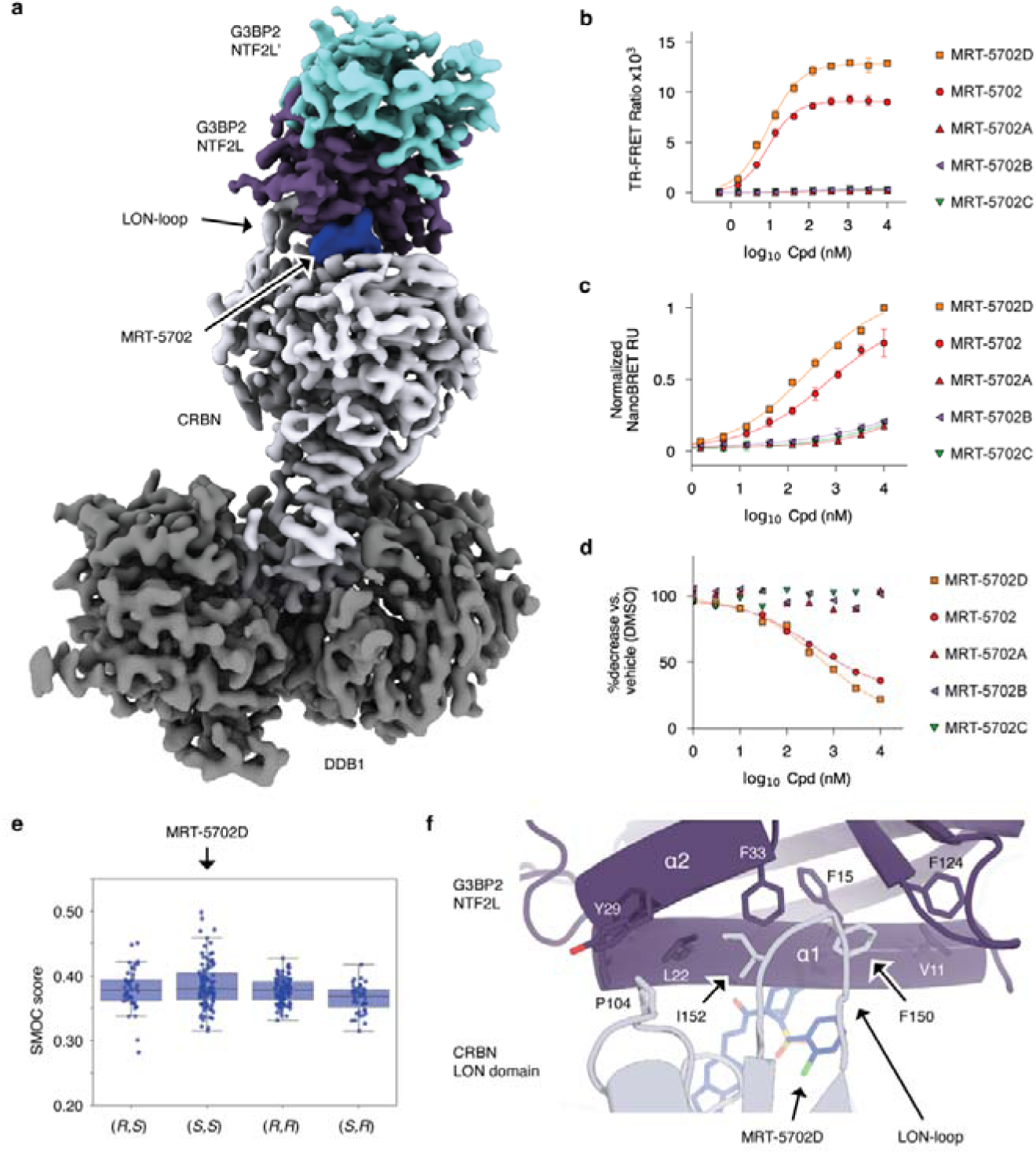
Cryo-EM structure of the DDB1/CRBN-MRT-5702-G3BP2 ternary complex. **a**, Composite cryo-EM map of the DDB1^ΔBPB^/CRBN-MRT-5702-G3BP2^NTF2L^ ternary complex. **b**, *In vitro* TR-FRET between DDB1/CRBN and G3BP2^NTF2L^ in the presence of racemic MRT-5702 or its separated enantiomers. **c**, Cellular NanoBRET between CRBN and full-length G3BP2 in the presence of racemic MRT-5702 or its separated enantiomers. **d**, Levels of overexpressed HiBiT-G3BP2 in CAL51 cells measured by luminescence after 24h compound treatment. **e**, Volume overlap of calculated low energy conformation of the different stereoisomers with the ligand density in the cryo-EM envelope (SMOC: segment-based Manders overlap coefficient). **f**, Close-up of the PPI interface between G3BP2^NTF2L^ and the CRBN LON-loop in the derived cryo-EM model.

To distinguish G3BP2-compatible configurations of MRT-5702, we separated the two pairs of enantiomers (denoted A-D) using supercritical fluid chromatography (**Extended Data Fig. 5 and 6**). The resulting compounds were profiled for their ability to induce ternary complex formation with CRBN and G3BP2 degradation in cells (**Fig. 2b-d**), which revealed MRT-5702D as the only configuration with G3BP2 activity. To assign the correct ligand configuration to MRT-5702D, we calculated ensembles of low energy ligand conformations for both pairs of enantiomers, modelled them into the MGD binding pocket of CRBN and removed all ligand conformations that caused major clashes with the surrounding protein chains (see Methods). The remaining conformers were scored according to their overlap with the ligand density in the cryo-EM envelope (**Fig. 2e**), which points to the (*S,S*)-configuration for the G3BP2-active compound MRT-5702D (**Extended Data Fig. 7**). This assignment is further supported by small molecule crystallography of MRT-5702A and MRT-5702C (see Methods), which could be attributed to the (*R,S*)- and (*S,R*)-configuration, respectively.

The cryo-EM model validated the predicted interface between the PPI hotspot of G3BP2 and CRBN residues F150 to I152 (**Fig. 2f** and **Extended Data Fig. 8a and b**). Both, F150 and I152, are part of an extended loop structure in the LON domain of CRBN that contributes approximately 50% of the overall protein-protein interface with the NTF2L domain of G3BP2 (**Fig. 3a** and **Extended Data Fig. 8c-e**). This CRBN loop (LON-loop; E146-I154) is largely solvent-exposed and exhibits discernible flexibility in other CRBN structures (**Fig. 3b**), which facilitates accommodation of the G3BP2 neosubstrate through small conformational adjustments that enhance the overall compatibility of the NTF2L homodimer with CRBN and MRT-5702D (**Fig. 3c** and **Extended Data Fig. 8b-e**). These rearrangements create an extended interface between G3BP2 and CRBN that provides opportunities for the proximal NTF2L protomer to engage in additional interactions with MRT-5702D (**Extended Data Fig. 8e**), the CRBN sensor loop (**Extended Data Fig. 8c**) and nearby residues (**Extended Data Fig. 8d**).

**Fig. 3.**
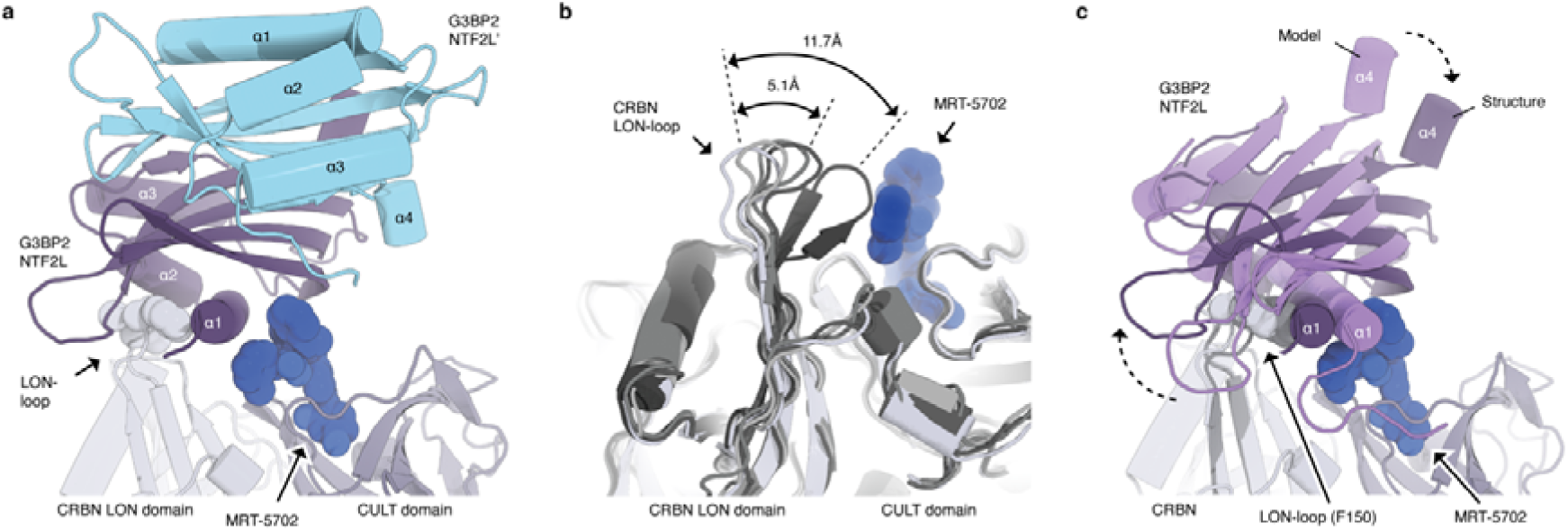
Conformational flexibility of the CRBN LON-loop enhances complementarity with G3BP2. **a**, Side view of the ternary complex interface between G3BP2^NTF2L^, MRT-5702 and the CRBN LON domain. The CRBN LON domain is shown in light grey, the CRBN CULT in dark grey. **b**, Superposition of G3BP2-engaged CRBN (light grey) with other CRBN structures (PDB entries 4tz4, 6h0g, 5v3o, 6h0f) highlights intrinsic flexibility of the CRBN LON-loop. **c**, Superposition of the G3BP2-CRBN docking model (generated with CRBN chain B of PDB entry 5fqd) with the cryo-EM-derived structural model. CRBN residue F150 shown as spheres. Differences in LON-loop conformation and overall pose of the interacting G3BP2^NTF2L^ protomer are highlighted.

Strikingly, G3BP2 lacks key contacts with CRBN residues commonly involved in neosubstrate recruitment^3,5–7^ (**Fig. 4a**), which distinguishes the binding mode of G3BP2 from other structurally characterized CRBN targets (**Fig. 4b and c**). As only one NTF2L protomer is engaged by the composite interface of CRBN and MRT-5702D, the PPI hotspot of the non-interacting protomer (NTF2L’) is in principle available for the recruitment of other interaction partners of G3BP2 (**Fig. 3a** and **Extended Data Fig. 2c**). Our ternary complex structure therefore rationalizes the observed reduction of USP10 through collateral co-degradation of a G3BP2 interaction partner (**Fig. 1b-d**). Taken together, our complementary cellular, biochemical, computational, and structural approaches discover, predict and rationalize G-loop-independent target recruitment to CRBN. These approaches can be more generally applied to the prospective discovery of novel neosubstrate binding modes on CRBN.

**Fig. 4.**
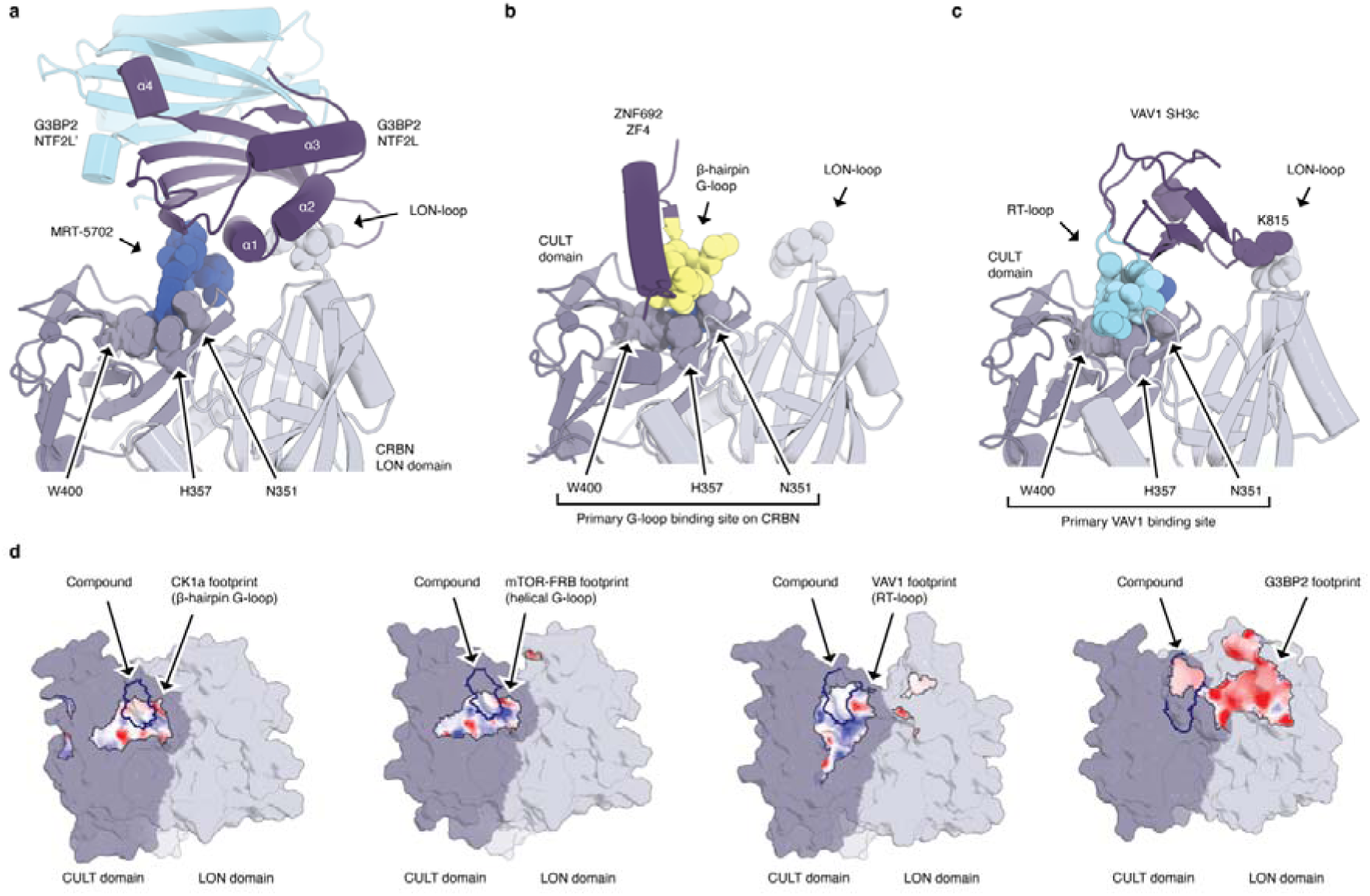
G3BP2 recruitment bypasses the canonical neosubstrate binding site on CRBN. **a**, G3BP2 ternary complex interface highlighting key contacts with MRT-5702 and the CRBN LON-loop. The CRBN LON domain is shown in light grey, the CRBN CULT domain in dark grey. **b**, Ternary complex interface between CRBN, pomalidomide and the β-hairpin G-loop target ZNF692. The primary protein-protein interface is formed between the G-loop residues (shown in yellow) of ZNF692 zinc-finger 4 (ZF4) and CRBN residues N351, H357, and W400. **c**, Ternary complex interface between CRBN, MRT-23227 and the G-loop-independent target VAV1. The primary protein-protein interface is formed between RT-loop residues (shown in cyan) of the VAV1 SH3c domain and CRBN residues N351, H357, and W400. The VAV1 nSrc-loop residue K815 forms a secondary contact with the CRBN LON-loop. **d**, Footprints of different CRBN neosubstrates mapped onto the CRBN/MGD neosurface. The MGD is indicated by a blue outline. Footprints are colored according to electrostatics.

Previously characterized CRBN neosubstrates^3,5–7,29–31^ are recruited to a primary site on the CRBN CULT domain that is composed of CRBN residues N351, H357, and W400. These three residues surround the MGD-binding pocket and form key hydrogen bond interactions with G-loops or the unconventional VAV1 RT-loop^3^ (**Fig. 4b and c**), contacts that partially mimic recognition of an endogenous CRBN degron^32–35^. G3BP2 completely bypasses this primary neosubstrate binding site and instead forms extended protein-protein interactions with the CRBN LON domain (**Fig. 4a** and **Extended Data Fig. 8**). The unconventional binding mode of G3BP2 therefore demonstrates for the first time that neosubstrate recruitment neither requires a G-loop recognition motif on the target protein nor is dependent on the primary G-loop binding site on CRBN. Contrary to canonical G-loop targets, G3BP2 does not appear to mimic naturally occurring interactions with CRBN. Instead, CRBN resembles an endogenous binding partner of G3BP2 and repurposes a PPI hotspot on the target protein. The protein-protein interface emerging from this reciprocal mimicry mechanism is further extended by the conformational flexibility of the CRBN LON-loop and complemented by direct contacts with the MGD, which together give rise to a unique CRBN/MGD neosurface that induces potent and selective degradation of G3BP2.

Protein surfaces are commonly equipped with privileged sites to engage in PPIs^19^, which can be harnessed by diverse interaction partners through the usage of conserved sequence-, structural-or surface-motifs^21–24,36^. Post-translational modifications^37^ and missense mutations^38^ frequently occur at or near such PPI sites and rewire the interactome with often binary outcomes, explaining an exquisite susceptibility of these hotspots to small molecule-mediated modulation^3,39–47^. The rationalization of the novel G3BP2 binding mode suggests that MGD-induced protein contacts between CRBN and its neosubstrates rely on at least three determinants: (i) a preexisting PPI hotspot on either side of the interface, (ii) partial molecular mimicry of an endogenous PPI and (iii) complementation of these contacts by MGD-induced neointeractions. Composite CRBN/MGD neosurfaces that mimic the footprint of natural PPIs, in short ‘*glueprints’,* provide a new conceptual framework to rationalize target engagement by CRBN and other E3 ligases, creating a roadmap for the expansion of the MGD-accessible target repertoire.

## Acknowledgements

Cryo-EM data was collected at the Cryo-EM Facility at MIT.nano, including use of the Talos Arctica gifted by the Arnold and Mabel Beckman Foundation. We thank the cryo-EM team (Sarah Sterling, Jenn Podgorski and Christopher Borsa) at MIT.nano for technical support on data collection, and Nicholas Swanson for consultation on preparing this manuscript and on depositing the structures. We thank Freya Harvey for assistance with the write-up of chemical synthesis pathways for the MRT compounds.

Author contributions:

## Conceptualization: SA, CQ, ED, JC, ST, PG, GP

Design: SA, CQ, ED, IL, AM, LW, RS, RA, BD, JT, BF, DB, JC, ST, PG, GP

Data acquisition/analysis/interpretation: SA, CQ, ED, IL, RB, MZ, LS, AM, LW, CS, MS, SG, DK, DW, GD, PT, RS, VS, PG, GP

Software: RB, PG

Supervision: SA, ED, IL, RS, RA, LD, BD, JT, AS, VZ, MW, RT, KL, BF, DB, JC, ST, PG, GP

Narrative and initial draft: SA, CQ, ED, JT, PG, GP

## Data and materials availability

All software code, intermediate files, and predictions will be deposited in zenodo.org. All software code was deposited in github repository (http://github.com/monterosatx/molecular_surface_mining). Protein Data Bank (PDB) files will be deposited in the Research Collaboratory for Structural Bioinformatics (RCSB) PDB and the cryo-EM data in the EMDB.

## Methods

### Cell Treatment for Global Proteomics Experiments

CAL-51 cells were cultured in DMEM (Gibco) supplemented with 10% FBS (Corning). For Global Proteomics profiling experiments, 12-well plates of sub confluent 70-90% CAL51 cells were incubated for 6 h at 37 °C in fresh medium DMEM high Glucose (Gibco) + 10% FBS (Corning) with 1 µM compound or 0.1% DMSO (Fisher Bioreagents). Cells were washed once with PBS and collected with 0.2 mL of TrypLE (Gibco) in 1.5 mL Eppendorf tubes protein LoBind. Cells were pelleted by centrifugation at 300xg for 4 min at 4 °C, supernatants were removed, and cells were washed with 1mL of PBS. After centrifugation, dried pellets were frozen to −80 °C.

### Global Proteomics Profiling Sample Preparation

Cell pellets were lysed using 90 µL PreOmics iST-NHS lysis buffer and sonicated using the UP200St ultrasonic processor from HUBER LAB for 10 s per sample at 35% amplitude. A tryptophan assay was used to determine lysate concentration. Plates were read with a Tecan Infinite 200 PRO and lysates normalized to 100 μg in 50 μL. The normalized lysates were then processed using the recommended PreOmics sample preparation protocol with minor modifications. In brief, proteins were reduced, alkylated, and digested for 3 h at 37 °C. The peptides were then labelled with tandem mass tag (TMT) reagent (1:4; peptide:TMT label) for 1 h at 22°C. Labeled peptides were then quenched with 5% hydroxylamine solution diluted in LC/MS-grade water. The peptides from the 16 conditions were then combined in a 1:1 ratio and purified with PreOmics desalting cartridges. Mixed and labeled peptides were subjected to high-pH reversed-phase HPLC fractionation on an Agilent X-bridge C18 column (3.5 µm particles, 2.1 mm internal diameter, and 15 cm in length). Using an Agilent 1200 LC system, a 66 min linear gradient from 10% to 40% acetonitrile in 10 mM ammonium formate, pH 10, separated the peptide mixture into a total of 60 fractions, which were then consolidated into 24 fractions containing an assortment of TMT-labeled fragments in each. Peptides concentrations were estimated by UV and after drying in a SpeedVac integrated vacuum concentrator, all 24 fractions from the plate were resuspended to 0.2 µg/uL with 0.1% formic acid based on the peptide UV chromatogram. Each of the fraction was loaded onto a 25 cm Aurora column (75 µm internal diameter, 1.6 µm particles) from Ion Optics using a Vanquish Neo HPLC system. The peptides were separated using a 168 min gradient from 4% to 30% buffer B (80% acetonitrile in 0.1% formic acid) equilibrated with buffer A (0.1% formic acid) at a flow rate of 400 nL/min. Eluted TMT peptides were analyzed using an Orbitrap Eclipse mass spectrometer.

### Data Acquisition and Analysis

MS1 scans were acquired at resolution 120,000 with 400-1400 mass over charge (m/z) scan range, Automatic Gain Control (AGC) target 4 x 10^5^, maximum injection time 50 ms. Then, MS2 precursors were isolated using the quadrupole (0.7 m/z window) with AGC 1 x 10^4^ and maximum injection time 50 ms. Precursors were fragmented by collision-induced dissociation (CID) at a normalized collision energy (NCE) of 35% and analyzed in the ion trap. Following MS2, synchronous precursor selection (SPS) MS3 scans were collected by using high energy collision-induced dissociation (HCD) and fragments were analyzed using the Orbitrap (NCE 55 %, AGC target 1 x 10^5^, maximum injection time 120 ms, resolution 60,000). Protein identification and quantification were performed using Proteome Discoverer 2.5.0.400 with the SEQUEST algorithm and UniProt human database (2019, 20602 protein sequences). Mass tolerance was set at 10 ppm for precursors and at 0.6 Da for fragments. A maximum of 2 missed cleavages were allowed. Methionine oxidation was set as dynamic modification, while TMT tags on peptide N termini/lysine residues and cysteine alkylation (+113.084) were set as static modifications. Adjustment of reporter ion intensities for isotopic impurities according to the manufacturer’s instructions was performed in Proteome Discoverer. Subsequently, protein identifications are inferred from unique peptides, i.e. peptides matching multiple protein entries are excluded. Protein relative quantification is performed using an in-house developed software in Python and R. This analysis included multiple steps; de-duplication of PSMs to peptide charge modification (PCM) level using the average, the sum and the maximum abundance of TMT channels to sort and pick the best PCM, addition of 1 to abundances of PCMs (for all channels) if any of the abundance values is below 0 (for subsequent log2 transformation). MSstatsTMT^48,49^ is used for global normalization of TMT channels (equalize medians), to normalize PCMs (for all channels) and summarize protein abundances using Tukey’s Median Polish, missing values are imputed using maxQuantileforCensored=NULL. As an intermediate step, before protein summarization, a custom algorithm is applied to find outliers of imputed values (cases where the model fails), and all PCMs for the given protein are reset to being missing. An arbitrary L2FC cut-off of −1 and +1 is applied to the dataset corresponding to a minimal −50% depletion (x0.5) or +100% enrichment (x2) respectively. The p-value cut-off for statistical significance is set at the standard 10-2 value.

### Cellular NanoBRET

The mammalian gene expression vector for transient transfection of HaloTag®-CRBN was purchased from Promega. NanoLuciferase (NanoLuc)-fusion vectors were purchased from VectorBuilder and were designed consisting of a N-terminal NanoLuc fused to a human target gene with a 3xGGGGS linker. HEK293 cells were cultured in Dulbecco’s modified Eagle’s medium (DMEM) (Gibco) +10% fetal bovine serum (FBS) (Gibco). Medium was removed from flasks by aspiration, cells were washed with phosphate buffered saline (PBS), and trypsin-EDTA was added to dissociate cells. Culture medium was added to the cells to neutralize trypsin, then cells were transferred in a centrifuge tube and centrifuged at 300×g for 3 min. Media was aspirated from the cell pellet and cells were resuspended in culture medium and passed through a 40 μm cell strainer. Cells were counted using a Cellometer® cell counter and then diluted to 4×10^5^ cells/mL in full media. Cells were seeded in 6-well plates, 2 mL/well for a total 8×10^5^ cells/well. Cells were incubated overnight at 37°C, 5% CO_2_. On day two, cells were transfected with a Nanoluc (Donor): HaloTag (Acceptor) DNA co-transfection mix using Fugene HD (Promega). HaloTag-CRBN and Target-NanoLuc fusion vectors were added to 100 μL phenol red-free OptiMEM (Gibco). Finally, 6 μL FuGENE HD was added to the transfection mix, and the mix was gently agitated. After incubation for 10-15 min at room temperature, each transfection mix was added dropwise to one well of the overnight-cultured cells.

Multiple wells were transfected for each target to perform the experiment. Cells were incubated overnight at 37°C, 5% CO_2_. After 18-20 hours, media was aspirated, cells were washed with PBS, and then trypsin was added. Dissociated cells were collected using phenol red-free OptiMEM +4% FBS, placed in a centrifuge tube, and centrifuged at 300×g for 3 minutes. Media was then aspirated from the pellet, and the pellet was resuspended in phenol red-free OptiMEM + 4% FBS. Cells were counted as previously described and diluted in phenol red-free OptiMEM +4% FBS to 1.5×10^5^ cells/mL. HaloTag 618 Ligand (Promega) was added to the cells. Cells treated with DMSO and without 618 ligand were used to calculate background. Cells (25 μL/well) were seeded in a white, solid bottom 384-well plate (Corning). Plates were then incubated overnight at 37°C and 5% CO_2_. On day four, compounds were added to the plates in 10-point, 1:3 dilution series starting at 10 μM. After treatment, plates were returned to the incubator for 3 h. After 3 h, NanoBRET Nano-Glo Substrate (Promega) was diluted in Phenol red-free OptiMEM +4% FBS and added to the plates using a multidrop Combi. Dual luminescence measurements were then taken on a PHERAstar® FSX using the LUMI 610 LP 450-80 H optic module. The ratio between signal recorded in the acceptor channel and the donor channel was used to determine the Bioluminescence Resonance Energy Transfer (BRET) at a well-to-well level. Average signal from wells containing no 618 ligand was used to subtract background and curves were normalized to the average DMSO treated wells containing 618 ligand. NanoBRET curves were further normalized, such that the values were rescaled to a range between 0 and 1 by subtracting the minimum and dividing by the range of the data (maximum – minimum). Correspondingly, the standard deviations were normalized by dividing by the same range factor.

### Western Blot

CAL51 cells were plated on a 24-well plate at a density of 0.4 x10^6^ cells/well, incubated overnight and treated with 1 µM MRT-5702 or DMSO. Controls including co-treatment of Bortezomib (0.2 μM) or MLN4924 (2 μM) were also performed. Cells were collected and lysed 30 min with RIPA lysis buffer (ThermoFisher) supplemented with protease inhibitor cocktail (Sigma), Phosphatase Inhibitor solution II (Sigma), and Phosphatase Inhibitor solution III (Sigma). Samples were analyzed by western blot using the following primary antibodies, incubated overnight at 4°C, diluted 1/1000: G3BP1 (Proteintech, Cat. No. 13057-2-AP), G3BP2 (Proteintech, Cat. No. 16276-1-AP) or USP10 (CST, Cat. No. 19374-1-AP).

Membranes were incubated for 2 h at room temperature with a secondary antibody, anti-rabbit HRP antibody diluted 1/5000 (Invitrogen, Cat. No. A16110), and GAPDH typically used as a loading control (Invitrogen, Cat. No. MA5-15738-A647). Membranes were developed on ChemiDoc imaging system (Biorad) using ECL (enhanced chemiluminescence) substrates (Biorad).

### HiBiT assay

CAL51 HiBiT-G3BP1 and G3BP2 cell lines were generated via stable transduction with a lentiviral vector carrying a constitutive expression cassette. For the experiment, cells were plated in 96-well white flat bottom plates (Corning) at 10,000 cells per well using Multiflo (BioTek/Agilent) and in 100 µl volume in DMEM experimental medium: DMEM (DMEM, high glucose, HEPES, no phenol red (ThermoFisher Scientific) supplemented with 10% FBS (Corning) and 1% penicillin/streptomycin (ThermoFisher Scientific). Cells were incubated for 16 h at 37°C, 5% CO2 before compound addition. Cells were then incubated at 37 °C, 5% CO_2_ for 24 h and then quantification of HiBiT-tagged protein abundance was performed using the NanoGlo HiBiT lytic detection system (Promega) according to manufacturer’s instructions. Signal was read on a Pherastar FSX.

### Protein expression & purification

CRBN/DDB1 expression: His-TEV-CRBN^41–422^ and untagged full-length DDB1 or Strep-TEV tagged CRBN^41–442^ and untagged DDB1^ΔBPB^ (amino acids 1-395-GNGNSG-706-1140) were cloned into pFastBac1 vectors. Both recombinant baculovirus stocks were prepared using Bac-to-Bac Baculovirus Expression System (ThermoFisher) and were used to co-infect Sf9 insect cells for co-expression of both proteins. Harvested insect cells were resuspended in lysis buffer containing 100 mM Tris-HCl pH 8.0, 150 mM NaCl, 1 mM TCEP, 5% glycerol, 1 mM PMSF,1 tablet cOmplete EDTA-free protein inhibitor (Roche) and 250 U/L Bezonase and homogenized. The cell lysate was cleared by centrifuging at 23,000 g at 4°Cfor 90 min.

His-CRBN/DDB1: The clarified supernatant was incubated with Ni-NTA resin at 4°C overnight, washed and eluted with a buffer containing 50 mM Tris-HCl pH 8.0, 200 mM NaCl, 1 mM TCEP, 5% glycerol and 300 mM imidazole. The eluate was diluted 10x in 50 mM HEPES pH 7.0, 1 mM TCEP, 5% glycerol and loaded onto a Q-HP column and eluted using a gradient of NaCl up to 1M. The eluate was loaded onto a Superdex 200 16/600 pg column equilibrated with 10 mM HEPES, pH 7.0, 150 mM NaCl and 1 mM TCEP. Samples were concentrated to 30 mg/mL, aliquoted, flash-frozen, and stored at −80°C.

Untagged CRBN/DDB1ΔBPB: The clarified supernatant was filtrated through 0.22 μm membrane filter and loaded onto a Strep-Tactin column pre-equilibrated with the binding buffer containing 50 mM Tris-HCl pH 8.0, 150 mM NaCl and 1 mM EDTA. The column was washed and the proteins were eluted with a buffer containing 50 mM Tris-HCl, pH 8.0, 150 mM NaCl, 1 mM EDTA and 50 mM D-Biotin. The eluates were desalted into a buffer supplemented with 10% v/v glycerol and cleaved with TEV protease at 4 °C overnight. The TEV reaction was loaded onto a Strep-Tactin column to remove the strep-tag and the uncleaved fraction, followed by dilution into 5 times the volume of buffer containing 50 mM MES pH 6.5, 5% glycerol and 1 mM TCEP. The diluted protein was loaded onto a Q-HP column and eluted using a gradient of NaCl up to 1M. The eluate was loaded onto a Superdex 200 16/600 pg column equilibrated with 10 mM HEPES, pH 7.4, 150 mM NaCl and 1 mM TCEP. Samples were concentrated to 30 mg/mL, aliquoted, flash-frozen, and stored at −80° C.

G3BP2: GST-TEV-G3BP2^NTF2L^ (amino acids 1-139) was cloned into pET21b and expressed in BL21CodonPlus(DE3) cells. Protein expression was induced in 2YT media using 0.5 mM IPTG at 16°C for 18 hrs. The cells were lysed by homogenization in 50 mM HEPES pH 7.5, 1M NaCl, 1 mM TCEP, 10% glycerol, and the lysate was clarified by centrifugation at 23000 xg at 4°C from 90 min. The clarified lysate was incubated with GST resin overnight at 4°C, washed and eluted with 50 mM HEPES pH 7.5, 200 mM NaCl, 1 mM TCEP, 10% glycerol and 10 mM GSH. The eluate was applied to a Superdex 200 16/600 column equilibrated with 20 mM HEPES pH 7.5, 200 mM NaCl, 1 mM TCEP. Fractions containing GST-G3BP2^NTF2L^ were pooled, spin-concentrated and flash frozen with liquid nitrogen for storage at −80°C. For untagged G3BP2^NTF2L^, after GST purification, the eluate was incubated with GST-tagged TEV protease at 4°C overnight and incubated 2 h at 4°C with 10ml of pre-equilibrated GST resin. The resin was loaded onto a gravity-flow column, the flow through was collected and diluted 10x in 50 mM HEPES pH 7.5, 1 mM TCEP, 10% glycerol before it was applied to Q-HP column (Cytiva, USA). The flow-through of the anion exchange was applied to a Superdex 200 16/600 column equilibrated with 20 mM HEPES pH 7.5, 200 mM NaCl, 1 mM TCEP. Fractions containing untagged G3BP2^NTF2L^ were pooled, spin-concentrated and flash frozen with liquid nitrogen for storage at −80°C.

### TR-FRET assay

Solutions containing 12.5 nM His-CRBN/DDB1, 25 nM GST-G3BP2^NTF2L^, 1x anti-GST-d2, and 1x anti-His-Tb Cryptate Gold were prepared in 20 mM HEPES, 20 mM NaCl, 0.2 mM TCEP, 0.2 mM EDTA, and 0.005% surfactant P-20 and transferred into 384-well Proxiplates (Revvity). Compounds were dispensed into assay wells at the appropriate concentration using an Echo 650 (Beckman). The samples were incubated for 45 min at room temperature. The TR-FRET signal was measured with a Pherastar FSX plate reader (BMG Labtech) using the HTRF red emission filter (10000 x 337 nm/620 nm/665 nm).

### Computational pipeline

#### Surface computation and visualization

All the surface methods apply in this paper are minimally modified from those presented before^3^, and all the computer code is the same presented therein and publicly available. The MaSIF program^20,50^ was used to compute all surfaces used in the paper, including electrostatics. MaSIF uses the PyMesh library^51^ to process surface meshes, and Adaptive Poisson-Boltzmann Solver (APBS)^52^ to compute electrostatics.

*1. Definition of buried surface areas*

We used the definition of buried surfaces (i.e. the interface of a PPI) identical to that presented elsewhere^20,50^. Under our definition, surface vertices belonging to that buried surface are those that are part of the molecular surface in the unbound structure but buried in the bound structure. PyMesh was applied to regularize the meshes at 1.0 Å resolution. The buried surface of a subunit in a complex structure was calculated by comparing surface vertices of the subunit’s surface mesh and the complex’s surface mesh. Vertices in the subunit’s mesh were labeled as ‘interface’ meshes if they were more than 2.0 Å away from any complex surface vertex.

*2. Prediction of CRBN PPI propensity*

The MaSIF-site program^20^ was used to compute a per-vertex propensity of the G3BP2 NTF2L surface (PDB: 5drv) to become buried in a PPI, as visualized in Fig. S10A. MaSIF-site uses as input the per-vertex chemical and geometrical features projected on the surface of a protein, and uses them to compute a score for each vertex shown to correspond to its propensity become buried in a protein-protein interaction^20^. MaSIF-site is built on a geometric deep learning architecture trained to predict the buried surface from a training set of thousands of proteins. The training set did not contain any G3BP2 or G3BP1 proteins. For visualization purposes, the propensity of each vertex, computed as a real value between 0 and 1, was display as a regression, from blue to white to red.

*3. Surface matching of the USP10 – CRBN similarity*

The MaSIF-seed program^20,50^ is a surface matching pipeline that uses geometric deep learning to encode surface patches as fingerprints. MaSIF-seed was designed and tested to match surfaces that are complementarity to each other based on their fingerprint, followed by an alignment step. MaSIF-seed was modified to match patches by similarity (or *mimicry*) instead of complementarity.

*4. Overall method description of MaSIF-seed*

MaSIF-seed^50^ method is a geometric deep learning approach designed to identify complementary structural motifs that can mediate protein-protein interactions. It works by analyzing protein molecular surfaces, which are divided into overlapping radial patches, to capture both chemical and geometric features relevant to binding. Every vertex on the surface becomes the center of its own radial patch of radius 12 Å. A neural network is then trained to generate “fingerprints” for these overlapping surface patches that predict their binding propensity. MaSIF-seed uses this fingerprint matching process to find complementary surface motifs from a large database of potential binding partners. The MaSIF-seed method involves an important alignment step after the initial identification of complementary surface motifs. Once the surface patches from the target and potential binding seeds are matched based on their fingerprints, these patches are aligned geometrically to evaluate how well they fit together. A second neural network, called the interface post alignment (IPA) score network is used to score the aligned surface patches specifically for complementarity.

*5. Definition of the USP10 interfaces as a surface patch.*

A structure of USP10 bound to G3BP2 NTF2L is not available, but there is one of G3BP1 NTF2L in the presence of USP10 (PDB: 7xhf). Since the USP10 interface is highly conserved in G3BP1 and G3BP2, we built a simple model of G3BP2 NTF2L bound to USP10 by aligning the G3BP2 NTF2L structure (PDB: 5drv) to the G3BP1 NTF2L in its structure with USP10 (PDB: 7xhf) and then removing G3BP1. The molecular surfaces of USP10 from its co-crystal structures with G3BP1 NTF2L was used as input to the algorithm. The buried surface in the G3BP1:USP10 complex was computed as described above. The interface patch was defined as the patch of radius 12 Å that captures the largest area of the USP10:G3BP1 region. The G3BP2 NTF2L in the model was used only for clashing computation purposes.

*6. Modification of MaSIF-seed for surface similarity*

MaSIF-seed was modified to match and align similar surface patches instead of complementary patches. MaSIF-seed by default inverts the input features and normals of the target patch to align for complementarity, but this is no longer required for similarity matching. To align patches by similarity, these features were no longer inverted. MaSIF-search fingerprints were computed for all the patches decomposed from the molecular surfaces of the CRBN closed conformation (PDB: 5fqd). In a first stage, patches with fingerprints and an Euclidean distance greater than 2.5 units in the embedding space were discarded. Patches that passed this filter were selected for a second stage alignment step. Since every vertex in the surface is the center of a patch, each vertex on the surface has a computed fingerprint. In the second stage alignment step, the RANSAC algorithm is used to perform an alignment between the patches of CRBN and those of USP10. In each iteration of RANSAC three vertices in the matched patch are randomly selected, and the most similar fingerprint in the query patch (USP10) are identified. The three matches are used to perform an alignment and the number of inlier vertices after the alignment is used to initially score the alignment. The alignment with the highest number of inliers is accepted. Finally, a scoring of similarity is performed using the scored fingerprints. Specifically, after an alignment all vertices of the aligned patch within a 3D distance of 2.0 Å from a vertex in the query patch are selected. The fingerprint distance *d* is computed between the selected vertices and each of the corresponding nearest neighbor’s fingerprint in the query patch. The score for the alignment is computed as 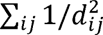 such that *i* is the vertex in the aligned patch and j is the vertex in the query patch and d is the fingerprint Euclidean distance. The interface post-alignment (IPA) scoring neural network ^50^ was set to a small value (0.50). MaSIF-seed contains a steric, atomic based post-processing filter which is used by default to filter matched results for steric compatibility with the complementary target. Since similarity was performed here, the input was modified so that the steric filter comparison is performed against G3BP2 in the model, removing any solution where no steric clashes involving Cα atoms were allowed, and up to 10 clashes involving other heavy atoms were allowed.

*7. Computation of surface similarity*

To visually display the surface similarity of CRBN and USP10 for representation in Fig. 1, after the surface-based alignment and scoring, a per-surface-vertex surface similarity was computed on the surfaces after alignment by the algorithm. The surface similarity score was computed in the same way that per-vertex shape complementarity has been previously defined ^53^. Let A and B be two proteins aligned under our surface alignment algorithm. Let P_A_ be the molecular surface of protein A, and x_A_ be a vertex on the surface of A with normal n_A_. Let P_B_ be the molecular surface of protein B, and x^’^*_A_* be the point on the surface of B closest to x_A_ with normal n^’^*_A_*.Then, the surface similarity for vertex x_A_ is defined as:

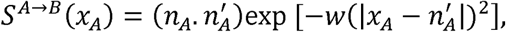

where w=0.25.

To provide a visual contrast with other surfaces and help the readers visualize surface similarity, we selected a CRBN F150A mutant as a counter example. Since using the masif-based methods on CRBN F150A did not produce a similar pose to that of wildtype CRBN, we structurally aligned CRBN F150A to the predicted CRBN pose and computed both its fingerprint similarity score (see preceding section) and its per-point surface similarity.

### Preparation of samples for cryo-electron microscopy

Purified CRBN^41–442^-DDB1^ΔBPB^ and untagged G3BP2^NTF2L^ were mixed at equimolar concentrations in a buffer containing 20 mM MES (pH 7.0), 100 mM NaCl, 1 mM TCEP and 0.05 mM Lauryldimethylamine-N-Oxide (Anatrace). The final concentrations of CRBN^41–442^-DDB1^ΔBPB^ and G3BP2 were 4.6 mg/ml and 1.1 mg/ml respectively. MRT-5702 was added to a final concentration of 100 uM. The sample was incubated on ice for 10 min before being clarified by centrifugation at 12,000 g at 4° C for 5 min. The sample was kept on ice throughout the sample preparation process. C-flat™ holey carbon 300 mesh Au R1.2/1.3 grids (Electron Microscopy Sciences) were glow discharged for 60 s at 20 mA with PELCO easiGlow^TM^ Glow Discharge Cleaning System (Ted Pella, Inc.) at room temperature in vacuum at 0.1 mbar. 3 µL of the sample were applied to each grid, blotted for 5 s with filter paper, plunged into liquid ethane in 100% humidity at 4° C using a Vitrobot Mark IV system (Thermo Fisher Scientific), and stored in liquid nitrogen.

### Cryo-EM data acquisition and analysis

Cryo-EM micrographs were collected using EPU (Thermo Fisher Scientific) on a Titan Krios G3i (Thermo Fisher Scientific) cryo-transmission electron microscope operating at 300 kV at a nominal magnification of 130,000X in super resolution mode on a Gatan K3 direct detector corresponding to effective pixel size of 0.654 Å (0.327 Å super resolution). Analysis of the micrographs was performed in CryoSPARC v4.2.1 (Structura Biotechnology) with the processing workflow displayed in Extended Data Fig. 3. In summary, the micrographs were pre-processed by motion correction and CTF estimation. Particles were picked with an elliptical blob picker with minimum diameter at 100 Å and maximum diameter at 170 Å, respectively, using the first 500 movies. The picked particles were extracted with a box size of 600 pixels. 2D classification was performed and the most promising classes were selected as template classes. Picked particles from template-based picker underwent three cycles of particle extraction, 2D classification, and 2D selection. In the first and second round, the extraction boxes were Fourier cropped from 600 pixels to 150 pixels and 300 pixels respectively, while the extraction boxes of the last round were not Fourier cropped. Selected particles from the last round of 2D classification were used to generate initial volumes *ab initio*, which were fed into three consecutive rounds of heterogenous refinement. Particles from the best volume corresponding to the DDB1^ΔBPB^/CRBN-MRT-5702-G3BP2^NTF2L^ complex were refined with non-uniform (NU) refinement, followed by global and local CTF refinement and a second round of NU refinement. The final particles were subject to particle subtraction using a soft mask covering DDB1, and the subtracted particles were used for local refinement using soft masks covering CRBN and G3BP2^NTF2L^. Final volumes were post-processed with DeepEMhancer to generate maps with improved side-chain densities to aid interpretation of the maps and manual building of the atomic model.

### Atomic model building and refinement

To generate the initial atomic model of the DDB1^ΔBPB^/CRBN-MRT-5702-G3BP2^NTF2L^complex, published models for CRBN-DDB1^ΔBPB^ (PDB: 5fqd) and G3BP2 (PDB: 5drv) were separately docked into the maps using UCSF ChimeraX^54^ version 1.4. Restraints and coordinates for MRT-5702D were generated and manually curated with elBOW and REEL in Phenix^55^ v1.21.5207. Iterative rounds of manual building and real-space refinement of the model were performed in Phenix^55^ and Coot^56^ version 0.9.8.1. In some cases, globally sharpened maps and locally refined maps and DeepEMhancer^57^ post-processed maps were used to guide manual building and visualization of the model, but real-space refinement and validation were always performed against unsharpened maps. Composite maps were generated with Phenix^55^ for the sole purpose of making figures. UCSF ChimeraX^54^ and Pymol 2.5.2 were used to visually evaluate the maps and the atomic models, as well as to generate figures. Quality of model-map agreement and geometry of the model were evaluated in Phenix^55^ and the results were listed in Extended Data Table 1.

### Calculation of low energy conformations of MRT-5702 Stereoisomers

Ligand conformation generation was carried out with OMEGA^58^ version 5.1.0 (10.1021/ci100031x) with Ewindow of 10 kcal/mol and Maxconfs of 800. The four different enantiomers of MRT-5702 were submitted as SMILES strings to OMEGA to avoid carry-over of conformation information before the calculations. MRT-5702 shares the chemical substructure of 1-(imidazo[1,2-a]pyridin-3-yl)dihydropyrimidine-2,4(1H,3H)-dione with that of MRT-3486 in the ternary complex x-ray structure between CRBN-DDB1 and NEK7 (PDB: 9nfq), which binds the tri-Trp pocket of CRBN and served as a template for ligand alignment. To this end, the position of this chemical substructure was extracted from 9nfq after superposition with the CRBN-DDB1 model derived from the cryo-EM map of MRT-5702 (aligned via CRBN residues 380, 386+ and 400). The pose of the extracted chemical substructure was energetically minimized within the CRBN pocket of the Cryo-EM model using Maestro version 14.0 to remove potential model bias from 9NFQ. The generated conformers of MRT-5702 were aligned to the minimized 1-(imidazo[1,2-a]pyridin-3-yl)dihydropyrimidine-2,4(1H,3H)-dione moiety using RDKit version 2024.03.5 (www.rdkit.org). Aligned conformers were assessed by PoseBusters^59^ for clashes with the cryo-EM model of CRBN-DDB1 or G3BP2 by calculating volume overlaps between ligands and the protein model. Ligand conformers with volume overlaps greater than 7.5% were discarded^59^. The remaining conformers of all four MRT-5702 stereoisomers were scored according to their ligand volume overlap with the experimental cryo-EM map by calculating the SMOC score (segment-based Manders overlap coeff.)^60^. The SMOC score is calculated as implemented in the python package of TEMPy^61^ version 2.0.0 on the full Cryo-EM map and plotted for each stereoisomer using seaborn^62^.

### Chemical synthesis of MRT-5702

**Scheme 1:**
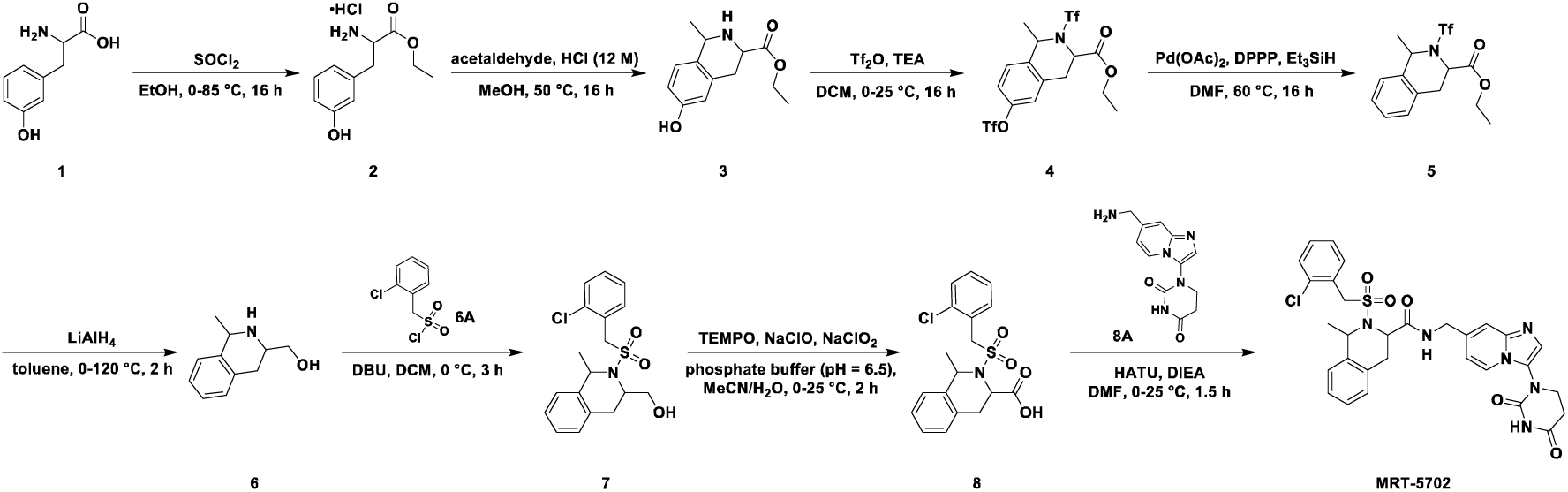
Synthesis of MRT-5702.

To a suspension of 2-amino-3-(3-hydroxyphenyl)propanoic acid (**1**) (10.0 g, 55.2 mmol, 1.00 *eq*) in ethanol (100 mL) was added dropwise sulfurous dichloride (15.1 g, 127 mmol, 9.22 mL, 2.33 *eq*) at 0 °C. The mixture was stirred at 85 °C for 16 h. The mixture was concentrated under reduced pressure to give intermediate **2** (15.0 g, crude) as yellow oil. A mixture of crude material **2** (15.0 g, 61.1 mmol, 1.00 *eq*) in methanol (250 mL) was added acetaldehyde (33.6 g, 305 mmol, 42.8 mL, 40% purity, 5.00 *eq*) and hydrochloric acid (12 M, 15.3 mL, 3.00 *eq*). The mixture was stirred at 50 °C for 16 h. The mixture was concentrated under reduced pressure to give a residue. The residue was triturated with a solution of petroleum ether/ethyl acetate (5/1, 200 mL) at 25 °C for 2 h. The mixture was filtered, and the filter cake was washed with petroleum ether (50 mL). The filter cake was collected and dried under reduced pressure to give compound **3** (15.5 g, 43.9 mmol, 72% yield) as a white solid.

Compound **3**: ^1^H NMR (400 MHz, DMSO-*d*_6_) δ = 9.90 - 9.46 (m, 2H), 7.15 - 7.03 (m, 1H), 6.75 - 6.68 (m, 1H), 6.65 - 6.60 (m, 1H), 4.61 - 4.43 (m, 2H), 4.33 - 4.19 (m, 2H), 3.28 - 3.16 (m, 2H), 1.65 - 1.53 (m, 3H), 1.38 - 1.14 (m, 3H). MS (ESI) m/z 236.1 [M+H]^+^

To a solution of **3** (15.5 g, 50.7 mmol, 1.00 *eq*) in dichloromethane (120 mL) were added triethylamine (15.4 g, 152 mmol, 21.2 mL, 3.00 *eq*) and trifluoromethanesulfonic anhydride (42.9 g, 152 mmol, 25.1 mL, 3.00 *eq*) at 0 °C. The mixture was stirred at 25 °C for 16 h. The mixture was concentrated under reduced pressure. The crude product was purified by flash silica gel chromatography (ISCO^®^; 220 g SepaFlash^®^ Silica Flash Column, Eluent of 0∼10% Ethyl acetate/Petroleum ether gradient at 80 mL/min) to give compound **4** (6.30 g, 12.5 mmol, 25% yield) as yellow oil.

Compound **4**: ^1^H NMR (400 MHz, DMSO-*d*_6_) δ = 7.63 - 7.46 (m, 2H), 7.44 - 7.36 (m, 1H), 5.44 - 5.19 (m, 1H), 5.07 - 4.70 (m, 1H), 4.27 - 3.85 (m, 2H), 3.57 - 3.40 (m, 2H), 1.51 (d, *J* = 6.8 Hz, 3H), 1.27 - 0.85 (m, 3H). MS (ESI) m/z 500.0 [M+H]^+^

To a solution of **4** (5.50 g, 11.0 mmol, 1.00 *eq*) in *N*,*N*-dimethylformamide (50 mL) were added triethylsilane (3.84 g, 33.0 mmol, 5.28 mL, 3.00 *eq*), palladium(II) acetate (494 mg, 2.20 mmol, 0.200 *eq*) and 1,3-bis(diphenylphosphaneyl)propane (908 mg, 2.20 mmol, 0.200 *eq*). The system was purged with nitrogen for three times. The mixture was stirred at 60 °C for 16 h under nitrogen atmosphere. After being cooled to room temperature, the reaction mixture was diluted with water (100 mL) and extracted with ethyl acetate (3 × 100 mL). The combined organic layers were washed with brine (120 mL), dried over anhydrous sodium sulfate, filtered and concentrated under reduced pressure to give a residue. The crude product was purified by reversed phase column (C18, 330 g, flow: 100 mL/min; gradient: from 65-70% water (0.1% formic acid) in acetonitrile over 60 min. to give intermediate **5** (3.22 g, 9.07 mmol, 82% yield) as a yellow solid.

Compound **5**: ^1^H NMR (400 MHz, DMSO-*d*_6_) δ = 7.39 - 7.15 (m, 4H), 5.29 - 5.08 (m, 1H), 4.98 - 4.63 (m, 1H), 4.25 - 3.85 (m, 2H), 3.31 - 3.15 (m, 2H), 1.58 - 1.43 (m, 3H), 1.29 - 0.84 (m, 3H). MS (ESI) m/z 352.0 [M+H]^+^

To a suspension of aluminum (III) lithium hydride (3.40 g, 9.68 mmol, 1.00 *eq*) in toluene (40 mL) was added compound **5** (2.5 M, 19.4 mL, 5.00 *eq*) at 0 °C under nitrogen atmosphere. The mixture was stirred at 120 °C for 2 h under nitrogen atmosphere. The mixture was cooled to 0 °C and was quenched with water (2 mL) followed by 15% sodium hydroxide solution (2 mL) and water (6 mL) at 0 °C. The reaction mixture was diluted with water (80 mL) and extracted with ethyl acetate (3 × 80 mL). The combined organic layers were washed with brine (100 mL), dried over anhydrous sodium sulfate, filtered and concentrated under reduced pressure to give 1.90g of crude **6** as a white solid. 450 mg of crude **6** (2.54 mmol, 1.00 *eq*) and 1.16g of 2,3,4,6,7,8,9,10-octahydropyrimido[1,2-*a*]azepine (7.62 mmol, 1.15 mL, 3.00 *eq*) in dichloromethane (8 mL) were added to a solution of (2-chlorophenyl)methanesulfonyl chloride (857 mg, 3.81 mmol, 1.50 *eq*) in dichloromethane (2 mL) dropwise at 0 °C. The mixture was stirred at 0 °C for 3 h.

The reaction mixture was concentrated under reduced pressure. The residue was purified by *Prep*-HPLC (column: Waters Xbridge Prep OBD C18 150 × 40 mm × 10 µm; mobile phase: [water (ammonium bicarbonate) - acetonitrile]; gradient: 35%-65% B over 20 min) to give compound **7** (130 mg, 352 μmol, 14% yield) as a yellow solid.

Compound **7**: ^1^H NMR (400 MHz, CDCl_3_) δ = 7.58 - 7.40 (m, 1H), 7.35 - 7.29 (m, 2H), 7.26 - 7.15 (m, 4H), 7.14 - 7.02 (m, 1H), 5.04 - 4.88 (m, 1H), 4.64 - 4.41 (m, 1H), 4.25 - 4.11 (m, 1H), 4.01 (q, *J* = 6.4 Hz, 1H), 3.77 (dd, *J* = 2.0, 6.0 Hz, 1H), 3.42 - 3.32 (m, 2H), 3.13 - 2.95 (m, 1H), 2.69 (s, 1H), 1.62 - 1.48 (m, 3H). MS (ESI) m/z 364.2 [M-H]^−^

To a solution of **7** (130 mg, 355 μmol, 1.00 *eq*) in phosphate buffer (3 mL) and acetonitrile (3 mL) was added 2,2,6,6-tetramethylpiperidinooxy (44.7 mg, 284 μmol, 0.800 *eq*) followed by the solution of sodium chlorite (64.3 mg, 711 μmol, 2.00 *eq*) in water (0.75 mL) and sodium hypochlorite (529 mg, 355 μmol, 439 μL, 5% purity, 1.00 *eq*) in portions at 0 °C. The mixture was stirred at 25 °C for 2 h. The reaction mixture was quenched with 1M hydrochloric acid and the mixture was adjusted to pH = 3-4. Then the mixture was diluted with ethyl acetate (20 mL) and water (20 mL).

The layers were separated, and the aqueous phase was extracted with ethyl acetate (2 × 20 mL). The combined organic layers were washed with brine (30 mL), dried over sodium sulfate, filtered and concentrated under reduced pressure to give compound **8** (100 mg, 258 μmol, 73% yield) as yellow oil.

Compound **8**: ^1^H NMR (400 MHz, CDCl_3_) δ = 7.64 - 7.43 (m, 1H), 7.33 (d, *J* = 7.6 Hz, 2H), 7.25 - 7.10 (m, 4H), 7.07 - 6.90 (m, 1H), 5.05 - 4.86 (m, 1H), 4.75 - 4.61 (m, 1H), 4.56 - 4.22 (m, 2H), 3.31 - 3.21 (m, 1H), 3.18 - 3.09 (m, 1H), 1.62 - 1.42 (m, 3H). MS (ESI) m/z 378.0 [M-H]^−^

To a solution of compound **8** (100 mg, 263 μmol, 1.00 *eq*) in *N*,*N*-dimethylformamide (3 mL) was added *N*,*N*-diisopropylethylamine (102 mg, 790 μmol, 138 μL, 3.00 *eq*) followed by *O*-(7-azabenzotriazol-1-yl)-*N*,*N*,*N’*,*N’*-tetramethyluronium hexafluorophosphate (150 mg, 395 μmol, 1.50 *eq*) at 0 °C. After stirring at 0 °C for 30 min, 1-(7-(aminomethyl)imidazo[1,2-*a*]pyridin-3-yl)dihydropyrimidine-2,4(1*H*,3*H*)- dione (27.75 mg, 342.23 μmol, 1.30 *eq*) was added to the mixture. The resulting mixture was stirred at 25 °C for 1 h. The reaction mixture was quenched with water (20 mL) and diluted with ethyl acetate (20 mL). The layers were separated, and the aqueous phase was extracted with ethyl acetate (2 × 20 mL). The combined organic layers were washed with saturated sodium bicarbonate solution (2 × 30 mL) followed by brine (30 mL), dried over sodium sulfate, filtered and concentrated under reduced pressure. The residue was purified by *Prep*-HPLC (column: Welch Xtimate C18 150 × 25 mm × 5 μm; mobile phase: [water (formic acid) - acetonitrile]; gradient: 10%- 40% B over 15 min) and lyophilized to **MRT-5702** (120 mg, 191 μmol, 73% yield) as a white solid.

Compound **MRT-5702**: ^1^H NMR (400 MHz, DMSO-*d*_6_) δ = 10.90 - 10.76 (m, 1H), 8.82 - 8.47 (m, 2H), 8.17 - 7.99 (m, 1H), 7.74 - 7.44 (m, 2H), 7.43 - 7.24 (m, 6H), 7.20 - 6.92 (m, 2H), 5.11 - 4.96 (m, 1H), 4.88 - 4.63 (m, 1H), 4.61 - 4.41 (m, 2H), 4.38 - 4.14 (m, 2H), 3.84 (t, *J* = 6.4 Hz, 2H), 3.20 - 3.06 (m, 2H), 2.85 (d, *J* = 2.0 Hz, 2H), 1.59 - 1.42 (m, 3H). MS (ESI) m/z 621.3 [M+H]^+^

### SFC Separation of MRT-5702

**Scheme 2:**
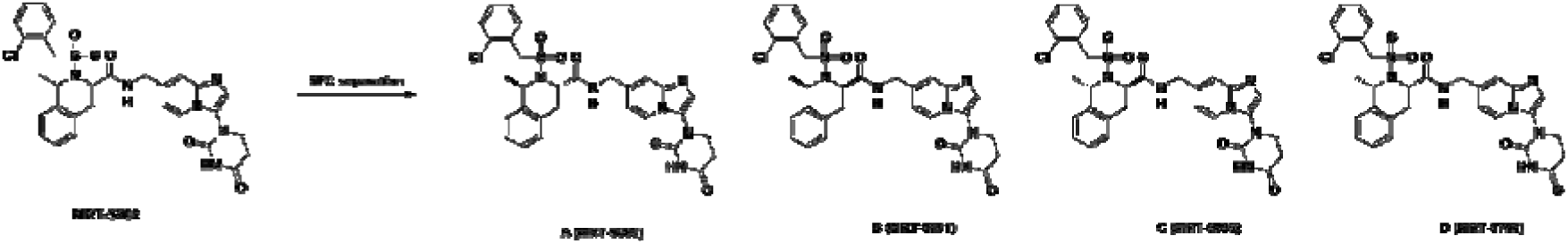
SFC Separation of MRT-5702. Configuration of MRT-9800 (MRT-5702A) and MRT-9799 (MRT-5702C) determined by X-Ray crystallography. The configuration of MRT-9798 (MRT-5702C) was access by modeling and MRT-9801 (MRT-5702B) is the remaining configuration.

**MRT-5702** (210 mg, 338 μmol, 1.00 *eq*) was separated by Chiral SFC (column: (s,s) WHELK-O1 (250 mm × 30 mm,10 μm); mobile phase: [carbon dioxide- propan-2-ol / acetonitrile]; B%: 62%, isocratic elution mode) to give Peak 1 and **B**. **B** was purified by *Prep*-HPLC (column: Phenomenex luna C18 150 × 25 mm × 10 μm; mobile phase: [water (formic acid) - acetonitrile]; gradient: 15%-45% B over 1 min).

Compound **B** (31.3 mg, 49.9 μmol, 15% yield) was obtained as a white solid. Peak1 was separated by Chiral SFC (column: DAICEL CHIRALPAK AS (250 mm × 30 mm, 10 μm); mobile phase: [carbon dioxide - ethanol (0.1% ammonia hydrate)]; B%:40%, isocratic elution mode) to give Peak 2 and **A**. **A** was purified by *Prep*-HPLC (column: Welch Xtimate C18 150 × 25 mm × 5 μm; mobile phase: [water (formic acid) - acetonitrile]; gradient: 10%-40% B over 15 min). Compound **A** (20.7 mg, 33.0 μmol, 10% yield) was obtained as a white solid. Peak 2 was separated by Chiral SFC (column: DAICEL CHIRALCEL OJ (250 mm × 30 mm, 10 μm); mobile phase: [carbon dioxide - ethanol / acetonitrile]; B%: 50%, isocratic elution mode) to give **compounds C** and **D**. **C** was purified by *Prep*-HPLC (column: Welch Xtimate C18 150 × 25 mm × 5 μm; mobile phase: [water (formic acid) - acetonitrile]; gradient: 13%-43% B over 10 min). Compound **C** (18.6 mg, 29.7 μmol, 9% yield) was obtained as a white solid. **D** was purified by *Prep*-HPLC (column: Welch Xtimate C18 150 × 25 mm × 5 μm; mobile phase: [water (formic acid) - acetonitrile]; gradient: 10%-40% B over 10 min). Compound **D** (29.0 mg, 46.2 μmol, 14% yield) was obtained as a white solid.

Compound **A**: ^1^H NMR (400 MHz, DMSO-*d*_6_) δ = 10.66 (s, 1H), 8.51 (t, *J* = 6.0 Hz, 1H), 8.13 (d, *J* = 7.2 Hz, 1H), 7.59 (dd, *J* = 2.0, 7.2 Hz, 1H), 7.55 - 7.46 (m, 2H), 7.43 - 7.32 (m, 2H), 7.24 - 7.08 (m, 5H), 6.56 (d, *J* = 7.2 Hz, 1H), 5.04 (q, *J* = 6.8 Hz, 1H), 4.82 (d, *J* = 13.6 Hz, 1H), 4.74 (d, *J* = 3.2 Hz, 1H), 4.61 (d, *J* = 13.6 Hz, 1H), 4.18 (d, *J* = 6.0 Hz, 2H), 3.77 (t, *J* = 6.8 Hz, 2H), 3.45 (dd, *J* = 5.6, 15.6 Hz, 1H), 3.18 - 3.06 (m, 1H), 2.82 (t, *J* = 6.4 Hz, 2H), 1.46 (d, *J* = 6.4 Hz, 3H). MS (ESI) m/z 621.3 [M+H]^+^

Compound **B**: ^1^H NMR (400 MHz, DMSO-*d*_6_) δ = 10.65 (s, 1H), 8.59 (t, *J* = 5.6 Hz, 1H), 8.25 (d, *J* = 6.8 Hz, 1H), 7.50 (s, 1H), 7.46 - 7.39 (m, 2H), 7.37 - 7.21 (m, 7H), 6.91 (dd, *J* = 1.2, 7.2 Hz, 1H), 5.01 (d, *J* = 7.2 Hz, 1H), 4.50 - 4.38 (m, 3H), 4.35 (d, *J* = 13.6 Hz, 1H), 4.20 (d, *J* = 13.6 Hz, 1H), 3.78 (t, *J* = 6.4 Hz, 2H), 3.23 - 3.06 (m, 2H), 2.82 (t, *J* = 5.6 Hz, 2H), 1.54 (d, *J* = 7.2 Hz, 3H). MS (ESI) m/z 621.3 [M+H]^+^

Compound **C**: ^1^H NMR (400 MHz, DMSO-*d*_6_) δ = 10.66 (s, 1H), 8.51 (t, *J* = 5.6 Hz, 1H), 8.14 (d, *J* = 7.2 Hz, 1H), 7.59 (dd, *J* = 1.6, 7.2 Hz, 1H), 7.54 - 7.47 (m, 2H), 7.43 - 7.33 (m, 2H), 7.25 - 7.09 (m, 5H), 6.58 (d, *J* = 7.2 Hz, 1H), 5.04 (q, *J* = 6.4 Hz, 1H), 4.82 (d, *J* = 13.6 Hz, 1H), 4.74 (d, *J* = 3.2 Hz, 1H), 4.61 (d, *J* = 13.6 Hz, 1H), 4.19 (d, *J* = 6.0 Hz, 2H), 3.77 (t, *J* = 6.4 Hz, 2H), 3.48 - 3.42 (m, 1H), 3.12 (dd, *J* = 2.0, 15.6 Hz, 1H), 2.83 (t, *J* = 6.4 Hz, 2H), 1.46 (d, *J* = 6.4 Hz, 3H).

Compound **D**: ^1^H NMR (400 MHz, DMSO-*d*_6_) δ = 10.65 (s, 1H), 8.58 (t, *J* = 6.0 Hz, 1H), 8.26 (d, *J* = 6.8 Hz, 1H), 7.51 (s, 1H), 7.47 - 7.38 (m, 2H), 7.37 - 7.21 (m, 7H), 6.92 (dd, *J* = 1.2, 7.2 Hz, 1H), 5.02 (q, *J* = 7.2 Hz, 1H), 4.52 - 4.38 (m, 3H), 4.35 (d, *J* = 13.6 Hz, 1H), 4.20 (d, *J* = 13.6 Hz, 1H), 3.78 (t, *J* = 6.4 Hz, 2H), 3.28 - 3.22 (m, 1H), 3.16 - 3.07 (m, 1H), 2.82 (t, *J* = 5.6 Hz, 2H), 1.54 (d, *J* = 7.2 Hz, 3H). MS (ESI) m/z 621.1 [M+H]^+^

### X-Ray

#### Equipment used for X-Ray

Rigaku Oxford Diffraction XtaLAB Synergy-S equipped with a HyPix-6000HE area detector Cryogenic system: Oxford Cryostream 800 Cu: λ=1.54184 Å, 50W Distance from the crystal to the CCD detector: d = 35 mm Tube Voltage: 50 kV; Tube Current: 1 mA 39

### MRT-5702A

**Scheme 3:**
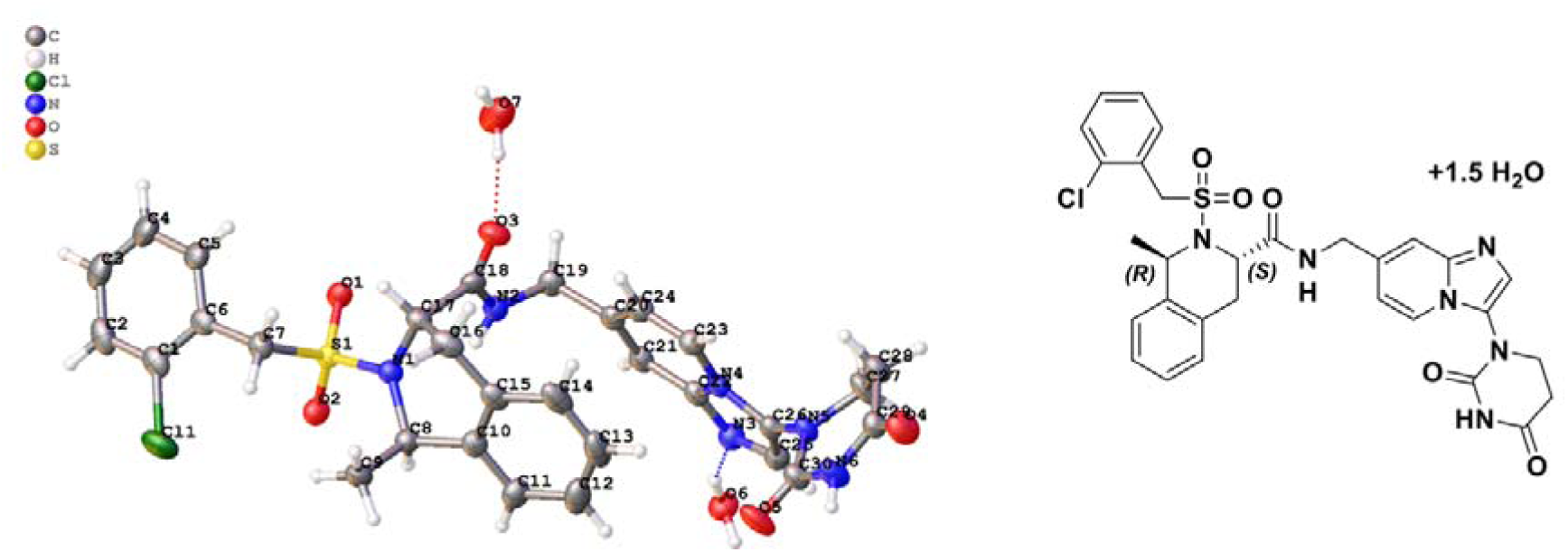
X-Ray Structure of MRT-5702A (MRT-9800) in (*R,S*) configuration.

The symmetry of the crystal structure was assigned the monoclinic space group C2 with the following parameters: a = 21.3601(3) Å, b = 11.2017(2) Å, c = 12.5762(2) Å, α = 90°, β = 97.521(2)°, γ = 90°, V = 2983.21(8) Å3, Z = 4, Dc = 1.443 g/cm^3^, μ(CuKα) = 2.272 mm^−1^, and F(000) = 1356.0.

Colorless block-shaped single crystals of MRT-5702A were obtained from methanol - water (4:1). A suitable crystal 0.30 × 0.10 × 0.10 mm3 was selected for testing. Data collection temperature: T = 149.99(10) K. Total of 18187 reflections were collected in the 2θ range from 7.09 to 133.17. The limiting indices were: −24 ≤ h ≤ 25, −13 ≤ k ≤ 13, −14 ≤ l ≤ 14; which yielded 5130 unique reflections (Rint = 0.0690). The structure was solved using SHELXT (Sheldrick, G. M. 2015. Acta Cryst. A71, 3-8) and refined using SHELXL (against F²) (Sheldrick, G. M. 2015. Acta Cryst. C71, 3-8). The total number of refined parameters was 408, compared with 5130 data. All reflections were included in the refinement. The goodness of fit on F² was 1.058 with a final R value for [I >= 2σ (I)] R_1_ = 0.0368 and wR_2_ = 0.0966. The largest differential peak and hole were 0.40 and −0.34 eÅ^−3^.

### Experimental

The crystal was kept at 149.99(10) K during data collection. Using Olex2^63^, the structure was solved with the SHELXT^64^ structure solution program using Intrinsic Phasing and refined with the SHELXL^65^ refinement package using Least Squares minimisation.

### Crystal structure determination

**Crystal Data** for C_30_H_32_ClN_6_O_6.5_S (*M* =648.12 g/mol): monoclinic, space group C2 (no. 5), *a* = 21.3601(3) Å, *b* = 11.2017(2) Å, *c* = 12.5762(2) Å, β = 97.521(2)°, *V* = 2983.21(8) Å^3^, *Z* = 4, *T* = 149.99(10) K, μ(Cu Kα) = 2.272 mm^−1^, *Dcalc* = 1.443 g/cm^3^, 18187 reflections measured (7.09° ≤ 2Θ ≤ 133.17°), 5130 unique (*R*_int_ = 0.0690, R_sigma_ = 0.0480) which were used in all calculations. The final *R*_1_ was 0.0368 (I > 2σ(I)) and *wR*_2_ was 0.0976 (all data).

### Refinement model description

Number of restraints - 1, number of constraints - unknown. Details:

1. Fixed Uiso

At 1.2 times of: All C(H) groups, All C(H,H) groups, All N(H) groups At 1.5 times of: All C(H,H,H) groups, All O(H,H) groups

1. 2. Restrained distances O7-O3 2.751 with sigma of 0.01
2. Others

Fixed Sof: H7C(0.5) H7D(0.5)

Fixed X: O7(0.5)

Fixed Y: O7(0.739329) Fixed Z: O7(0.5)

1. 4. a Free rotating group: O6(H6A,H6B), O7(H7C,H7D)

4.b Ternary CH refined with riding coordinates: C8(H8), C17(H17)

4.c Secondary CH2 refined with riding coordinates: C7(H7A,H7B), C19(H19A,H19B), C16(H16A,H16B), C27(H27A,H27B), C28(H28A,H28B)

4.d Aromatic/amide H refined with riding coordinates: N2(H2), N6(H6), C24(H24), C23(H23), C25(H25), C21(H21), C11(H11), C2(H2A), C14(H14), C13(H13), C3(H3), C12(H12), C4(H4), C5(H5)

4.e Idealised Me refined as rotating group: C9(H9A,H9B,H9C)

### MRT-5702C

**Scheme 4:**
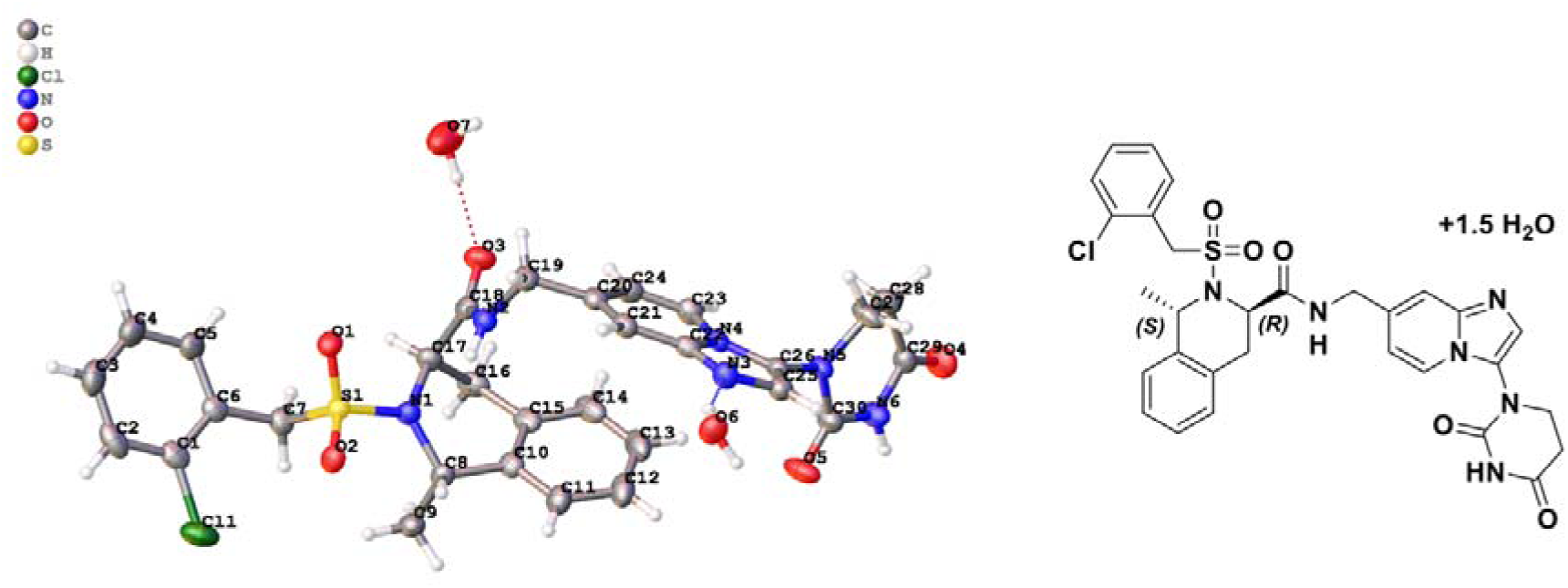
X-Ray Structure of MRT-5702C (MR-9799) in (*S,R*) configuration.

The symmetry of the crystal structure was assigned the monoclinic space group C2 with the following parameters: a = 21.3441(3) Å, b = 11.1992(2) Å, c = 12.5765(2) Å, α = 90°, β = 97.5310(10)°, γ = 90°, V = 2980.31(8) Å3, Z = 4, Dc = 1.444 g/cm^3^, μ(CuKα) = 2.274 mm^−1^, and F(000) = 1356.0.

Colorless block-shaped single crystals of MRT-5702C were obtained from methanol −water (4:1). A suitable crystal 0.25 × 0.10 × 0.10 mm^3^ was selected for testing. Data collection temperature: T = 150.00(10) K. Total of 18726 reflections were collected in the 2θ range from 7.09 to 133.186°. The limiting indices were: −25 ≤ h ≤ 25, −13 ≤ k ≤ 13, −14 ≤ l ≤ 14; which yielded 5178 unique reflections (Rint = 0.0453). The structure was solved using SHELXT (Sheldrick, G. M. 2015. Acta Cryst. A71, 3-8) and refined using SHELXL (against F²) (Sheldrick, G. M. 2015. Acta Cryst. C71, 3-8). The total number of refined parameters was 408, compared with 5178 data. All reflections were included in the refinement. The goodness of fit on F² was 1.047 with a final R value for [I >= 2σ (I)] R_1_ = 0.0331 and wR_2_ = 0.0847. The largest differential peak and hole were 0.35 and −0.32 eÅ^−3^.

### Experimental

The crystal was kept at 150.00(10) K during data collection. Using Olex2^63^, the structure was solved with the SHELXT^64^ structure solution program using Intrinsic Phasing and refined with the SHELXL^65^ refinement package using Least Squares minimization.

### Crystal structure determination

**Crystal Data** for C_30_H_32_ClN_6_O_6.5_S (*M* =648.12 g/mol): monoclinic, space group C2 (no. 5), *a* = 21.3441(3) Å, *b* = 11.1992(2) Å, *c* = 12.5765(2) Å, β = 97.5310(10)°, *V* = 2980.31(8) Å^3^, *Z* = 4, *T* = 150.00(10) K, μ(Cu Kα) = 2.274 mm^−1^, *Dcalc* = 1.444 g/cm^3^, 18726 reflections measured (7.09° ≤ 2Θ ≤ 133.186°), 5178 unique (*R*_int_ = 0.0453, R_sigma_ = 0.0382) which were used in all calculations. The final *R*_1_ was 0.0331 (I > 2σ(I)) and *wR*_2_ was 0.0856 (all data).

### Refinement model description

Number of restraints - 2, number of constraints - unknown.

Details:

1. Fixed Uiso At 1.2 times of:

All C(H) groups, All C(H,H) groups, All N(H) groups

At 1.5 times of: All C(H,H,H) groups, All O(H,H) groups

1. Restrained distances O3-O7

2.748 with sigma of 0.001 O3-H7C

1.884 with sigma of 0.001

1. Others

Fixed Sof: H7C(0.5) H7D(0.5) Fixed X: O7(0.5)

Fixed Y: O7(0.261271) Fixed Z: O7(0.5)

1. Free rotating group:

O6(H6A,H6B), O7(H7C,H7D)

4.a Ternary CH refined with riding coordinates:

C8(H8), C17(H17)

4.a Secondary CH2 refined with riding coordinates:

C7(H7A,H7B), C16(H16A,H16B), C19(H19A,H19B), C28(H28A,H28B), C27(H27A,H27B)

4.a Aromatic/amide H refined with riding coordinates:

N2(H2), N6(H6), C23(H23), C24(H24), C21(H21), C11(H11), C25(H25), C2(H2A),

C14(H14), C13(H13), C12(H12), C3(H3), C4(H4), C5(H5)

4.a Idealised Me refined as rotating group: C9(H9A,H9B,H9C)

---

**Extended Data Fig. 1.**
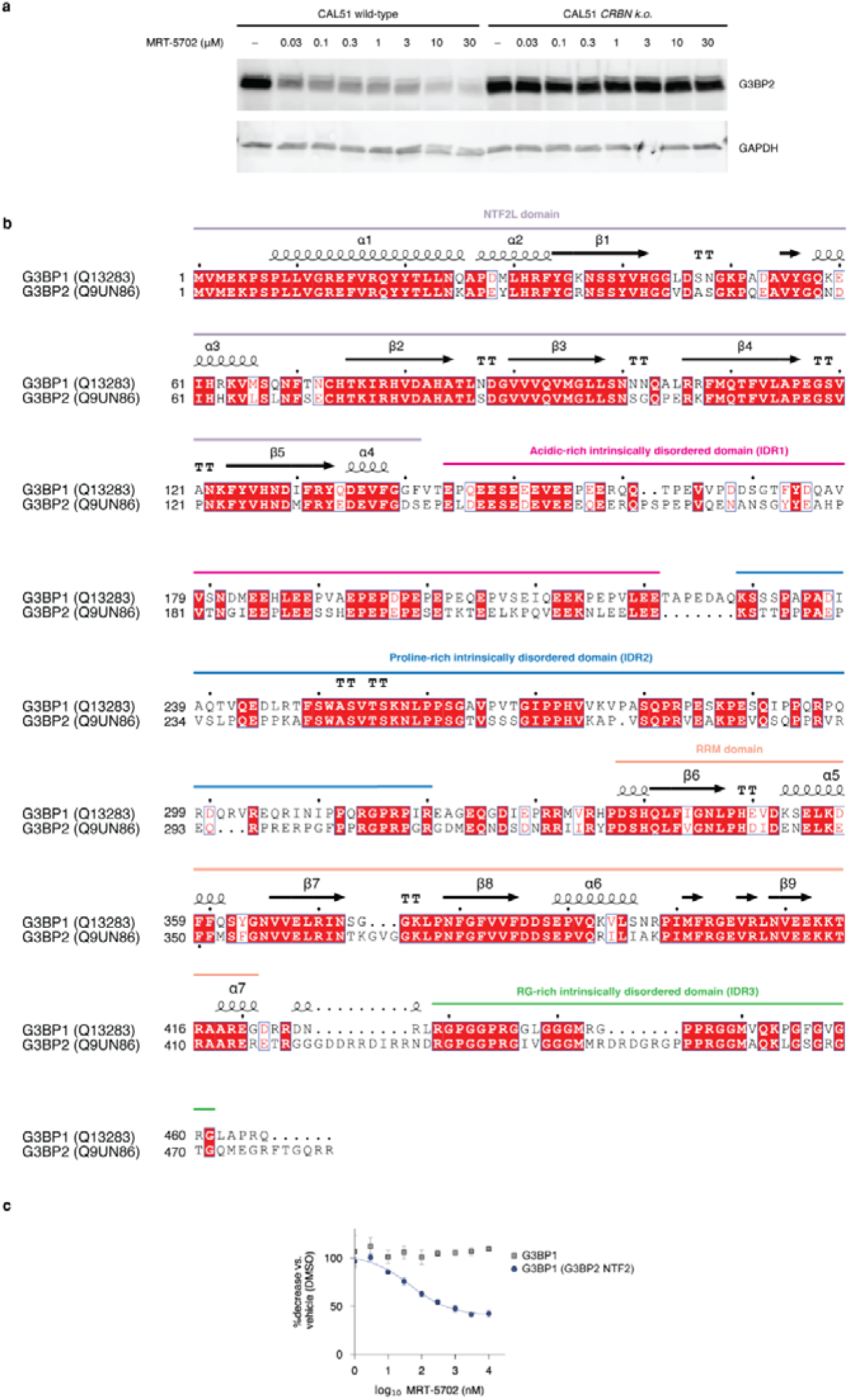
G3BP2 is a G-loop-independent CRBN target. **a**, G3BP2 Western blot in CAL51 cells using MRT-5702 (24 h treatment) and rescue of degradation by genetic *CRBN* knock-out. **b**, Sequence alignment of human G3BP1 and G3BP2 generated with the ENDscript server^17^. Protein domains are annotated. **c**, Luminescent signal derived from overexpressed wild-type HiBiT-G3BP1 and a domain-swapped variant in CAL51 cells measured after 24 h compound treatment.

**Extended Data Fig. 2.**
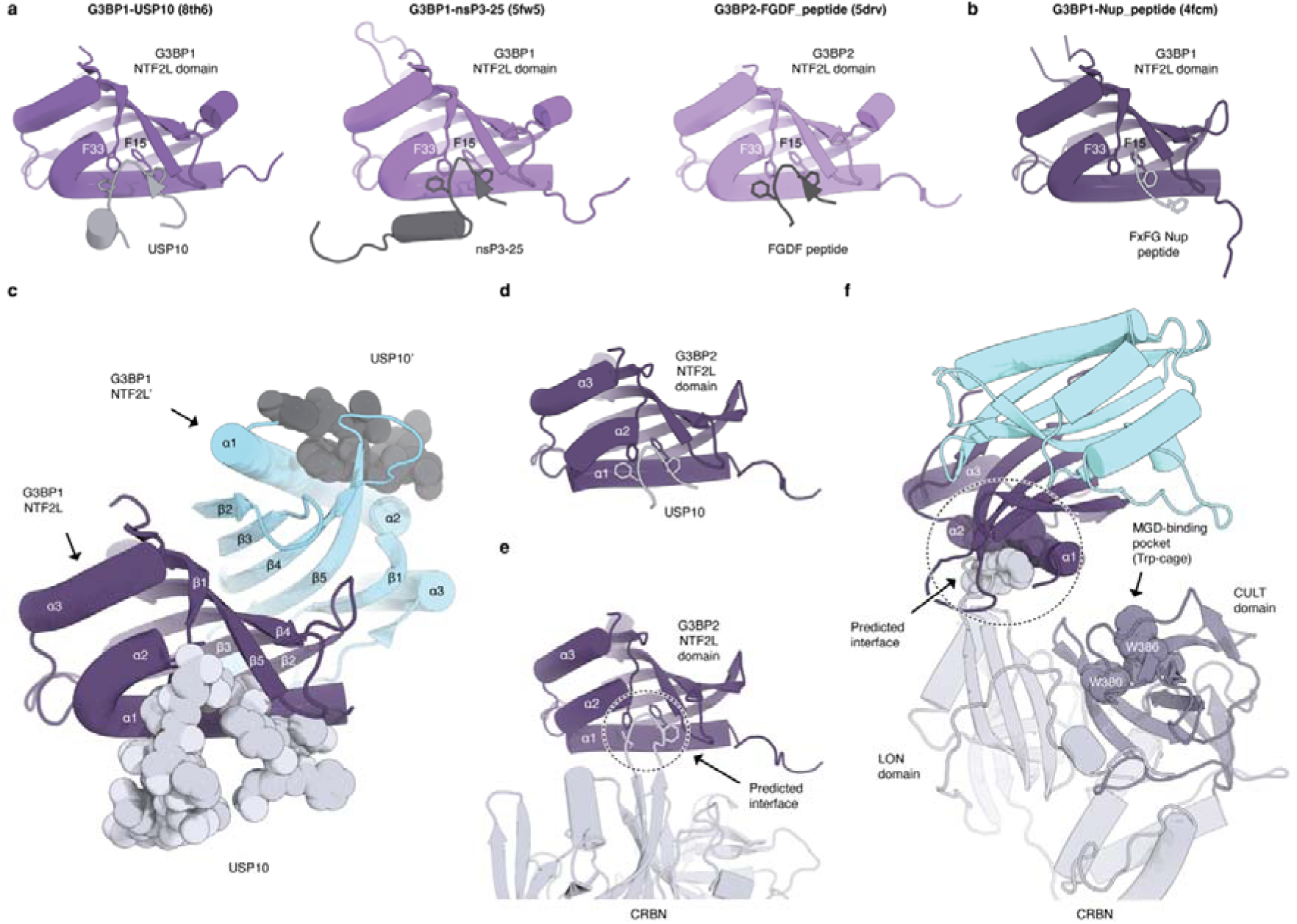
Recognition of FGDF and FxFG motifs by the NTF2L domains of G3BP proteins and ternary complex docking model with CRBN. **a**, Structures of G3BP1 and G3BP2 NTF2L domains bound to FGDF motifs from various endogenous interaction partners. **b**, FxFG motif recognition by the G3BP1 NTF2L domain. **c**, NTF2L homodimer of G3BP1 bound to two individual USP10 peptides (PDB: 7xhf). **d**, Model of a G3BP2 NTF2L domain (PDB: 5drv) bound to the FGDF motif of USP10. **e**, Mimicry-guided docking model of CRBN engaging the FGDF binding site on the G3BP2 NTF2L domain through the LON domain residues F150 and I152. **f**, Mimicry-guided docking model of the G3BP2 NTF2L homodimer with closed CRBN (PDB: 5fqd). The three tryptophan residues of the MGD-binding pocket (Trp-cage) in the CRBN CULT domain are shown as spheres.

**Extended Data Fig. 3.**
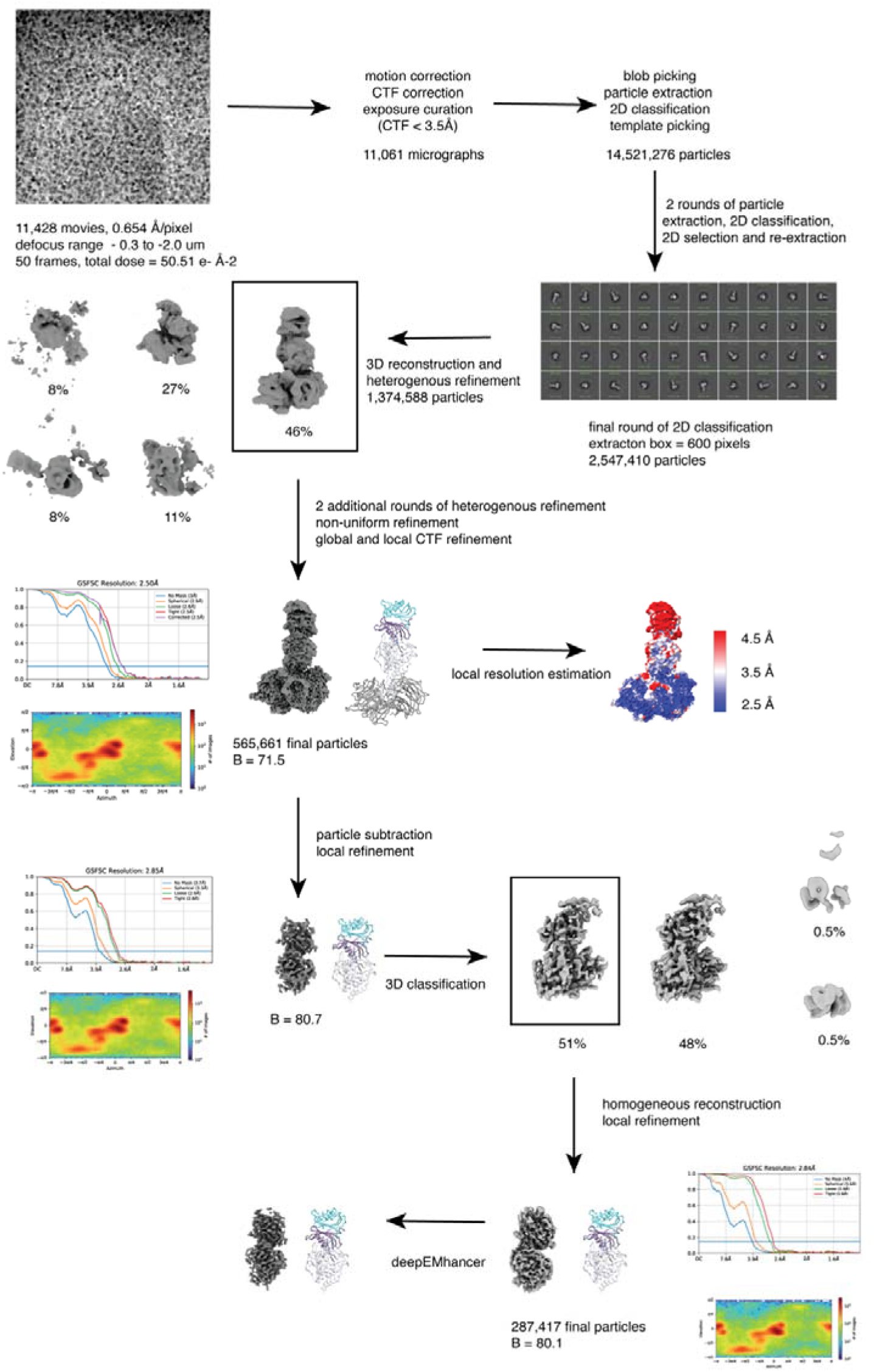
Cryo-EM 3D reconstruction workflow. CryoSPARC v4.2.1 was used for image analysis, 2D classification and 3D reconstruction. Multiple rounds of 2D classification were performed to remove particles not representing the ternary complex. Initial 3D models were created *ab initio* and the best volume was selected for heterogenous refinement alongside decoy volumes generated from aborted *ab initio* jobs. Multiple rounds of heterogenous refinements were performed until most clutter was removed from the volume corresponding to the DDB1/CRBN-MRT-5702-G3BP2^NTF2L^ complex. Non- uniform refinement, local refinement and post-processing were performed as described in Methods. Plots of Fourier shell correlation (GSFSC) and angular distribution for the refinements are shown. The listed resolutions were calculated based on GSFSC at a cutoff of 0.143.

**Extended Data Fig. 4.**
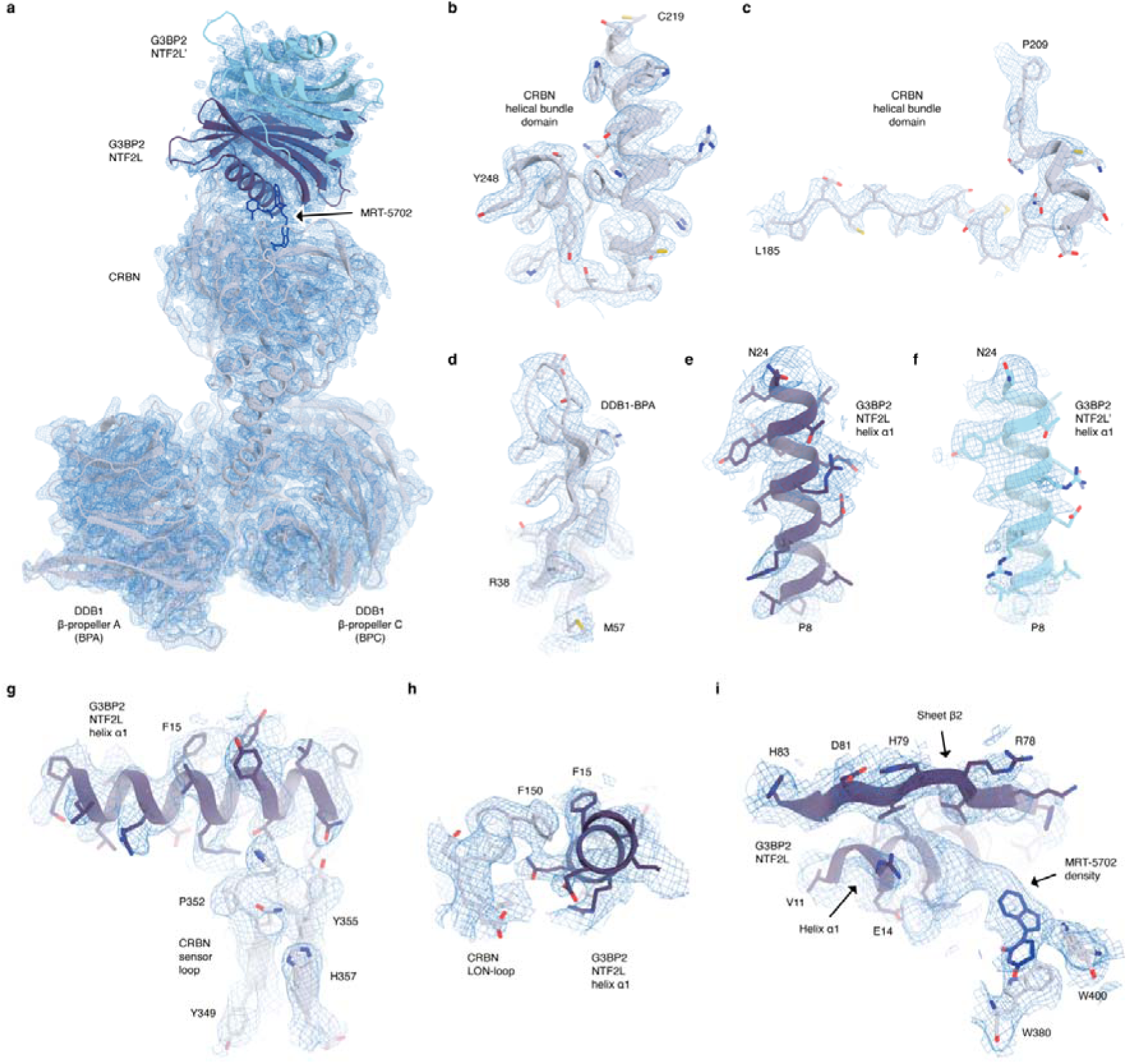
Representative regions of the cryo-EM envelope with the atomic model. **a**, Entire ternary complex structure composed of DDB1^ΔBPB^/CRBN^ΔN^^40^, MRT-5702 and G3BP2^NTF2L^. **b**, Region of the CRBN helical bundle domain (residues 219-248). **c**, Region of the CRBN helical bundle domain (residues 185-209). **d**, Region of DDB1 β-propeller A (residues 38-57). **e**, Helix α1 (residues 8-24) of the interacting G3BP2^NTF2L^ protomer. **f**, Helix α1 (residues 8-24) of the distal G3BP2^NTF2L’^ protomer. **g**, Interface between the CRBN sensor loop and G3BP2^NTF2L^ helix α1. **h**, Interface between the CRBN LON-loop and G3BP2^NTF2L^ helix α1. **i**, Density for MRT-5702 at the interface of the interacting protomer of G3BP2^NTF2L^. The 1- (imidazo[1,2-a]pyridin-3-yl)dihydropyrimidine-2,4(1H,3H)-dione of MRT-5702 is modelled into the density proximal to the MGD- binding site (Trp-cage) in CRBN.

**Extended Data Fig. 5.**
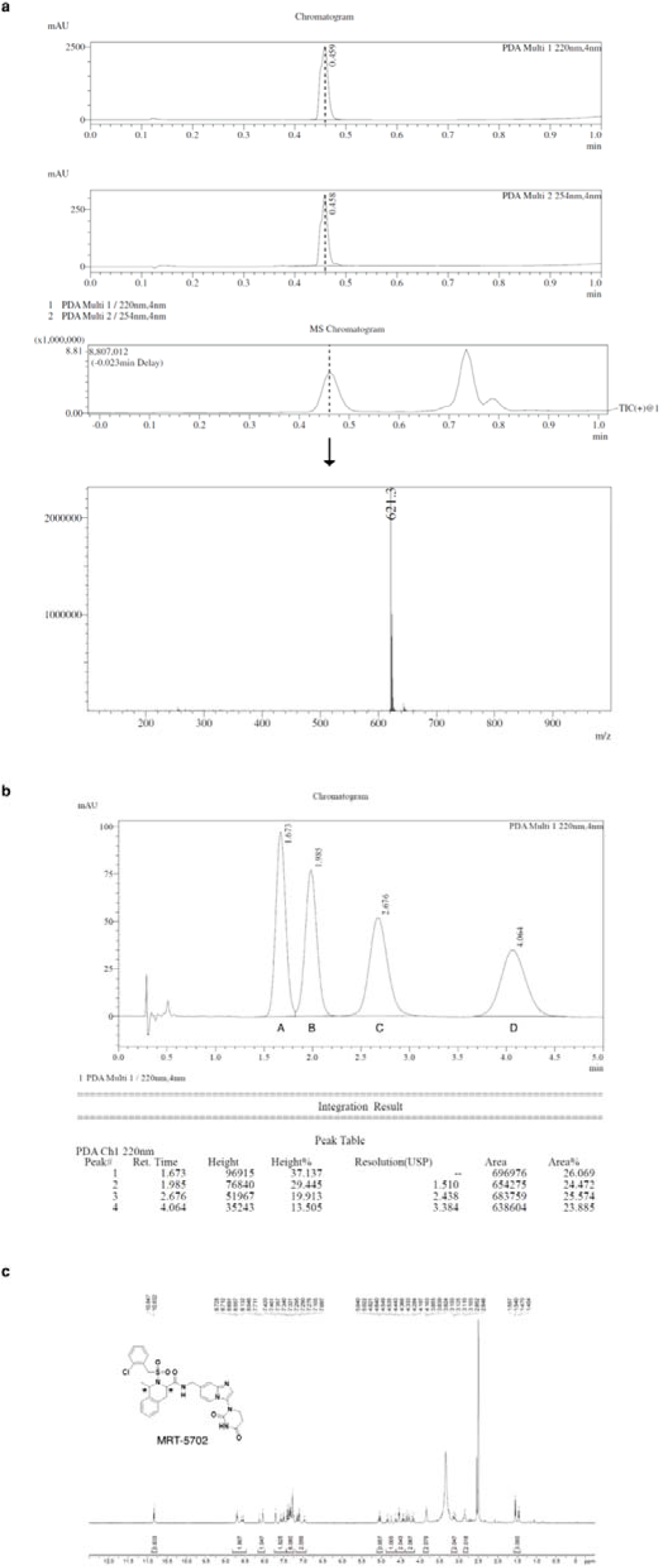
Characterization of MRT-5702 through LC-MS, supercritical fluid chromatography (SFC) and NMR. **a**, LC-MS chromatograms and m/z mass spectrum. **b**, SFC chromatogram. **c**, ^1^H NMR (d^6^-DMSO) of MRT-5702. Chiral centers of MRT-5702 are indicated with asterisks in the chemical structure.

**Extended Data Fig. 6.**
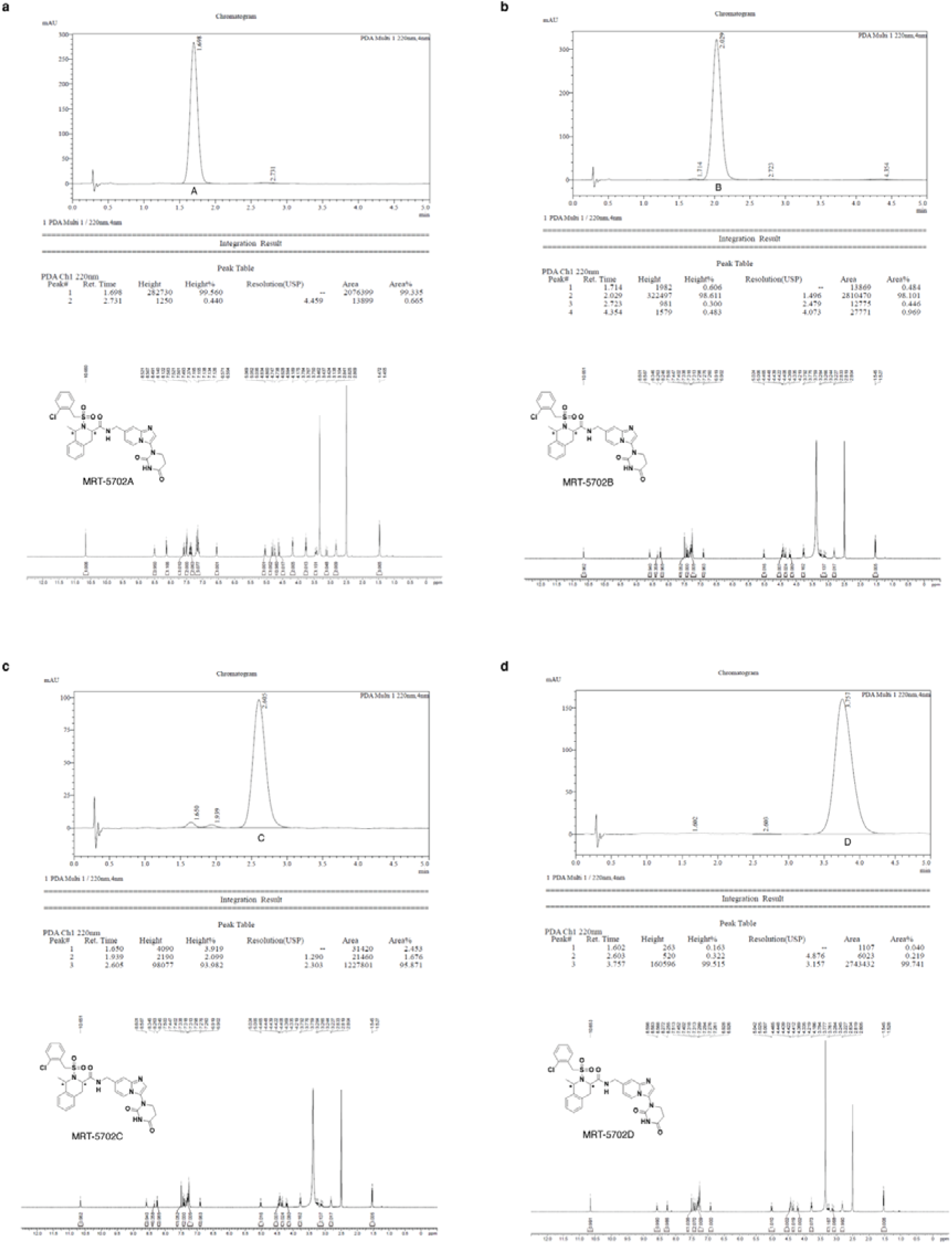
SCF-separated stereoisomers of MRT-5702. **a-d**, SFC chromatogram and ^1^H NMR (d^6^-DMSO) of MRT-5702A, MRT-5702B, MRT-5702C, and MRT-5702D.

**Extended Data Fig. 7.**
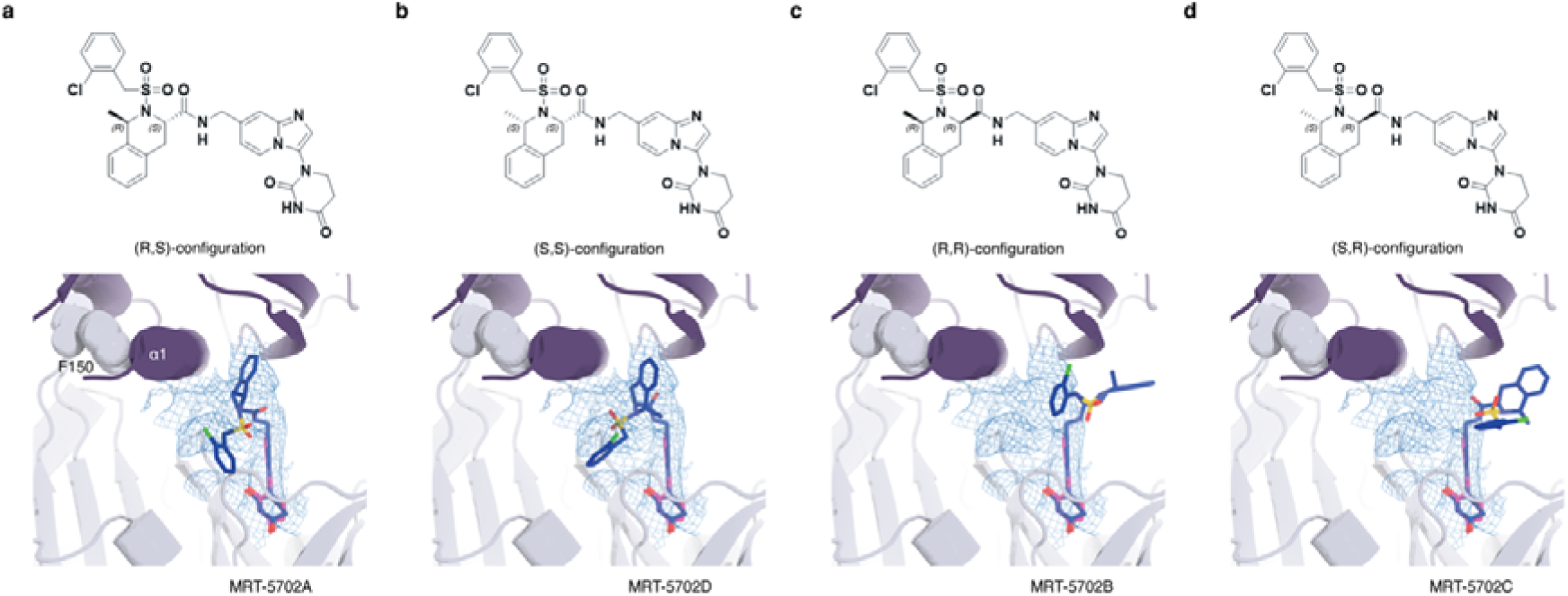
MRT-5702 stereoisomers. **a-d**, Chemical structures of the different configurations of MRT-5702 (top) and corresponding low energy conformers of each ligand configuration producing best overlaps (SMOC score) with the compound volume in the cryo-EM envelope (bottom).

**Extended Data Fig. 8.**
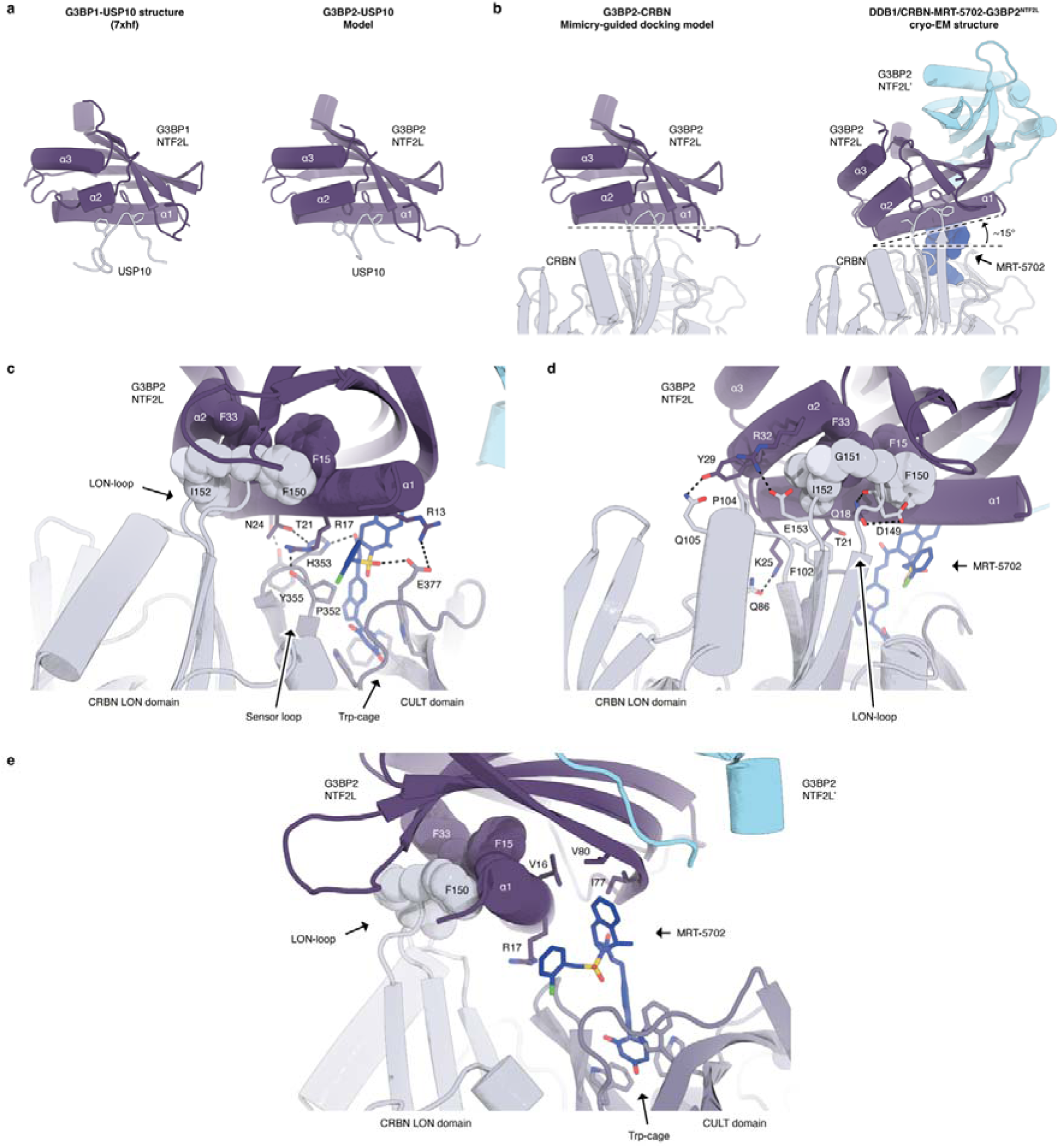
Details of the ternary complex interface. **a**, Comparison of the G3BP1-USP10 structure (PDB: 7xhf) and the generated G3BP2-USP10 model. **b**, Comparison of the mimicry-guided docking model and the CRBN-MRT-5702- G3BP2^NTF2L^ cryo-EM structure. Differences in the pose of the interacting G3BP2^NTF2L^ protomer are highlighted. **c**, Opportunities for interactions between G3BP2^NTF2L^ helix α1, MRT-5702 and the CRBN CULT domain. **d**, Opportunities for interactions between G3BP2^NTF2L^ helix α1, MRT-5702 and the CRBN LON domain. **e**, Detailed interactions between MRT-5702 and G3BP2^NTF2L^.

**Extended Data Table 1:** Cryo-EM data collection, refinement and validation statistics.

|  |  |
| --- | --- |
| **Data collection** |  |
| Microscope | Titan Krios G3i |
| Detector | BioQuantum K3 |
| Magnification | ×130,000 |
| Voltage | 300 kV |
| Electron exposure (e-/ Å<sup>2</sup>) | 50.5056 |
| Defocus range (µm) | -0.3 to -2.0 |
| Pixel size (Å) | 0.654 |
| **Volume reconstruction** |  |
| Refined particles (no.) | 1,374,588 |
| Final particles (no.) | 565,661 |
| Point-group | C1 |
| Resolution (global, Å) |  |
| FSC 0.5 (unmasked/masked) | 3.7/2.9 |
| FSC 0.143 (unmasked/masked) | 3.0/2.5 |
| Resolution range (local, Å) | 2.2 – 7.0 |
| Map sharpening methods | Unsharpened (for model building, refinement and validation)<br>Global sharpening, local resolution refinement and<br>DeepEMhancer (for model building and visualization) |
| **Model composition** |  |
| Protein | 1407 |
| Ligands | 1 |
| **Model refinement** |  |
| Refinement package | phenix.real_space_refine |
| CC | 0.78 (overall) /0.63 (ligand) |
| Average FSC (Å, FSC 0.5) | 2.9 |
| B factors (Å<sup>2</sup>) |  |
| Protein residues | 92.64 |
| Ligands | 125.96 |
| R.m.s. deviations |  |
| Bond lengths (Å) | 0.007 |
| Bond angles (°) | 0.806 |
| **Validation** |  |
| MolProbity score | 1.79 |
| CaBLAM outliers (%) | 1.1 |
| Clashscore | 6.6 |
| Rotamer outliers (%) | 1.82 |
| Cβ outliers (%) | 0 |
| Ramachandran plot |  |
| Favored (%) | 96.53 |
| Allowed (%) | 3.47 |
| Disallowed (%) | 0 |

**Table 1.** Crystal data and structure refinement for EW40755-419-P1B_auto.

|  |  |
| --- | --- |
| Identification code | EW40755-419-P1B_auto |
| Empirical formula | C<sub>30</sub>H<sub>32</sub>ClN<sub>6</sub>O<sub>6.5</sub>S |
| Formula weight | 648.12 |
| Temperature/K | 149.99(10) |
| Crystal system | monoclinic |
| Space group | C2 |
| a/Å | 21.3601(3) |
| b/Å | 11.2017(2) |
| c/Å | 12.5762(2) |
| α/° | 90 |
| β/° | 97.521(2) |
| γ/° | 90 |
| Volume/Å<sup>3</sup> | 2983.21(8) |
| Z | 4 |
| ρ<sub>calc</sub>/g/cm<sup>3</sup> | 1.443 |
| μ/mm<sup>-1</sup> | 2.272 |
| F(000) | 1356.0 |
| Crystal size/mm<sup>3</sup> | 0.3 × 0.1 × 0.1 |
| Radiation | Cu Kα (λ = 1.54184) |
| 2θ range for data collection/° | 7.09 to 133.17 |
| Index ranges | -24 ≤ h ≤ 25, -13 ≤ k ≤ 13, -14 ≤ l ≤ 14 |
| Reflections collected | 18187 |
| Independent reflections | 5130 [R<sub>int</sub> = 0.0690, R<sub>sigma</sub> = 0.0480] |
| Data/restraints/parameters | 5130/1/408 |
| Goodness-of-fit on F<sup>2</sup> | 1.058 |
| Final R indexes [I>=2σ (I)] | R<sub>1</sub> = 0.0368, wR<sub>2</sub> = 0.0966 |
| Final R indexes [all data] | R<sub>1</sub> = 0.0380, wR<sub>2</sub> = 0.0976 |
| Largest diff. peak/hole / e Å<sup>-3</sup> | 0.40/-0.34 |
| Flack parameter | 0.028(10) |

**Table 2.**
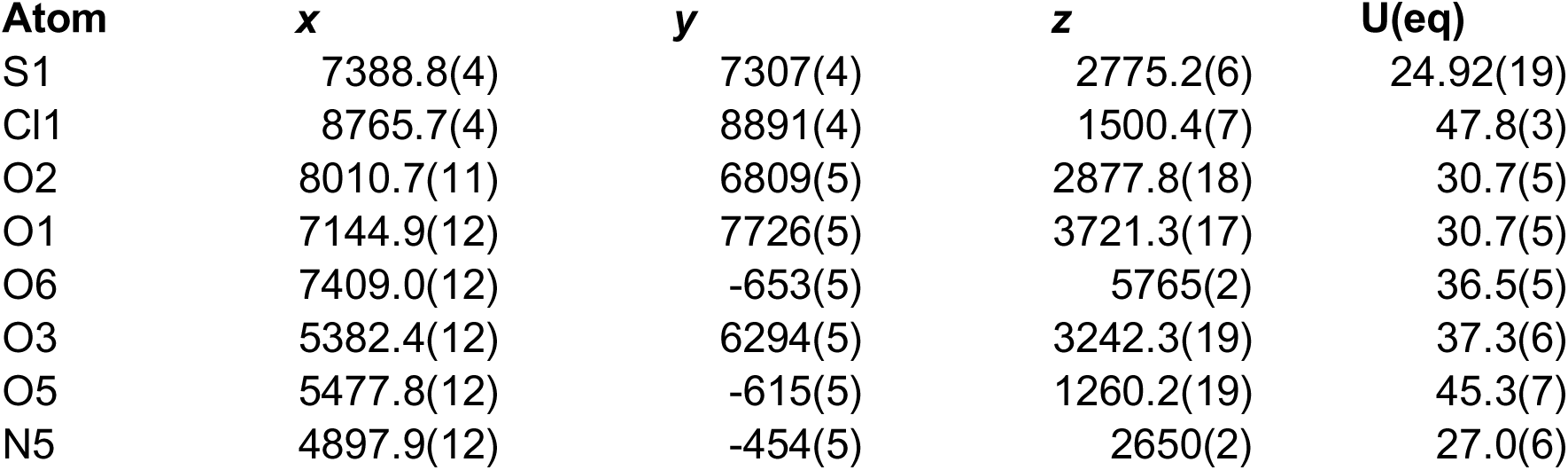
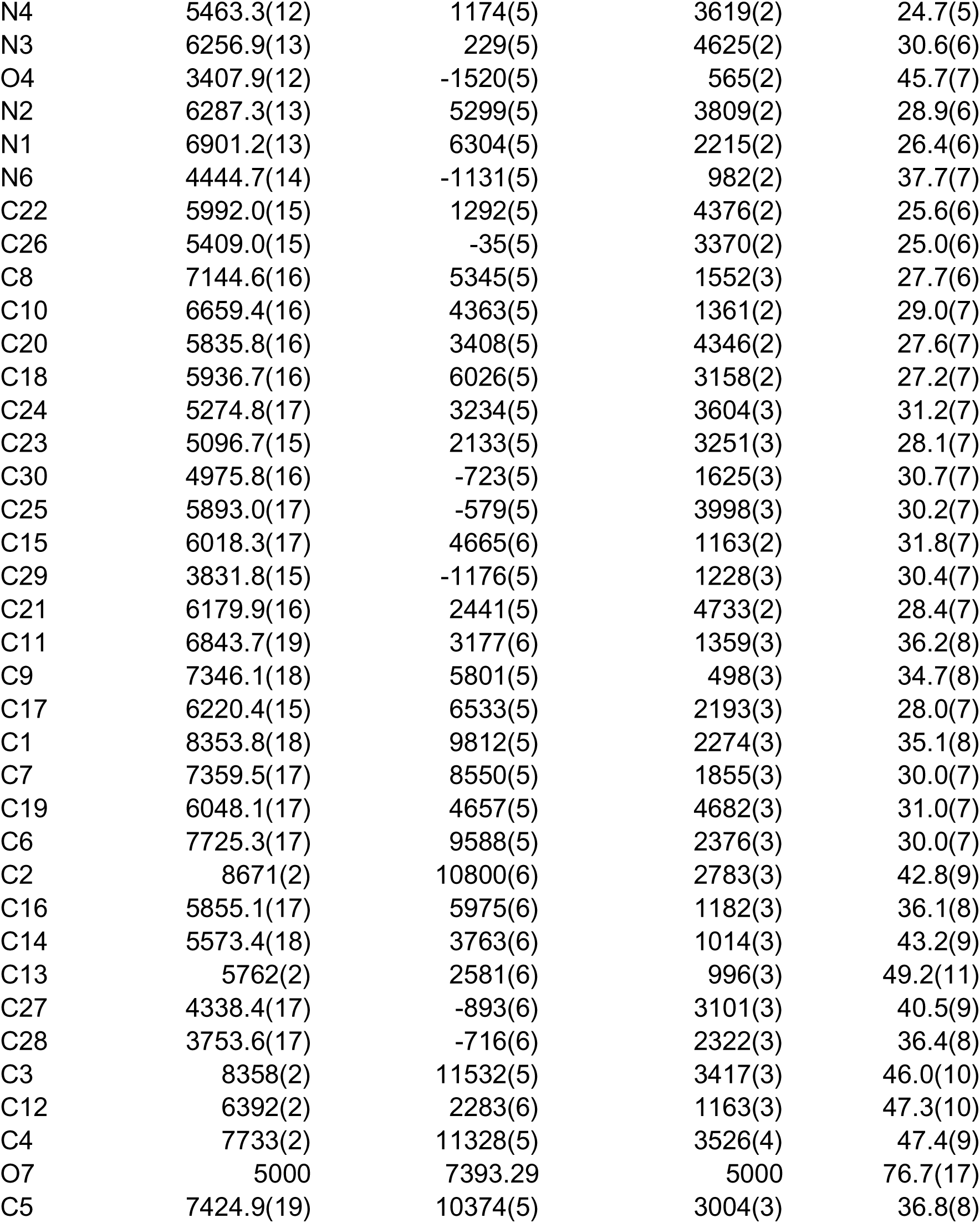
Fractional Atomic Coordinates (×10^4^) and Equivalent Isotropic Displacement Parameters (Å^2^×10^3^) for EW40755-419-P1B_auto. U_eq_ is defined as 1/3 of the trace of the orthogonalised U_IJ_ tensor.

| Atom | x | y | z | U(eq) |
| --- | --- | --- | --- | --- |
| S1 | 7388.8(4) | 7307(4) | 2775.2(6) | 24.92(19) |
| Cl1 | 8765.7(4) | 8891(4) | 1500.4(7) | 47.8(3) |
| O2 | 8010.7(11) | 6809(5) | 2877.8(18) | 30.7(5) |
| O1 | 7144.9(12) | 7726(5) | 3721.3(17) | 30.7(5) |
| O6 | 7409.0(12) | -653(5) | 5765(2) | 36.5(5) |
| O3 | 5382.4(12) | 6294(5) | 3242.3(19) | 37.3(6) |
| O5 | 5477.8(12) | -615(5) | 1260.2(19) | 45.3(7) |
| N5 | 4897.9(12) | -454(5) | 2650(2) | 27.0(6) |
| N4 | 5463.3(12) | 1174(5) | 3619(2) | 24.7(5) |
| N3 | 6256.9(13) | 229(5) | 4625(2) | 30.6(6) |
| O4 | 3407.9(12) | -1520(5) | 565(2) | 45.7(7) |
| N2 | 6287.3(13) | 5299(5) | 3809(2) | 28.9(6) |
| N1 | 6901.2(13) | 6304(5) | 2215(2) | 26.4(6) |
| N6 | 4444.7(14) | -1131(5) | 982(2) | 37.7(7) |
| C22 | 5992.0(15) | 1292(5) | 4376(2) | 25.6(6) |
| C26 | 5409.0(15) | -35(5) | 3370(2) | 25.0(6) |
| C8 | 7144.6(16) | 5345(5) | 1552(3) | 27.7(6) |
| C10 | 6659.4(16) | 4363(5) | 1361(2) | 29.0(7) |
| C20 | 5835.8(16) | 3408(5) | 4346(2) | 27.6(7) |
| C18 | 5936.7(16) | 6026(5) | 3158(2) | 27.2(7) |
| C24 | 5274.8(17) | 3234(5) | 3604(3) | 31.2(7) |
| C23 | 5096.7(15) | 2133(5) | 3251(3) | 28.1(7) |
| C30 | 4975.8(16) | -723(5) | 1625(3) | 30.7(7) |
| C25 | 5893.0(17) | -579(5) | 3998(3) | 30.2(7) |
| C15 | 6018.3(17) | 4665(6) | 1163(2) | 31.8(7) |
| C29 | 3831.8(15) | -1176(5) | 1228(3) | 30.4(7) |
| C21 | 6179.9(16) | 2441(5) | 4733(2) | 28.4(7) |
| C11 | 6843.7(19) | 3177(6) | 1359(3) | 36.2(8) |
| C9 | 7346.1(18) | 5801(5) | 498(3) | 34.7(8) |
| C17 | 6220.4(15) | 6533(5) | 2193(3) | 28.0(7) |
| C1 | 8353.8(18) | 9812(5) | 2274(3) | 35.1(8) |
| C7 | 7359.5(17) | 8550(5) | 1855(3) | 30.0(7) |
| C19 | 6048.1(17) | 4657(5) | 4682(3) | 31.0(7) |
| C6 | 7725.3(17) | 9588(5) | 2376(3) | 30.0(7) |
| C2 | 8671(2) | 10800(6) | 2783(3) | 42.8(9) |
| C16 | 5855.1(17) | 5975(6) | 1182(3) | 36.1(8) |
| C14 | 5573.4(18) | 3763(6) | 1014(3) | 43.2(9) |
| C13 | 5762(2) | 2581(6) | 996(3) | 49.2(11) |
| C27 | 4338.4(17) | -893(6) | 3101(3) | 40.5(9) |
| C28 | 3753.6(17) | -716(6) | 2322(3) | 36.4(8) |
| C3 | 8358(2) | 11532(5) | 3417(3) | 46.0(10) |
| C12 | 6392(2) | 2283(6) | 1163(3) | 47.3(10) |
| C4 | 7733(2) | 11328(5) | 3526(4) | 47.4(9) |
| O7 | 5000 | 7393.29 | 5000 | 76.7(17) |
| C5 | 7424.9(19) | 10374(5) | 3004(3) | 36.8(8) |

**Table 3.**
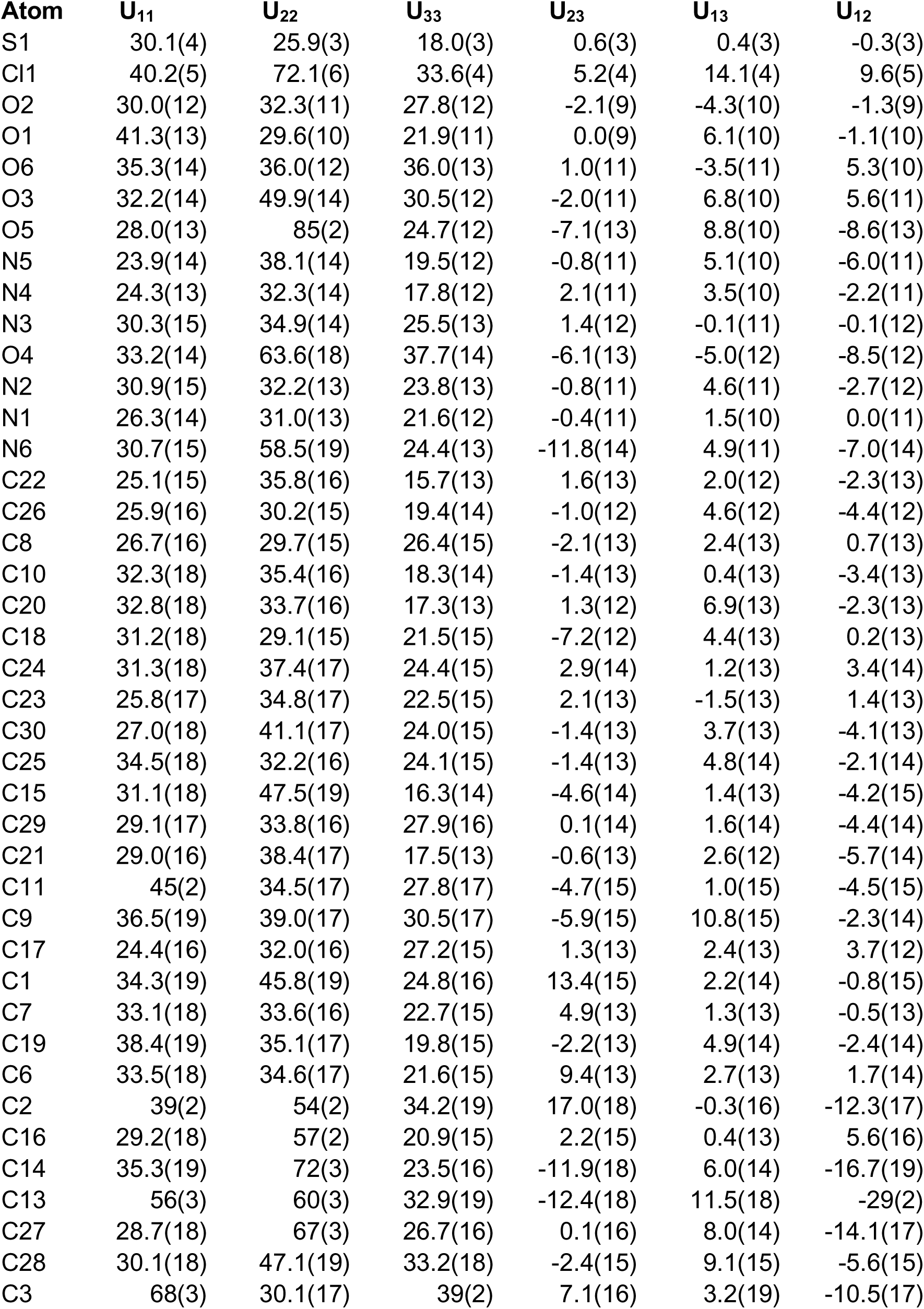
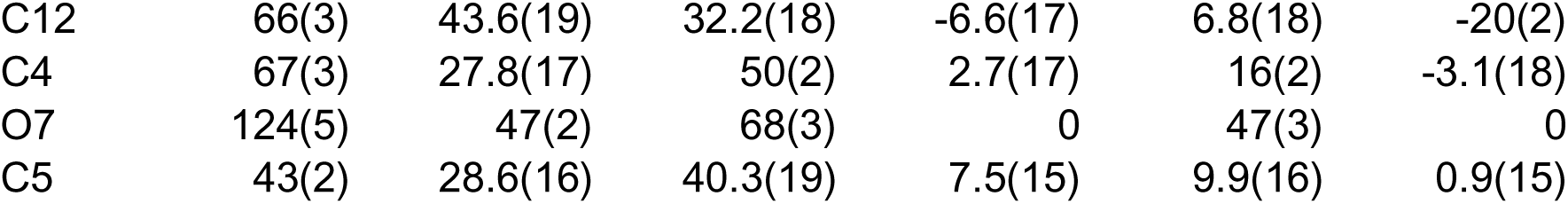
Anisotropic Displacement Parameters (Å^2^×10^3^) for EW40755-419- P1B_auto. The Anisotropic displacement factor exponent takes the form: - 2π^2^[h^2^a*^2^U_11_+2hka*b*U_12_+…].

| Atom | $U_{11}$ | $U_{22}$ | $U_{33}$ | $U_{23}$ | $U_{13}$ | $U_{12}$ |
| --- | --- | --- | --- | --- | --- | --- |
| S1 | 30.1(4) | 25.9(3) | 18.0(3) | 0.6(3) | 0.4(3) | -0.3(3) |
| Cl1 | 40.2(5) | 72.1(6) | 33.6(4) | 5.2(4) | 14.1(4) | 9.6(5) |
| O2 | 30.0(12) | 32.3(11) | 27.8(12) | -2.1(9) | -4.3(10) | -1.3(9) |
| O1 | 41.3(13) | 29.6(10) | 21.9(11) | 0.0(9) | 6.1(10) | -1.1(10) |
| O6 | 35.3(14) | 36.0(12) | 36.0(13) | 1.0(11) | -3.5(11) | 5.3(10) |
| O3 | 32.2(14) | 49.9(14) | 30.5(12) | -2.0(11) | 6.8(10) | 5.6(11) |
| O5 | 28.0(13) | 85(2) | 24.7(12) | -7.1(13) | 8.8(10) | -8.6(13) |
| N5 | 23.9(14) | 38.1(14) | 19.5(12) | -0.8(11) | 5.1(10) | -6.0(11) |
| N4 | 24.3(13) | 32.3(14) | 17.8(12) | 2.1(11) | 3.5(10) | -2.2(11) |
| N3 | 30.3(15) | 34.9(14) | 25.5(13) | 1.4(12) | -0.1(11) | -0.1(12) |
| O4 | 33.2(14) | 63.6(18) | 37.7(14) | -6.1(13) | -5.0(12) | -8.5(12) |
| N2 | 30.9(15) | 32.2(13) | 23.8(13) | -0.8(11) | 4.6(11) | -2.7(12) |
| N1 | 26.3(14) | 31.0(13) | 21.6(12) | -0.4(11) | 1.5(10) | 0.0(11) |
| N6 | 30.7(15) | 58.5(19) | 24.4(13) | -11.8(14) | 4.9(11) | -7.0(14) |
| C22 | 25.1(15) | 35.8(16) | 15.7(13) | 1.6(13) | 2.0(12) | -2.3(13) |
| C26 | 25.9(16) | 30.2(15) | 19.4(14) | -1.0(12) | 4.6(12) | -4.4(12) |
| C8 | 26.7(16) | 29.7(15) | 26.4(15) | -2.1(13) | 2.4(13) | 0.7(13) |
| C10 | 32.3(18) | 35.4(16) | 18.3(14) | -1.4(13) | 0.4(13) | -3.4(13) |
| C20 | 32.8(18) | 33.7(16) | 17.3(13) | 1.3(12) | 6.9(13) | -2.3(13) |
| C18 | 31.2(18) | 29.1(15) | 21.5(15) | -7.2(12) | 4.4(13) | 0.2(13) |
| C24 | 31.3(18) | 37.4(17) | 24.4(15) | 2.9(14) | 1.2(13) | 3.4(14) |
| C23 | 25.8(17) | 34.8(17) | 22.5(15) | 2.1(13) | -1.5(13) | 1.4(13) |
| C30 | 27.0(18) | 41.1(17) | 24.0(15) | -1.4(13) | 3.7(13) | -4.1(13) |
| C25 | 34.5(18) | 32.2(16) | 24.1(15) | -1.4(13) | 4.8(14) | -2.1(14) |
| C15 | 31.1(18) | 47.5(19) | 16.3(14) | -4.6(14) | 1.4(13) | -4.2(15) |
| C29 | 29.1(17) | 33.8(16) | 27.9(16) | 0.1(14) | 1.6(14) | -4.4(14) |
| C21 | 29.0(16) | 38.4(17) | 17.5(13) | -0.6(13) | 2.6(12) | -5.7(14) |
| C11 | 45(2) | 34.5(17) | 27.8(17) | -4.7(15) | 1.0(15) | -4.5(15) |
| C9 | 36.5(19) | 39.0(17) | 30.5(17) | -5.9(15) | 10.8(15) | -2.3(14) |
| C17 | 24.4(16) | 32.0(16) | 27.2(15) | 1.3(13) | 2.4(13) | 3.7(12) |
| C1 | 34.3(19) | 45.8(19) | 24.8(16) | 13.4(15) | 2.2(14) | -0.8(15) |
| C7 | 33.1(18) | 33.6(16) | 22.7(15) | 4.9(13) | 1.3(13) | -0.5(13) |
| C19 | 38.4(19) | 35.1(17) | 19.8(15) | -2.2(13) | 4.9(14) | -2.4(14) |
| C6 | 33.5(18) | 34.6(17) | 21.6(15) | 9.4(13) | 2.7(13) | 1.7(14) |
| C2 | 39(2) | 54(2) | 34.2(19) | 17.0(18) | -0.3(16) | -12.3(17) |
| C16 | 29.2(18) | 57(2) | 20.9(15) | 2.2(15) | 0.4(13) | 5.6(16) |
| C14 | 35.3(19) | 72(3) | 23.5(16) | -11.9(18) | 6.0(14) | -16.7(19) |
| C13 | 56(3) | 60(3) | 32.9(19) | -12.4(18) | 11.5(18) | -29(2) |
| C27 | 28.7(18) | 67(3) | 26.7(16) | 0.1(16) | 8.0(14) | -14.1(17) |
| C28 | 30.1(18) | 47.1(19) | 33.2(18) | -2.4(15) | 9.1(15) | -5.6(15) |
| C3 | 68(3) | 30.1(17) | 39(2) | 7.1(16) | 3.2(19) | -10.5(17) |
| C12 | 66(3) | 43.6(19) | 32.2(18) | -6.6(17) | 6.8(18) | -20(2) |
| C4 | 67(3) | 27.8(17) | 50(2) | 2.7(17) | 16(2) | -3.1(18) |
| O7 | 124(5) | 47(2) | 68(3) | 0 | 47(3) | 0 |
| C5 | 43(2) | 28.6(16) | 40.3(19) | 7.5(15) | 9.9(16) | 0.9(15) |

**Table 4.** Bond Lengths for EW40755-419-P1B_auto.

| Atom | Atom | Length/$\text{\AA}$ | Atom | Atom | Length/$\text{\AA}$ |
| --- | --- | --- | --- | --- | --- |
| S1 | O2 | 1.431(2) | C8 | C10 | 1.509(5) |
| S1 | O1 | 1.439(2) | C8 | C9 | 1.534(5) |
| S1 | N1 | 1.629(3) | C10 | C15 | 1.401(5) |
| S1 | C7 | 1.807(3) | C10 | C11 | 1.386(5) |
| Cl1 | C1 | 1.734(4) | C20 | C24 | 1.432(5) |
| O3 | C18 | 1.239(4) | C20 | C21 | 1.363(5) |
| O5 | C30 | 1.226(4) | C20 | C19 | 1.513(5) |
| N5 | C26 | 1.404(4) | C18 | C17 | 1.535(4) |
| N5 | C30 | 1.355(4) | C24 | C23 | 1.348(5) |
| N5 | C27 | 1.473(4) | C15 | C16 | 1.509(5) |
| N4 | C22 | 1.385(4) | C15 | C14 | 1.383(5) |
| N4 | C26 | 1.392(4) | C29 | C28 | 1.499(5) |
| N4 | C23 | 1.374(4) | C11 | C12 | 1.391(6) |
| N3 | C22 | 1.337(4) | C17 | C16 | 1.535(5) |
| N3 | C25 | 1.373(5) | C1 | C6 | 1.388(5) |
| O4 | C29 | 1.210(4) | C1 | C2 | 1.407(6) |
| N2 | C18 | 1.317(5) | C7 | C6 | 1.503(5) |
| N2 | C19 | 1.460(4) | C6 | C5 | 1.394(5) |
| N1 | C8 | 1.494(4) | C2 | C3 | 1.376(6) |
| N1 | C17 | 1.473(4) | C14 | C13 | 1.385(7) |
| N6 | C30 | 1.382(5) | C13 | C12 | 1.375(7) |
| N6 | C29 | 1.385(4) | C27 | C28 | 1.497(6) |
| C22 | C21 | 1.405(5) | C3 | C4 | 1.378(7) |
| C26 | C25 | 1.360(5) | C4 | C5 | 1.375(6) |

**Table 5.** Bond Angles for EW40755-419-P1B_auto.

| Atom | Atom | Atom | Angle/$^\circ$ | Atom | Atom | Atom | Angle/$^\circ$ |
| --- | --- | --- | --- | --- | --- | --- | --- |
| O2 | S1 | O1 | 119.02(14) | C23 | C24 | C20 | 121.0(3) |
| O2 | S1 | N1 | 107.67(14) | C24 | C23 | N4 | 118.7(3) |

| Atom | Atom | Atom | Angle/° | Atom | Atom | Atom | Angle/° |
| --- | --- | --- | --- | --- | --- | --- | --- |
| O2 | S1 | C7 | 108.19(15) | O5 | C30 | N5 | 123.8(3) |
| O1 | S1 | N1 | 107.63(14) | O5 | C30 | N6 | 120.3(3) |
| O1 | S1 | C7 | 106.99(14) | N5 | C30 | N6 | 115.9(3) |
| N1 | S1 | C7 | 106.74(15) | C26 | C25 | N3 | 111.6(3) |
| C26 | N5 | C27 | 117.7(3) | C10 | C15 | C16 | 117.0(3) |
| C30 | N5 | C26 | 120.4(3) | C14 | C15 | C10 | 119.1(4) |
| C30 | N5 | C27 | 119.9(3) | C14 | C15 | C16 | 123.8(4) |
| C22 | N4 | C26 | 106.5(3) | O4 | C29 | N6 | 120.1(3) |
| C23 | N4 | C22 | 122.3(3) | O4 | C29 | C28 | 125.3(3) |
| C23 | N4 | C26 | 131.2(3) | N6 | C29 | C28 | 114.5(3) |
| C22 | N3 | C25 | 105.2(3) | C20 | C21 | C22 | 119.7(3) |
| C18 | N2 | C19 | 123.1(3) | C10 | C11 | C12 | 120.0(4) |
| C8 | N1 | S1 | 119.2(2) | N1 | C17 | C18 | 114.3(3) |
| C17 | N1 | S1 | 117.5(2) | N1 | C17 | C16 | 109.8(3) |
| C17 | N1 | C8 | 122.0(3) | C16 | C17 | C18 | 107.1(3) |
| C30 | N6 | C29 | 127.5(3) | C6 | C1 | Cl1 | 120.8(3) |
| N4 | C22 | C21 | 118.7(3) | C6 | C1 | C2 | 120.9(4) |
| N3 | C22 | N4 | 111.0(3) | C2 | C1 | Cl1 | 118.3(3) |
| N3 | C22 | C21 | 130.3(3) | C6 | C7 | S1 | 110.2(2) |
| N4 | C26 | N5 | 120.3(3) | N2 | C19 | C20 | 111.5(3) |
| C25 | C26 | N5 | 133.9(3) | C1 | C6 | C7 | 123.6(3) |
| C25 | C26 | N4 | 105.7(3) | C1 | C6 | C5 | 117.4(3) |
| N1 | C8 | C10 | 109.3(3) | C5 | C6 | C7 | 119.0(3) |
| N1 | C8 | C9 | 113.6(3) | C3 | C2 | C1 | 119.5(4) |
| C10 | C8 | C9 | 111.7(3) | C15 | C16 | C17 | 108.4(3) |
| C15 | C10 | C8 | 119.2(3) | C15 | C14 | C13 | 120.3(4) |
| C11 | C10 | C8 | 120.6(3) | C12 | C13 | C14 | 120.7(4) |
| C11 | C10 | C15 | 120.2(3) | N5 | C27 | C28 | 110.7(3) |
| C24 | C20 | C19 | 120.2(3) | C27 | C28 | C29 | 111.8(3) |
| C21 | C20 | C24 | 119.4(3) | C2 | C3 | C4 | 120.5(4) |
| C21 | C20 | C19 | 120.4(3) | C13 | C12 | C11 | 119.6(4) |
| O3 | C18 | N2 | 124.7(3) | C5 | C4 | C3 | 119.5(4) |
| O3 | C18 | C17 | 117.3(3) | C4 | C5 | C6 | 122.2(4) |
| N2 | C18 | C17 | 118.0(3) |  |  |  |  |

**Table 6.**
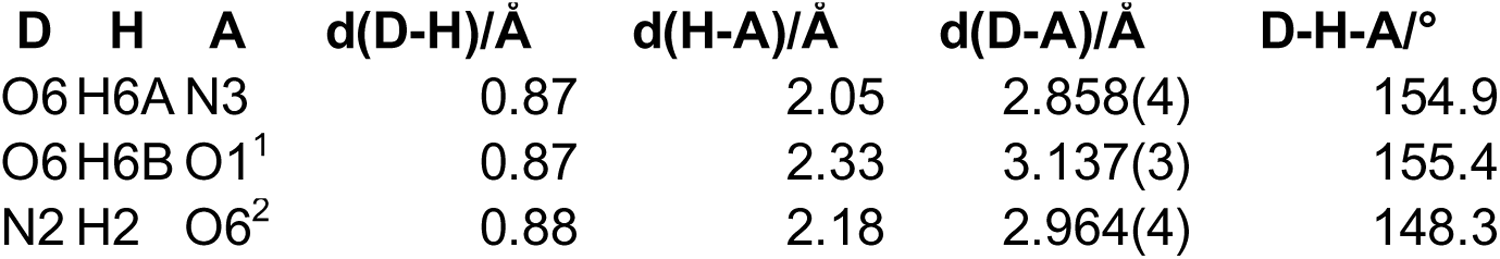
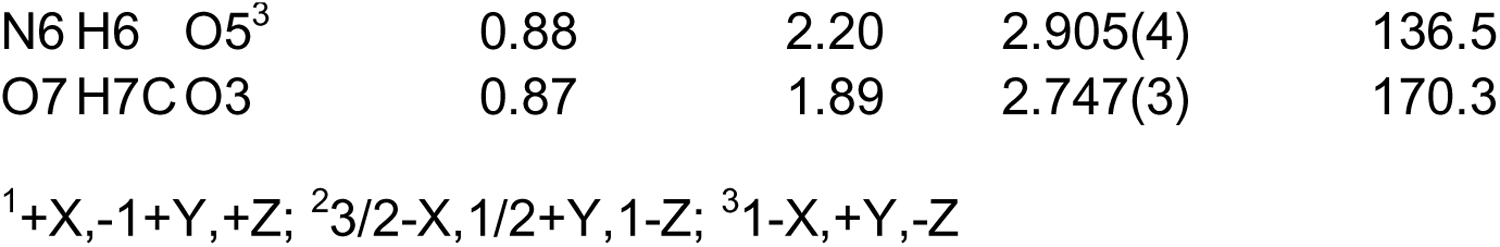
Hydrogen Bonds for EW40755-419-P1B_auto.

**Table 7.**
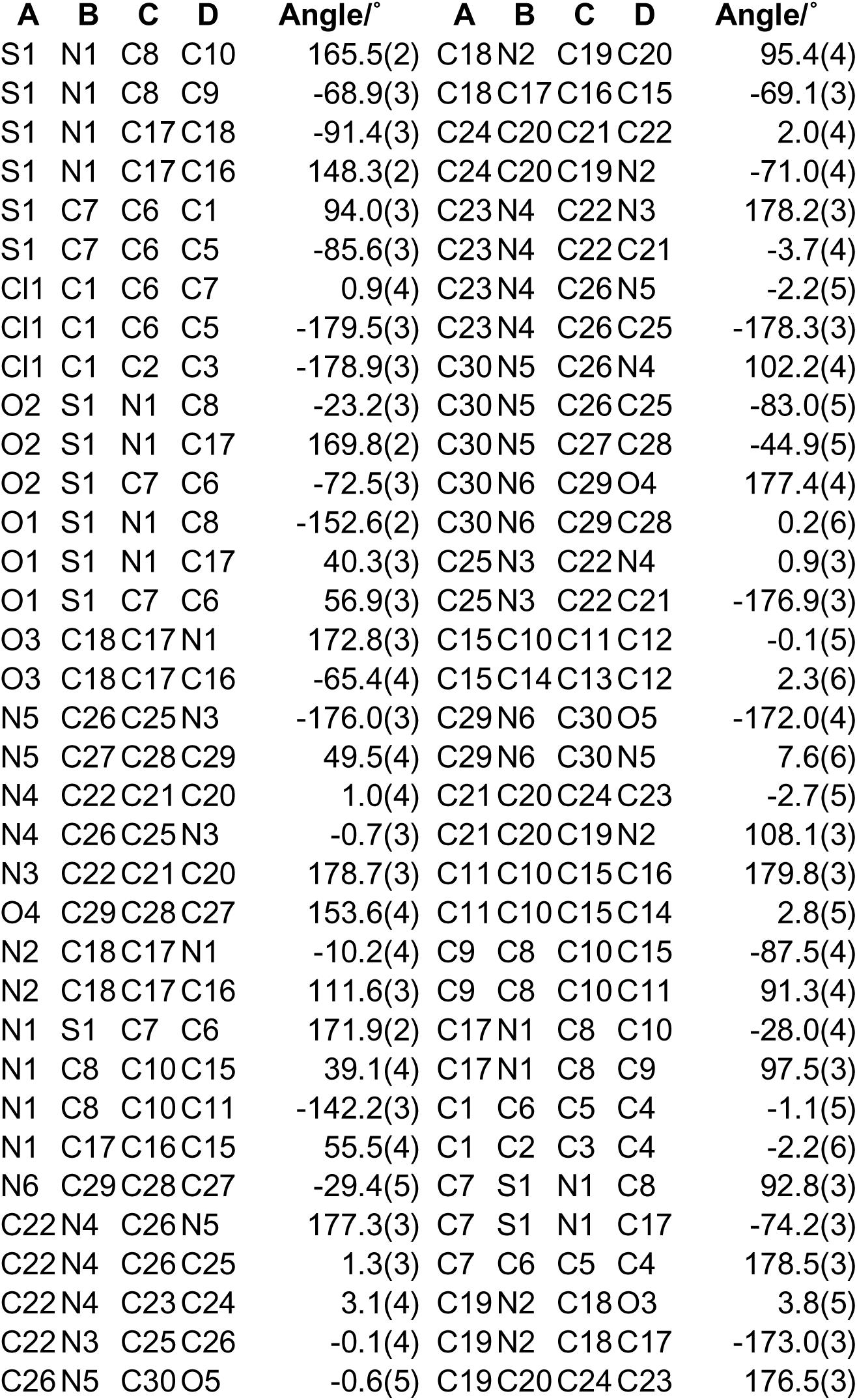
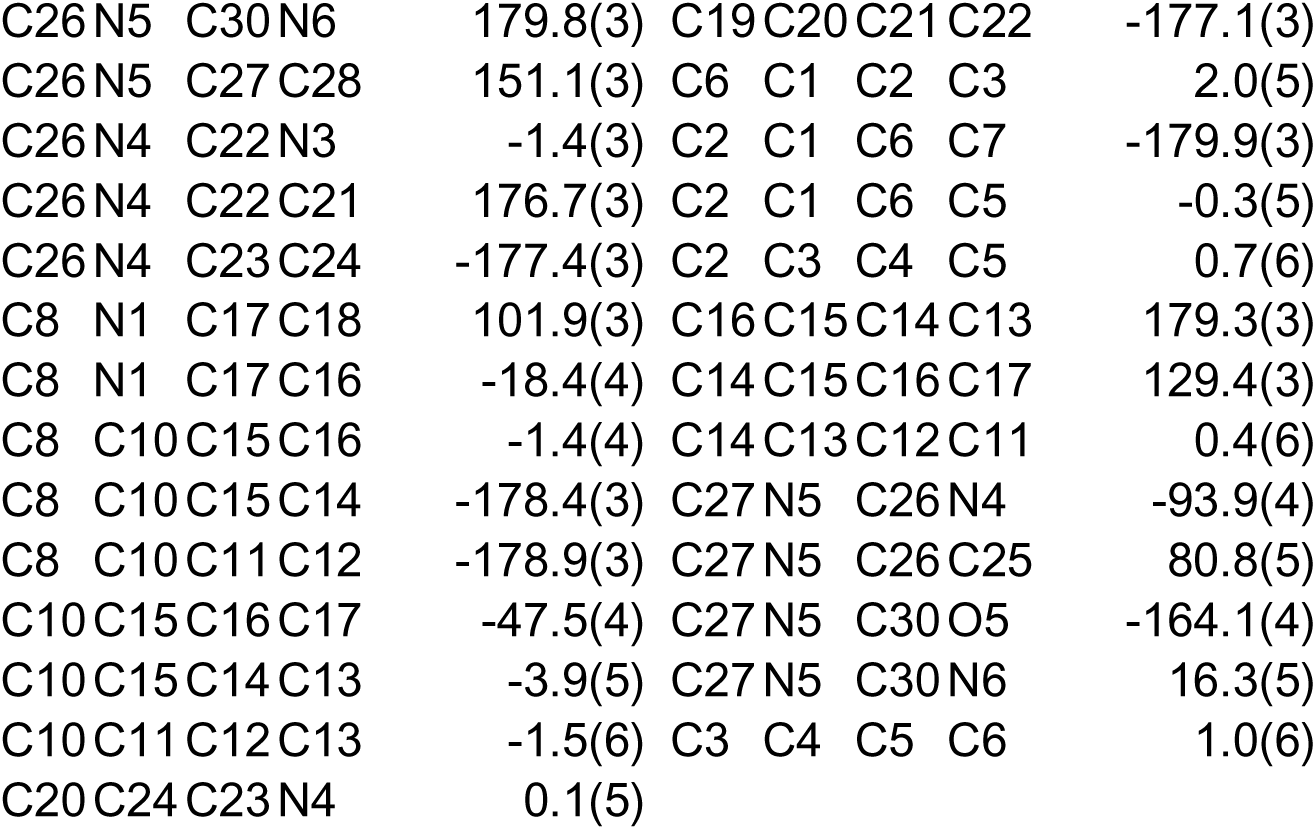
Torsion Angles for EW40755-419-P1B_auto.

**Table 8.**
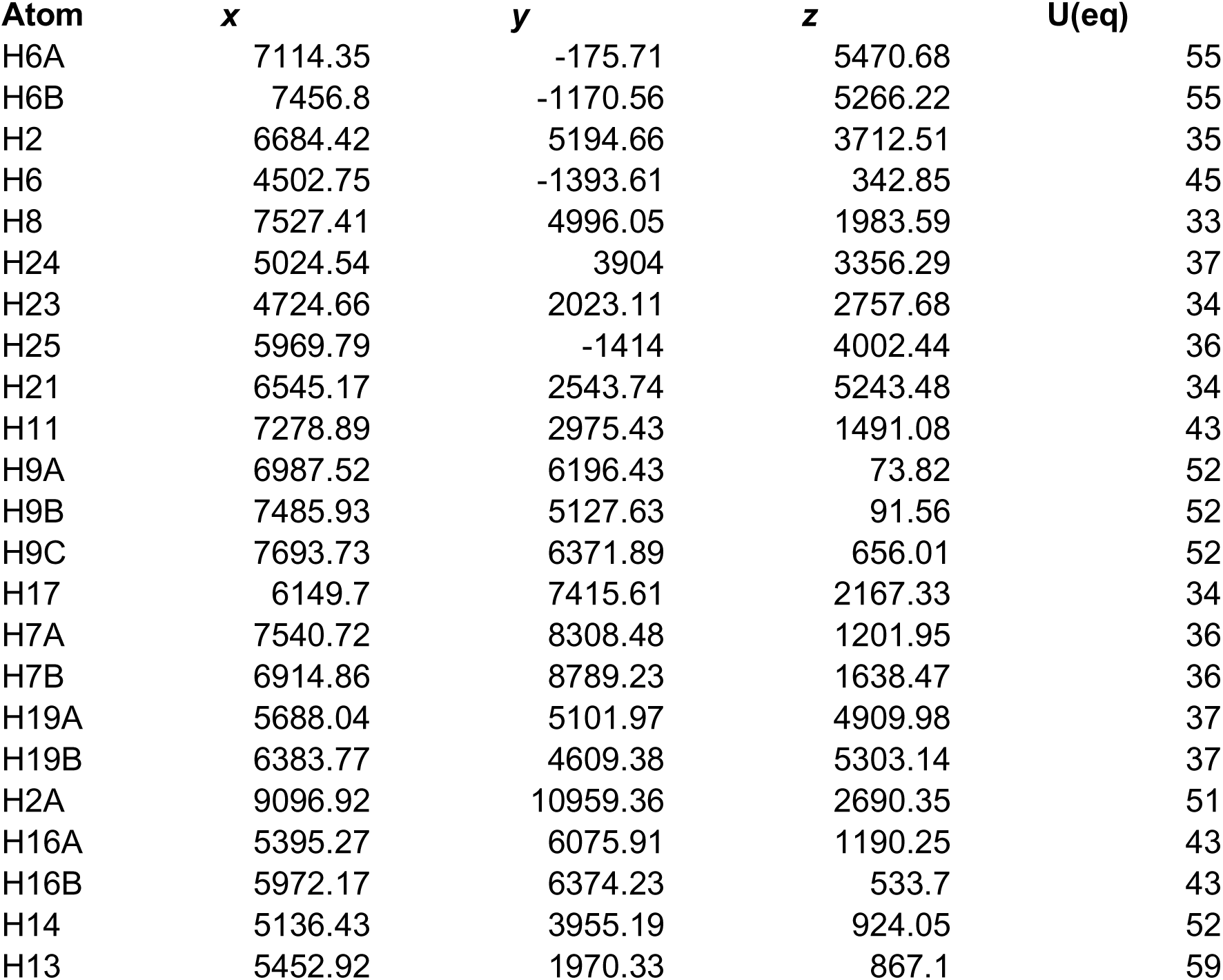
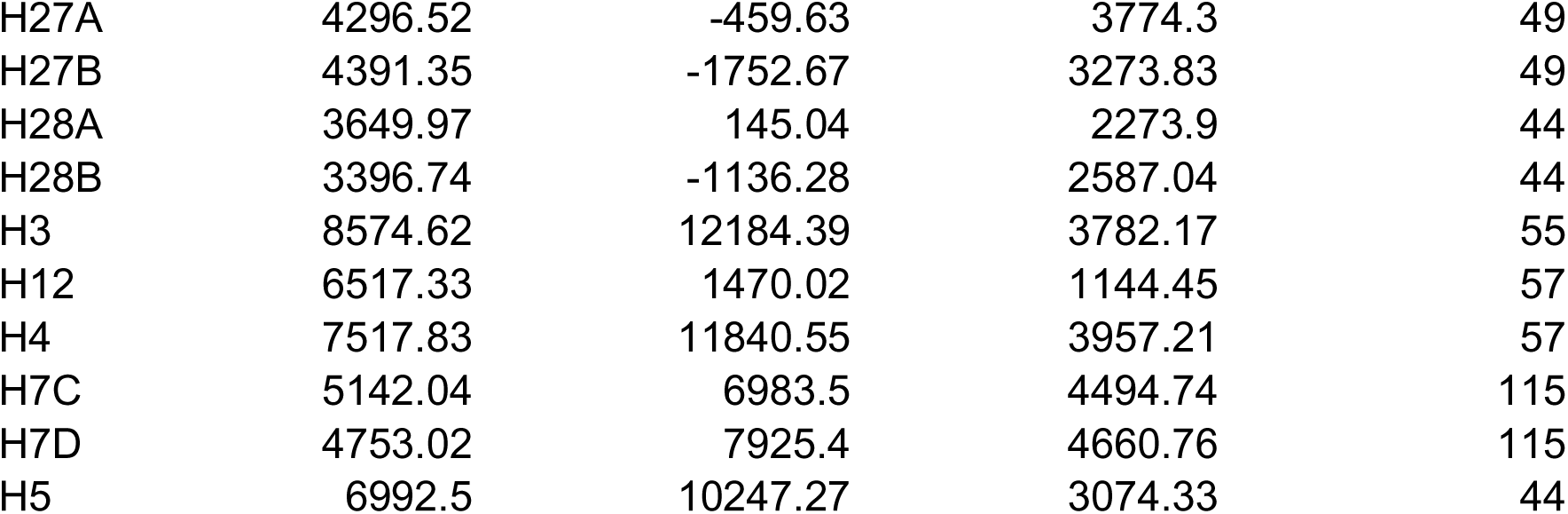
Hydrogen Atom Coordinates (Å×10^4^) and Isotropic Displacement Parameters (Å^2^×10^3^) for EW40755-419-P1B_auto.

| Atom | x | y | z | U(eq) |
| --- | --- | --- | --- | --- |
| H6A | 7114.35 | -175.71 | 5470.68 | 55 |
| H6B | 7456.8 | -1170.56 | 5266.22 | 55 |
| H2 | 6684.42 | 5194.66 | 3712.51 | 35 |
| H6 | 4502.75 | -1393.61 | 342.85 | 45 |
| H8 | 7527.41 | 4996.05 | 1983.59 | 33 |
| H24 | 5024.54 | 3904 | 3356.29 | 37 |
| H23 | 4724.66 | 2023.11 | 2757.68 | 34 |
| H25 | 5969.79 | -1414 | 4002.44 | 36 |
| H21 | 6545.17 | 2543.74 | 5243.48 | 34 |
| H11 | 7278.89 | 2975.43 | 1491.08 | 43 |
| H9A | 6987.52 | 6196.43 | 73.82 | 52 |
| H9B | 7485.93 | 5127.63 | 91.56 | 52 |
| H9C | 7693.73 | 6371.89 | 656.01 | 52 |
| H17 | 6149.7 | 7415.61 | 2167.33 | 34 |
| H7A | 7540.72 | 8308.48 | 1201.95 | 36 |
| H7B | 6914.86 | 8789.23 | 1638.47 | 36 |
| H19A | 5688.04 | 5101.97 | 4909.98 | 37 |
| H19B | 6383.77 | 4609.38 | 5303.14 | 37 |
| H2A | 9096.92 | 10959.36 | 2690.35 | 51 |
| H16A | 5395.27 | 6075.91 | 1190.25 | 43 |
| H16B | 5972.17 | 6374.23 | 533.7 | 43 |
| H14 | 5136.43 | 3955.19 | 924.05 | 52 |
| H13 | 5452.92 | 1970.33 | 867.1 | 59 |
| H27A | 4296.52 | -459.63 | 3774.3 | 49 |
| H27B | 4391.35 | -1752.67 | 3273.83 | 49 |
| H28A | 3649.97 | 145.04 | 2273.9 | 44 |
| H28B | 3396.74 | -1136.28 | 2587.04 | 44 |
| H3 | 8574.62 | 12184.39 | 3782.17 | 55 |
| H12 | 6517.33 | 1470.02 | 1144.45 | 57 |
| H4 | 7517.83 | 11840.55 | 3957.21 | 57 |
| H7C | 5142.04 | 6983.5 | 4494.74 | 115 |
| H7D | 4753.02 | 7925.4 | 4660.76 | 115 |
| H5 | 6992.5 | 10247.27 | 3074.33 | 44 |

**Table 9.** Atomic Occupancy for EW40755-419-P1B_auto.

| Atom | Occupancy | Atom | Occupancy | Atom | Occupancy |
| --- | --- | --- | --- | --- | --- |
| H7C | 0.5 | H7D | 0.5 |  |  |

**Table 1.** Crystal data and structure refinement.

|  |  |
| --- | --- |
| Empirical formula | C<sub>30</sub>H<sub>32</sub>ClN<sub>6</sub>O<sub>6.5</sub>S |
| Formula weight | 648.12 |
| Temperature/K | 150.00(10) |
| Crystal system | monoclinic |
| Space group | C2 |
| a/Å | 21.3441(3) |
| b/Å | 11.1992(2) |
| c/Å | 12.5765(2) |
| α/° | 90 |
| β/° | 97.5310(10) |
| γ/° | 90 |
| Volume/Å<sup>3</sup> | 2980.31(8) |
| Z | 4 |
| ρ<sub>calc</sub>/g/cm<sup>3</sup> | 1.444 |
| μ/mm<sup>-1</sup> | 2.274 |
| F(000) | 1356.0 |
| Crystal size/mm<sup>3</sup> | 0.25 × 0.1 × 0.1 |
| Radiation | Cu Kα (λ = 1.54184) |
| 2θ range for data collection/° | 7.09 to 133.186 |
| Index ranges | -25 ≤ h ≤ 25, -13 ≤ k ≤ 13, -14 ≤ l ≤ 14 |
| Reflections collected | 18726 |
| Independent reflections | 5178 [R<sub>int</sub> = 0.0453, R<sub>sigma</sub> = 0.0382] |
| Data/restraints/parameters | 5178/2/408 |
| Goodness-of-fit on F<sup>2</sup> | 1.047 |
| Final R indexes [I ≥ 2σ (I)] | R<sub>1</sub> = 0.0331, wR<sub>2</sub> = 0.0847 |
| Final R indexes [all data] | R<sub>1</sub> = 0.0342, wR<sub>2</sub> = 0.0856 |
| Largest diff. peak/hole / e Å<sup>-3</sup> | 0.35/-0.32 |
| Flack parameter | -0.011(8) |

**Table 2.**
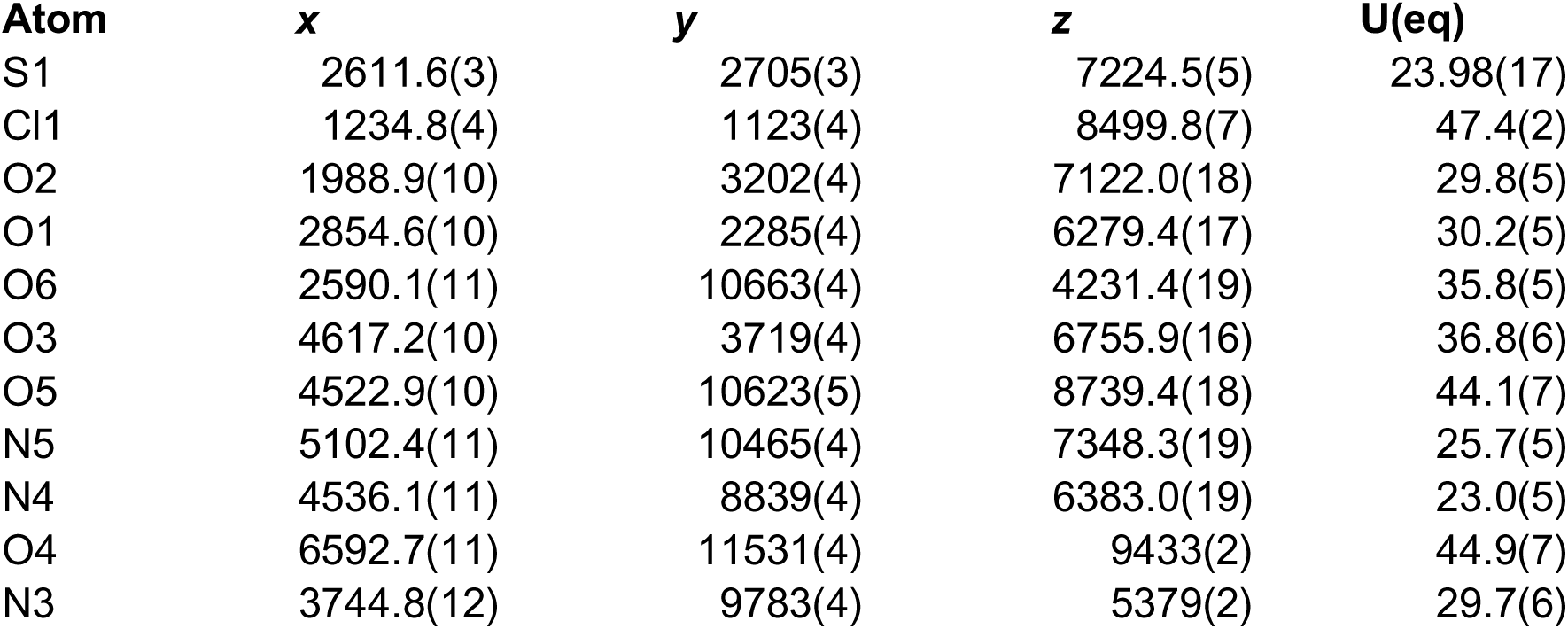
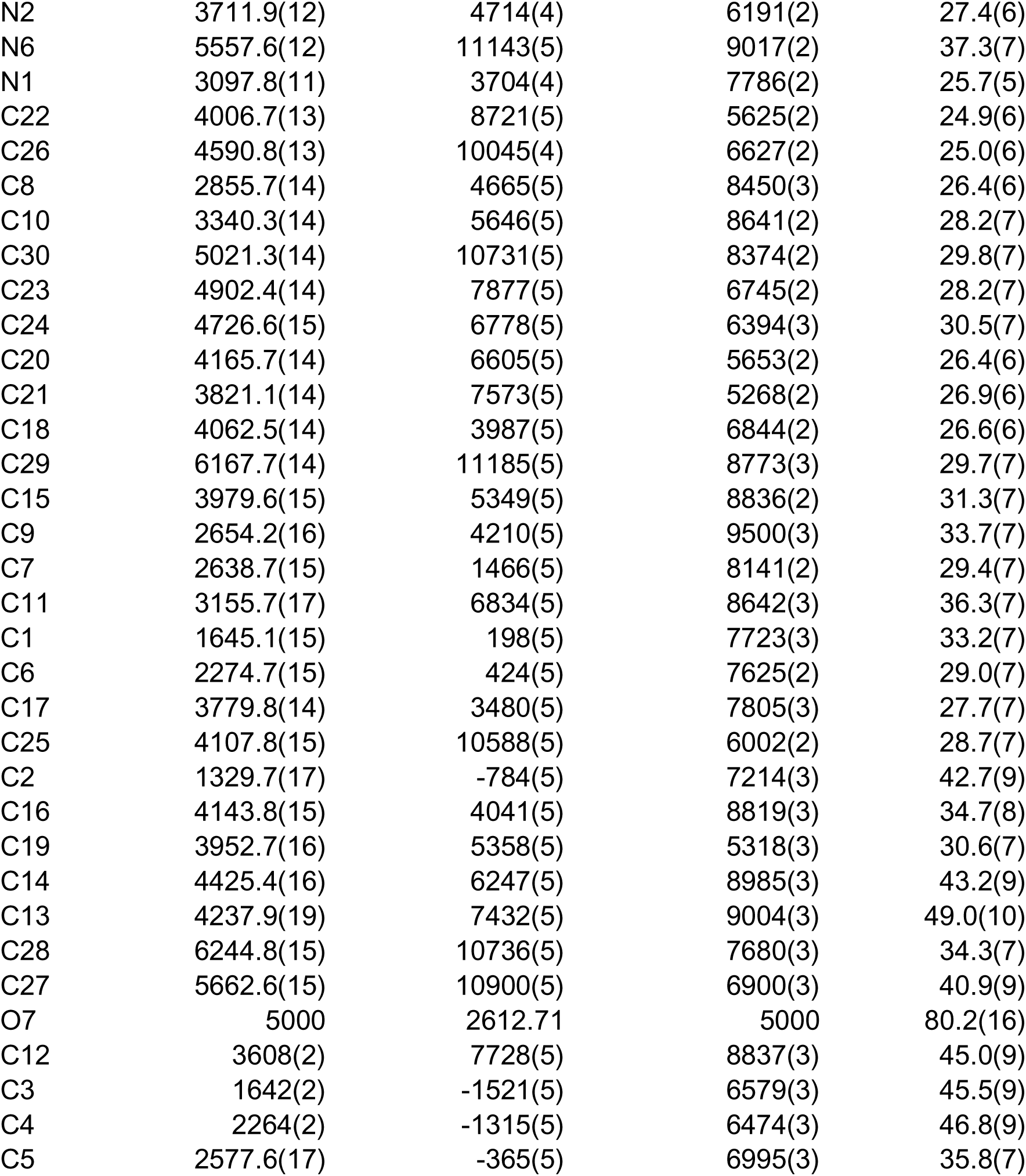
Fractional Atomic Coordinates (×10^4^) and Equivalent Isotropic Displacement Parameters (Å^2^×10^3^) for EW40755-419-P3B_auto. U_eq_ is defined as 1/3 of the trace of the orthogonalised U_IJ_ tensor.

| Atom | x | y | z | U(eq) |
| --- | --- | --- | --- | --- |
| S1 | 2611.6(3) | 2705(3) | 7224.5(5) | 23.98(17) |
| Cl1 | 1234.8(4) | 1123(4) | 8499.8(7) | 47.4(2) |
| O2 | 1988.9(10) | 3202(4) | 7122.0(18) | 29.8(5) |
| O1 | 2854.6(10) | 2285(4) | 6279.4(17) | 30.2(5) |
| O6 | 2590.1(11) | 10663(4) | 4231.4(19) | 35.8(5) |
| O3 | 4617.2(10) | 3719(4) | 6755.9(16) | 36.8(6) |
| O5 | 4522.9(10) | 10623(5) | 8739.4(18) | 44.1(7) |
| N5 | 5102.4(11) | 10465(4) | 7348.3(19) | 25.7(5) |
| N4 | 4536.1(11) | 8839(4) | 6383.0(19) | 23.0(5) |
| O4 | 6592.7(11) | 11531(4) | 9433(2) | 44.9(7) |
| N3 | 3744.8(12) | 9783(4) | 5379(2) | 29.7(6) |
| N2 | 3711.9(12) | 4714(4) | 6191(2) | 27.4(6) |
| N6 | 5557.6(12) | 11143(5) | 9017(2) | 37.3(7) |
| N1 | 3097.8(11) | 3704(4) | 7786(2) | 25.7(5) |
| C22 | 4006.7(13) | 8721(5) | 5625(2) | 24.9(6) |
| C26 | 4590.8(13) | 10045(4) | 6627(2) | 25.0(6) |
| C8 | 2855.7(14) | 4665(5) | 8450(3) | 26.4(6) |
| C10 | 3340.3(14) | 5646(5) | 8641(2) | 28.2(7) |
| C30 | 5021.3(14) | 10731(5) | 8374(2) | 29.8(7) |
| C23 | 4902.4(14) | 7877(5) | 6745(2) | 28.2(7) |
| C24 | 4726.6(15) | 6778(5) | 6394(3) | 30.5(7) |
| C20 | 4165.7(14) | 6605(5) | 5653(2) | 26.4(6) |
| C21 | 3821.1(14) | 7573(5) | 5268(2) | 26.9(6) |
| C18 | 4062.5(14) | 3987(5) | 6844(2) | 26.6(6) |
| C29 | 6167.7(14) | 11185(5) | 8773(3) | 29.7(7) |
| C15 | 3979.6(15) | 5349(5) | 8836(2) | 31.3(7) |
| C9 | 2654.2(16) | 4210(5) | 9500(3) | 33.7(7) |
| C7 | 2638.7(15) | 1466(5) | 8141(2) | 29.4(7) |
| C11 | 3155.7(17) | 6834(5) | 8642(3) | 36.3(7) |
| C1 | 1645.1(15) | 198(5) | 7723(3) | 33.2(7) |
| C6 | 2274.7(15) | 424(5) | 7625(2) | 29.0(7) |
| C17 | 3779.8(14) | 3480(5) | 7805(3) | 27.7(7) |
| C25 | 4107.8(15) | 10588(5) | 6002(2) | 28.7(7) |
| C2 | 1329.7(17) | -784(5) | 7214(3) | 42.7(9) |
| C16 | 4143.8(15) | 4041(5) | 8819(3) | 34.7(8) |
| C19 | 3952.7(16) | 5358(5) | 5318(3) | 30.6(7) |
| C14 | 4425.4(16) | 6247(5) | 8985(3) | 43.2(9) |
| C13 | 4237.9(19) | 7432(5) | 9004(3) | 49.0(10) |
| C28 | 6244.8(15) | 10736(5) | 7680(3) | 34.3(7) |
| C27 | 5662.6(15) | 10900(5) | 6900(3) | 40.9(9) |
| O7 | 5000 | 2612.71 | 5000 | 80.2(16) |
| C12 | 3608(2) | 7728(5) | 8837(3) | 45.0(9) |
| C3 | 1642(2) | -1521(5) | 6579(3) | 45.5(9) |
| C4 | 2264(2) | -1315(5) | 6474(3) | 46.8(9) |
| C5 | 2577.6(17) | -365(5) | 6995(3) | 35.8(7) |

**Table 3.**
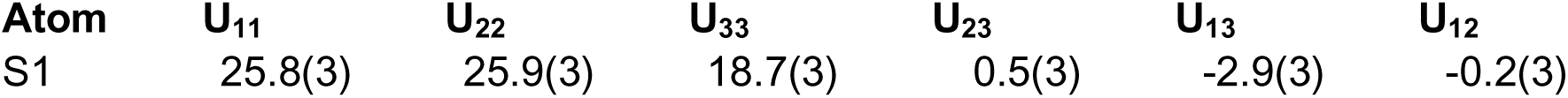
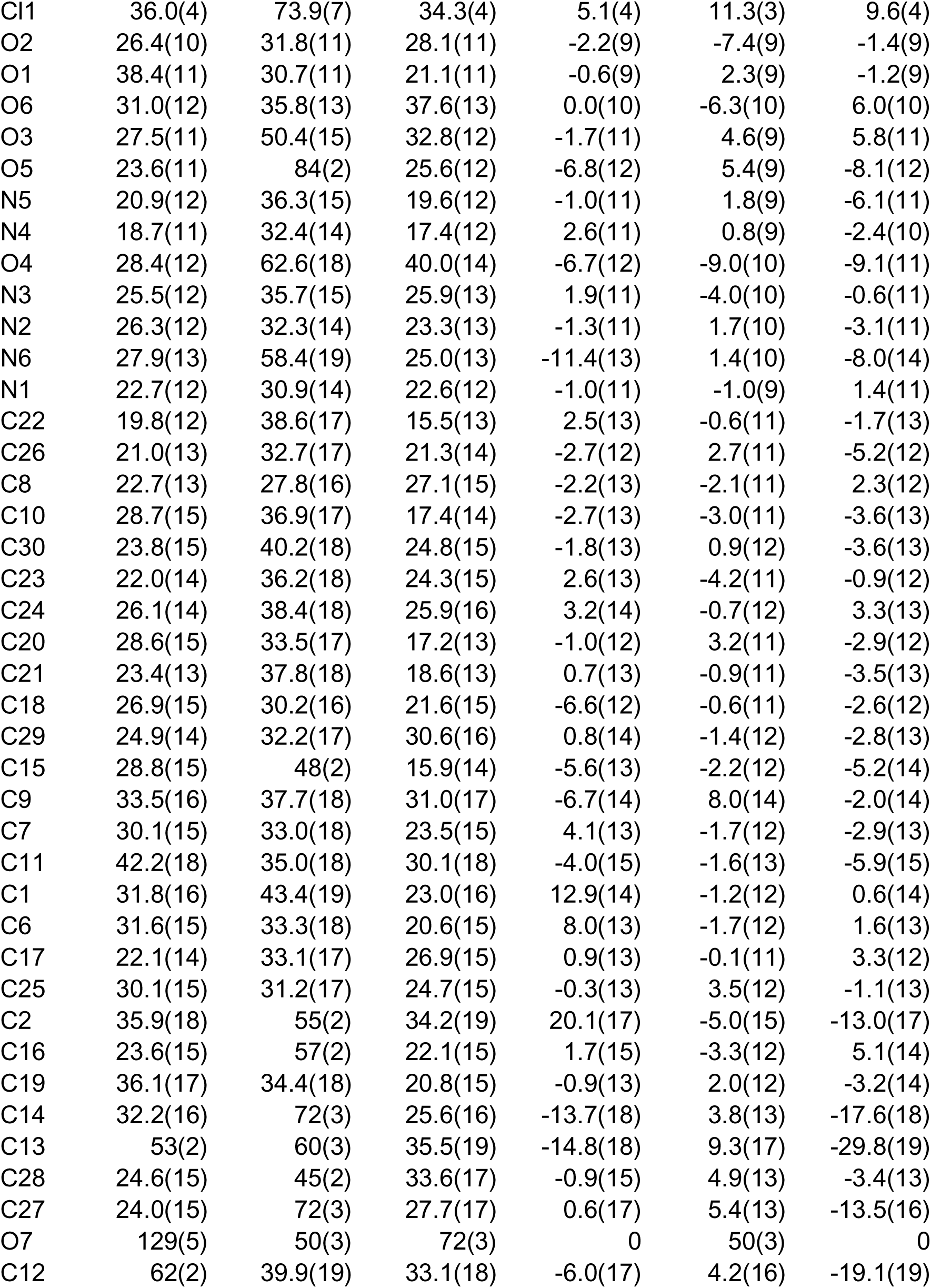
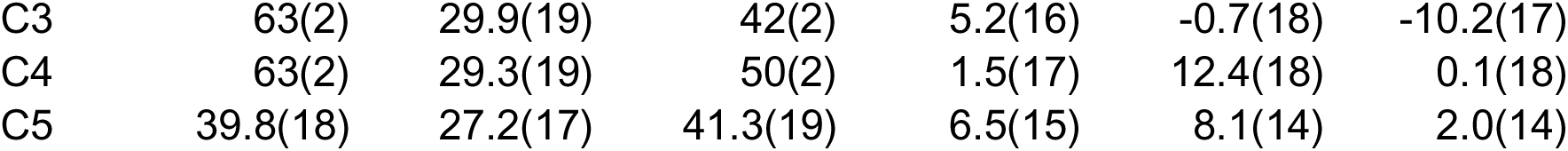
Anisotropic Displacement Parameters (Å^2^×10^3^) for EW40755-419- P3B_auto. The Anisotropic displacement factor exponent takes the form: - 2π^2^[h^2^a*^2^U_11_+2hka*b*U_12_+…].

| Atom | $U_{11}$ | $U_{22}$ | $U_{33}$ | $U_{23}$ | $U_{13}$ | $U_{12}$ |
| --- | --- | --- | --- | --- | --- | --- |
| S1 | 25.8(3) | 25.9(3) | 18.7(3) | 0.5(3) | -2.9(3) | -0.2(3) |
| Cl1 | 36.0(4) | 73.9(7) | 34.3(4) | 5.1(4) | 11.3(3) | 9.6(4) |
| O2 | 26.4(10) | 31.8(11) | 28.1(11) | -2.2(9) | -7.4(9) | -1.4(9) |
| O1 | 38.4(11) | 30.7(11) | 21.1(11) | -0.6(9) | 2.3(9) | -1.2(9) |
| O6 | 31.0(12) | 35.8(13) | 37.6(13) | 0.0(10) | -6.3(10) | 6.0(10) |
| O3 | 27.5(11) | 50.4(15) | 32.8(12) | -1.7(11) | 4.6(9) | 5.8(11) |
| O5 | 23.6(11) | 84(2) | 25.6(12) | -6.8(12) | 5.4(9) | -8.1(12) |
| N5 | 20.9(12) | 36.3(15) | 19.6(12) | -1.0(11) | 1.8(9) | -6.1(11) |
| N4 | 18.7(11) | 32.4(14) | 17.4(12) | 2.6(11) | 0.8(9) | -2.4(10) |
| O4 | 28.4(12) | 62.6(18) | 40.0(14) | -6.7(12) | -9.0(10) | -9.1(11) |
| N3 | 25.5(12) | 35.7(15) | 25.9(13) | 1.9(11) | -4.0(10) | -0.6(11) |
| N2 | 26.3(12) | 32.3(14) | 23.3(13) | -1.3(11) | 1.7(10) | -3.1(11) |
| N6 | 27.9(13) | 58.4(19) | 25.0(13) | -11.4(13) | 1.4(10) | -8.0(14) |
| N1 | 22.7(12) | 30.9(14) | 22.6(12) | -1.0(11) | -1.0(9) | 1.4(11) |
| C22 | 19.8(12) | 38.6(17) | 15.5(13) | 2.5(13) | -0.6(11) | -1.7(13) |
| C26 | 21.0(13) | 32.7(17) | 21.3(14) | -2.7(12) | 2.7(11) | -5.2(12) |
| C8 | 22.7(13) | 27.8(16) | 27.1(15) | -2.2(13) | -2.1(11) | 2.3(12) |
| C10 | 28.7(15) | 36.9(17) | 17.4(14) | -2.7(13) | -3.0(11) | -3.6(13) |
| C30 | 23.8(15) | 40.2(18) | 24.8(15) | -1.8(13) | 0.9(12) | -3.6(13) |
| C23 | 22.0(14) | 36.2(18) | 24.3(15) | 2.6(13) | -4.2(11) | -0.9(12) |
| C24 | 26.1(14) | 38.4(18) | 25.9(16) | 3.2(14) | -0.7(12) | 3.3(13) |
| C20 | 28.6(15) | 33.5(17) | 17.2(13) | -1.0(12) | 3.2(11) | -2.9(12) |
| C21 | 23.4(13) | 37.8(18) | 18.6(13) | 0.7(13) | -0.9(11) | -3.5(13) |
| C18 | 26.9(15) | 30.2(16) | 21.6(15) | -6.6(12) | -0.6(11) | -2.6(12) |
| C29 | 24.9(14) | 32.2(17) | 30.6(16) | 0.8(14) | -1.4(12) | -2.8(13) |
| C15 | 28.8(15) | 48(2) | 15.9(14) | -5.6(13) | -2.2(12) | -5.2(14) |
| C9 | 33.5(16) | 37.7(18) | 31.0(17) | -6.7(14) | 8.0(14) | -2.0(14) |
| C7 | 30.1(15) | 33.0(18) | 23.5(15) | 4.1(13) | -1.7(12) | -2.9(13) |
| C11 | 42.2(18) | 35.0(18) | 30.1(18) | -4.0(15) | -1.6(13) | -5.9(15) |
| C1 | 31.8(16) | 43.4(19) | 23.0(16) | 12.9(14) | -1.2(12) | 0.6(14) |
| C6 | 31.6(15) | 33.3(18) | 20.6(15) | 8.0(13) | -1.7(12) | 1.6(13) |
| C17 | 22.1(14) | 33.1(17) | 26.9(15) | 0.9(13) | -0.1(11) | 3.3(12) |
| C25 | 30.1(15) | 31.2(17) | 24.7(15) | -0.3(13) | 3.5(12) | -1.1(13) |
| C2 | 35.9(18) | 55(2) | 34.2(19) | 20.1(17) | -5.0(15) | -13.0(17) |
| C16 | 23.6(15) | 57(2) | 22.1(15) | 1.7(15) | -3.3(12) | 5.1(14) |
| C19 | 36.1(17) | 34.4(18) | 20.8(15) | -0.9(13) | 2.0(12) | -3.2(14) |
| C14 | 32.2(16) | 72(3) | 25.6(16) | -13.7(18) | 3.8(13) | -17.6(18) |
| C13 | 53(2) | 60(3) | 35.5(19) | -14.8(18) | 9.3(17) | -29.8(19) |
| C28 | 24.6(15) | 45(2) | 33.6(17) | -0.9(15) | 4.9(13) | -3.4(13) |
| C27 | 24.0(15) | 72(3) | 27.7(17) | 0.6(17) | 5.4(13) | -13.5(16) |
| O7 | 129(5) | 50(3) | 72(3) | 0 | 50(3) | 0 |
| C12 | 62(2) | 39.9(19) | 33.1(18) | -6.0(17) | 4.2(16) | -19.1(19) |
| C3 | 63(2) | 29.9(19) | 42(2) | 5.2(16) | -0.7(18) | -10.2(17) |
| C4 | 63(2) | 29.3(19) | 50(2) | 1.5(17) | 12.4(18) | 0.1(18) |
| C5 | 39.8(18) | 27.2(17) | 41.3(19) | 6.5(15) | 8.1(14) | 2.0(14) |

**Table 4.** Bond Lengths for EW40755-419-P3B_auto.

| Atom | Atom | Length/$\text{\AA}$ | Atom | Atom | Length/$\text{\AA}$ |
| --- | --- | --- | --- | --- | --- |
| S1 | O2 | 1.431(2) | C8 | C10 | 1.507(4) |
| S1 | O1 | 1.437(2) | C8 | C9 | 1.528(5) |
| S1 | N1 | 1.624(3) | C10 | C15 | 1.395(4) |
| S1 | C7 | 1.801(3) | C10 | C11 | 1.388(5) |
| Cl1 | C1 | 1.738(4) | C23 | C24 | 1.344(5) |
| O3 | C18 | 1.240(4) | C24 | C20 | 1.431(4) |
| O5 | C30 | 1.219(4) | C20 | C21 | 1.364(5) |
| N5 | C26 | 1.406(4) | C20 | C19 | 1.511(5) |
| N5 | C30 | 1.357(4) | C18 | C17 | 1.530(4) |
| N5 | C27 | 1.471(4) | C29 | C28 | 1.494(5) |
| N4 | C22 | 1.386(4) | C15 | C16 | 1.507(5) |
| N4 | C26 | 1.387(4) | C15 | C14 | 1.380(5) |
| N4 | C23 | 1.374(4) | C7 | C6 | 1.501(5) |
| O4 | C29 | 1.210(4) | C11 | C12 | 1.391(5) |
| N3 | C22 | 1.332(4) | C1 | C6 | 1.389(5) |
| N3 | C25 | 1.367(4) | C1 | C2 | 1.399(6) |
| N2 | C18 | 1.318(4) | C6 | C5 | 1.400(5) |
| N2 | C19 | 1.462(4) | C17 | C16 | 1.537(5) |
| N6 | C30 | 1.391(4) | C2 | C3 | 1.379(6) |
| N6 | C29 | 1.377(4) | C14 | C13 | 1.387(7) |
| N1 | C8 | 1.495(4) | C13 | C12 | 1.374(6) |
| N1 | C17 | 1.474(4) | C28 | C27 | 1.490(5) |
| C22 | C21 | 1.402(5) | C3 | C4 | 1.372(6) |
| C26 | C25 | 1.355(4) | C4 | C5 | 1.377(6) |

**Table 5.** Bond Angles for EW40755-419-P3B_auto.

| Atom | Atom | Atom | Angle/$^\circ$ | Atom | Atom | Atom | Angle/$^\circ$ |
| --- | --- | --- | --- | --- | --- | --- | --- |
| O2 | S1 | O1 | 118.99(13) | C21 | C20 | C24 | 119.3(3) |
| O2 | S1 | N1 | 107.70(13) | C21 | C20 | C19 | 120.5(3) |
| O2 | S1 | C7 | 108.01(15) | C20 | C21 | C22 | 119.9(3) |

| Atom | Atom | Atom | Angle/° | Atom | Atom | Atom | Angle/° |
| --- | --- | --- | --- | --- | --- | --- | --- |
| O1 | S1 | N1 | 107.82(13) | O3 | C18 | N2 | 124.4(3) |
| O1 | S1 | C7 | 107.00(14) | O3 | C18 | C17 | 117.4(3) |
| N1 | S1 | C7 | 106.72(14) | N2 | C18 | C17 | 118.1(3) |
| C26 | N5 | C27 | 117.7(3) | O4 | C29 | N6 | 120.3(3) |
| C30 | N5 | C26 | 120.1(2) | O4 | C29 | C28 | 125.2(3) |
| C30 | N5 | C27 | 120.2(3) | N6 | C29 | C28 | 114.5(3) |
| C22 | N4 | C26 | 106.4(2) | C10 | C15 | C16 | 116.9(3) |
| C23 | N4 | C22 | 122.1(3) | C14 | C15 | C10 | 119.4(4) |
| C23 | N4 | C26 | 131.5(3) | C14 | C15 | C16 | 123.6(3) |
| C22 | N3 | C25 | 105.4(2) | C6 | C7 | S1 | 110.5(2) |
| C18 | N2 | C19 | 123.1(3) | C10 | C11 | C12 | 120.0(4) |
| C29 | N6 | C30 | 127.7(3) | C6 | C1 | Cl1 | 120.4(3) |
| C8 | N1 | S1 | 119.4(2) | C6 | C1 | C2 | 121.1(3) |
| C17 | N1 | S1 | 117.6(2) | C2 | C1 | Cl1 | 118.5(3) |
| C17 | N1 | C8 | 121.8(2) | C1 | C6 | C7 | 123.7(3) |
| N4 | C22 | C21 | 118.6(3) | C1 | C6 | C5 | 117.3(3) |
| N3 | C22 | N4 | 110.8(3) | C5 | C6 | C7 | 118.9(3) |
| N3 | C22 | C21 | 130.6(3) | N1 | C17 | C18 | 114.5(2) |
| N4 | C26 | N5 | 120.3(3) | N1 | C17 | C16 | 109.5(3) |
| C25 | C26 | N5 | 133.8(3) | C18 | C17 | C16 | 107.2(3) |
| C25 | C26 | N4 | 105.8(3) | C26 | C25 | N3 | 111.6(3) |
| N1 | C8 | C10 | 109.5(2) | C3 | C2 | C1 | 119.6(3) |
| N1 | C8 | C9 | 113.6(3) | C15 | C16 | C17 | 108.4(3) |
| C10 | C8 | C9 | 111.8(3) | N2 | C19 | C20 | 111.7(3) |
| C15 | C10 | C8 | 119.3(3) | C15 | C14 | C13 | 120.3(3) |
| C11 | C10 | C8 | 120.7(3) | C12 | C13 | C14 | 120.5(3) |
| C11 | C10 | C15 | 120.0(3) | C27 | C28 | C29 | 112.4(3) |
| O5 | C30 | N5 | 124.3(3) | N5 | C27 | C28 | 110.9(3) |
| O5 | C30 | N6 | 120.4(3) | C13 | C12 | C11 | 119.7(4) |
| N5 | C30 | N6 | 115.3(3) | C4 | C3 | C2 | 120.2(4) |
| C24 | C23 | N4 | 119.1(3) | C3 | C4 | C5 | 120.0(4) |
| C23 | C24 | C20 | 120.8(3) | C4 | C5 | C6 | 121.7(3) |
| C24 | C20 | C19 | 120.2(3) |  |  |  |  |

**Table 6.**
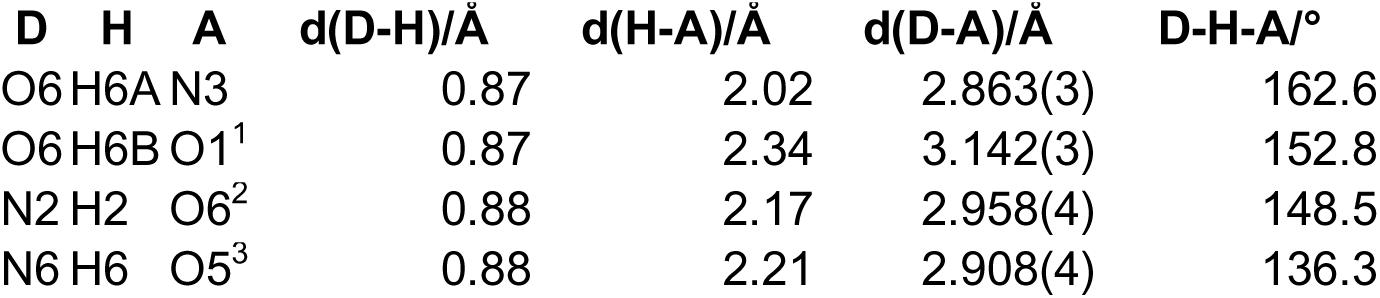
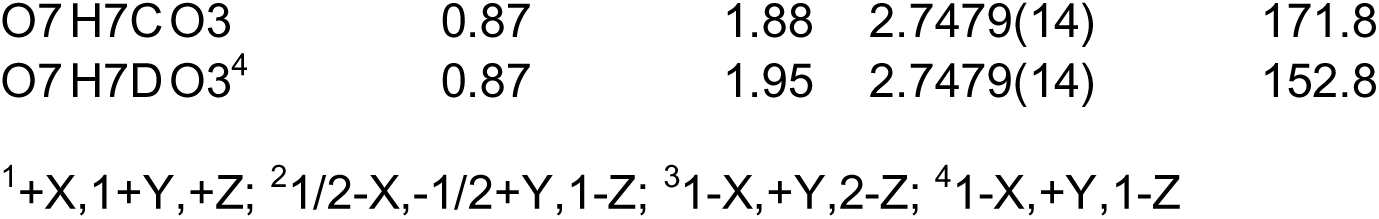
Hydrogen Bonds for EW40755-419-P3B_auto.

| D | H | A | d(D-H)/Å | d(H-A)/Å | d(D-A)/Å | D-H-A/° |
| --- | --- | --- | --- | --- | --- | --- |
| O6 | H6A | N3 | 0.87 | 2.02 | 2.863(3) | 162.6 |
| O6 | H6B | O1<sup>1</sup> | 0.87 | 2.34 | 3.142(3) | 152.8 |
| N2 | H2 | O6<sup>2</sup> | 0.88 | 2.17 | 2.958(4) | 148.5 |
| N6 | H6 | O5<sup>3</sup> | 0.88 | 2.21 | 2.908(4) | 136.3 |
| O7 | H7C | O3 | 0.87 | 1.88 | 2.7479(14) | 171.8 |
| O7 | H7D | O3<sup>4</sup> | 0.87 | 1.95 | 2.7479(14) | 152.8 |

<sup>1</sup>+X,1+Y,+Z; <sup>2</sup>1/2-X,-1/2+Y,1-Z; <sup>3</sup>1-X,+Y,2-Z; <sup>4</sup>1-X,+Y,1-Z

**Table 7.**
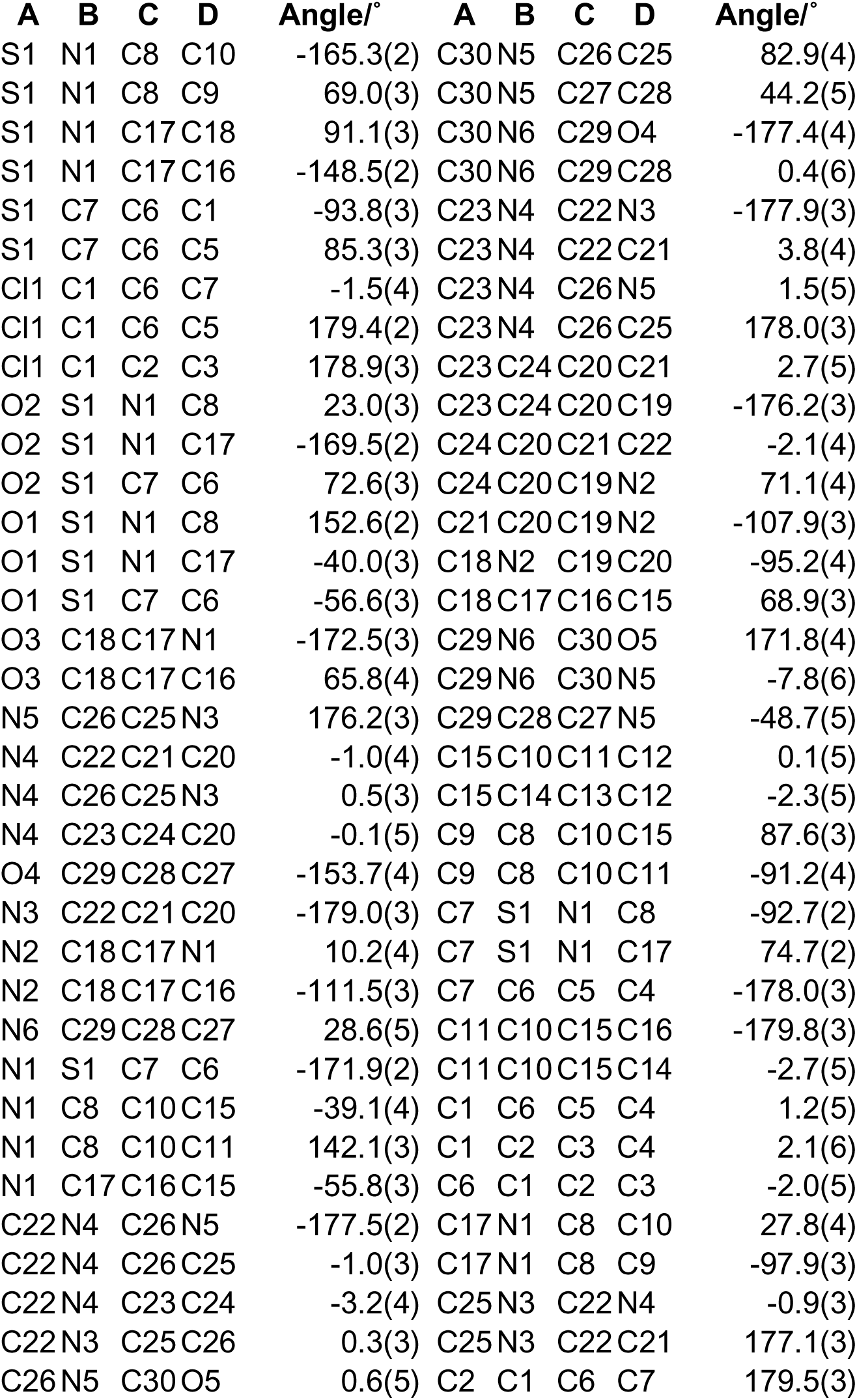
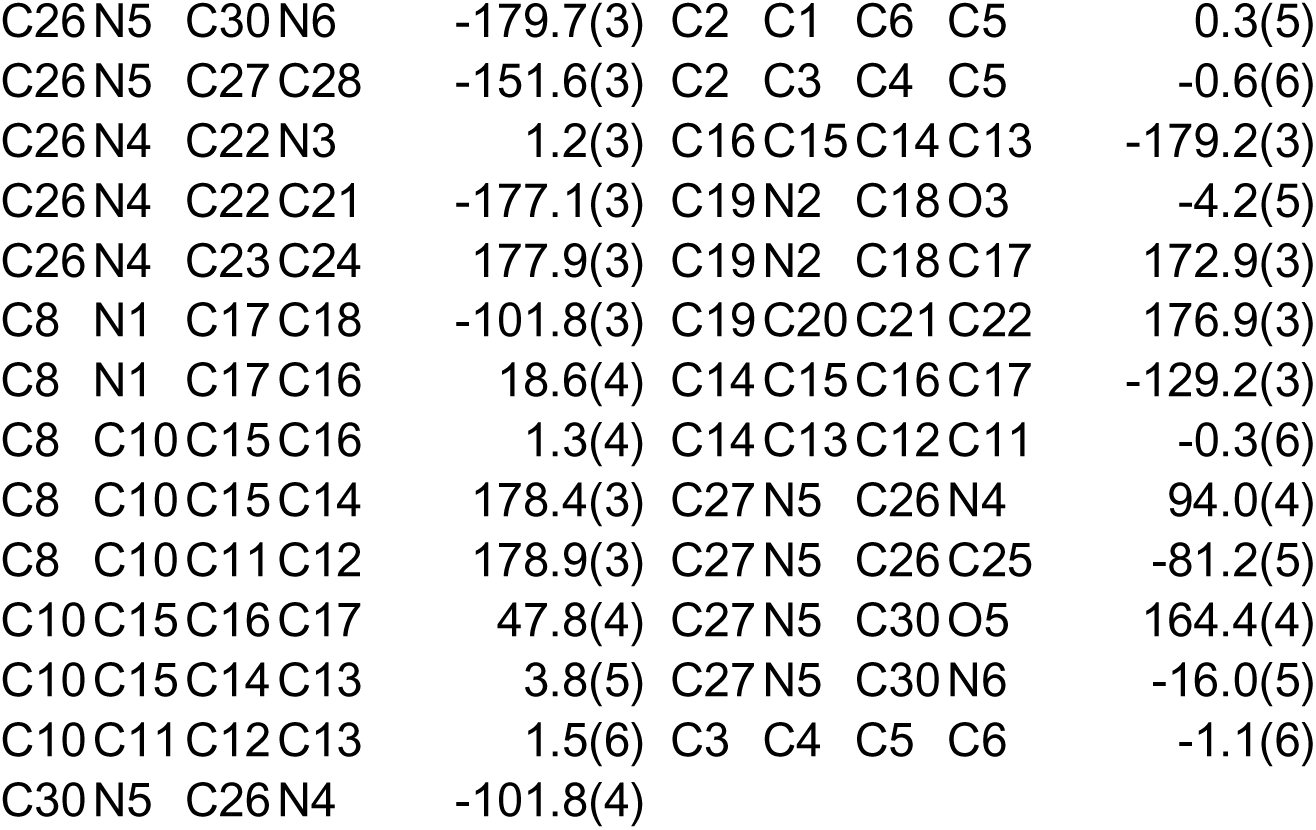
Torsion Angles for EW40755-419-P3B_auto.

**Table 8.**
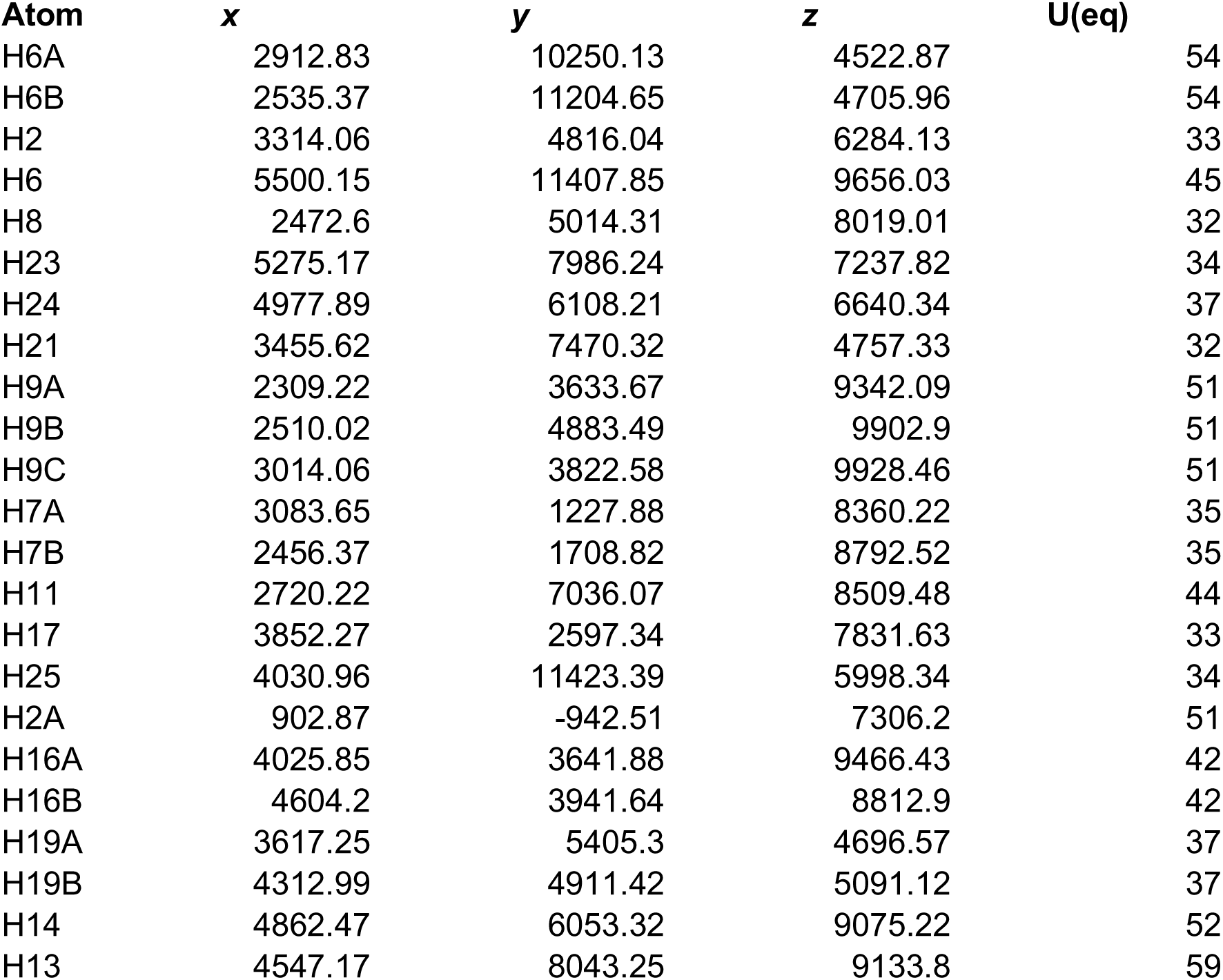
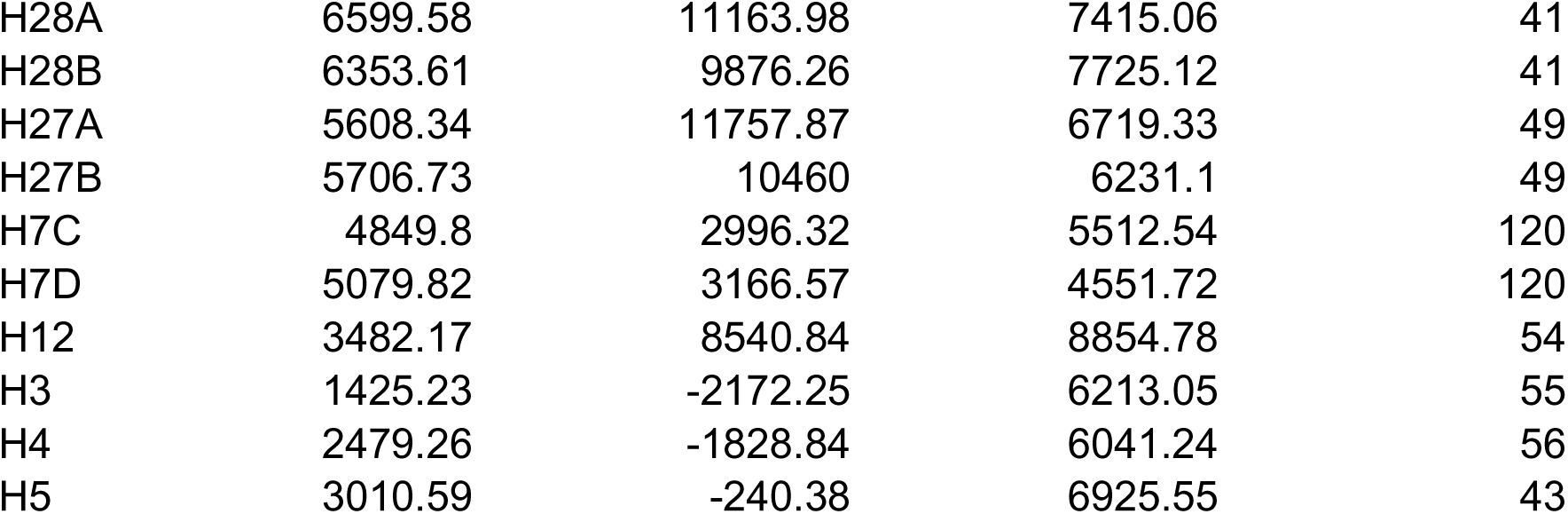
Hydrogen Atom Coordinates (Å×10^4^) and Isotropic Displacement Parameters (Å^2^×10^3^) for EW40755-419-P3B_auto.

| Atom | x | y | z | U(eq) |
| --- | --- | --- | --- | --- |
| H6A | 2912.83 | 10250.13 | 4522.87 | 54 |
| H6B | 2535.37 | 11204.65 | 4705.96 | 54 |
| H2 | 3314.06 | 4816.04 | 6284.13 | 33 |
| H6 | 5500.15 | 11407.85 | 9656.03 | 45 |
| H8 | 2472.6 | 5014.31 | 8019.01 | 32 |
| H23 | 5275.17 | 7986.24 | 7237.82 | 34 |
| H24 | 4977.89 | 6108.21 | 6640.34 | 37 |
| H21 | 3455.62 | 7470.32 | 4757.33 | 32 |
| H9A | 2309.22 | 3633.67 | 9342.09 | 51 |
| H9B | 2510.02 | 4883.49 | 9902.9 | 51 |
| H9C | 3014.06 | 3822.58 | 9928.46 | 51 |
| H7A | 3083.65 | 1227.88 | 8360.22 | 35 |
| H7B | 2456.37 | 1708.82 | 8792.52 | 35 |
| H11 | 2720.22 | 7036.07 | 8509.48 | 44 |
| H17 | 3852.27 | 2597.34 | 7831.63 | 33 |
| H25 | 4030.96 | 11423.39 | 5998.34 | 34 |
| H2A | 902.87 | -942.51 | 7306.2 | 51 |
| H16A | 4025.85 | 3641.88 | 9466.43 | 42 |
| H16B | 4604.2 | 3941.64 | 8812.9 | 42 |
| H19A | 3617.25 | 5405.3 | 4696.57 | 37 |
| H19B | 4312.99 | 4911.42 | 5091.12 | 37 |
| H14 | 4862.47 | 6053.32 | 9075.22 | 52 |
| H13 | 4547.17 | 8043.25 | 9133.8 | 59 |
| H28A | 6599.58 | 11163.98 | 7415.06 | 41 |
| H28B | 6353.61 | 9876.26 | 7725.12 | 41 |
| H27A | 5608.34 | 11757.87 | 6719.33 | 49 |
| H27B | 5706.73 | 10460 | 6231.1 | 49 |
| H7C | 4849.8 | 2996.32 | 5512.54 | 120 |
| H7D | 5079.82 | 3166.57 | 4551.72 | 120 |
| H12 | 3482.17 | 8540.84 | 8854.78 | 54 |
| H3 | 1425.23 | -2172.25 | 6213.05 | 55 |
| H4 | 2479.26 | -1828.84 | 6041.24 | 56 |
| H5 | 3010.59 | -240.38 | 6925.55 | 43 |

**Table 9.** Atomic Occupancy for EW40755-419-P3B_auto.

| Atom | Occupancy | Atom | Occupancy | Atom | Occupancy |
| --- | --- | --- | --- | --- | --- |
| H7C | 0.5 | H7D | 0.5 |  |  |

